# Plastid phylogenomics of the Gynoxoid group (Senecioneae, Asteraceae) highlights the importance of motif-based sequence alignment amid low genetic distances

**DOI:** 10.1101/2021.04.23.441144

**Authors:** Belen Escobari, Thomas Borsch, Taylor S. Quedensley, Michael Gruenstaeudl

## Abstract

**PREMISE:** The genus *Gynoxys* and relatives form a species-rich lineage of Andean shrubs and trees with low genetic distances within the sunflower subtribe Tussilaginineae. Previous molecular phylogenetic investigations of the Tussilaginineae have included few, if any, representatives of this Gynoxoid group or reconstructed ambiguous patterns of relationships for it.

**METHODS:** We sequenced complete plastid genomes of 21 species of the Gynoxoid group and related Tussilaginineae and conducted detailed comparisons of the phylogenetic relationships supported by the gene, intron, and intergenic spacer partitions of these genomes. We also evaluated the impact of manual, motif-based adjustments of automatic DNA sequence alignments on phylogenetic tree inference.

**RESULTS:** Our results indicate that the inclusion of all plastid genome partitions is needed to infer fully resolved phylogenetic trees of the Gynoxoid group. Whole plastome-based tree inference suggests that the genera *Gynoxys* and *Nordenstamia* are polyphyletic and form the core clade of the Gynoxoid group. This clade is sister to a clade of *Aequatorium* and *Paragynoxys* and also includes some but not all representatives of *Paracalia*.

**CONCLUSIONS:** The concatenation and combined analysis of all plastid genome partitions and the construction of manually curated, motif-based DNA sequence alignments are found to be instrumental in the recovery of strongly supported relationships of the Gynoxoid group. We demonstrate that the correct assessment of homology in genome-level plastid sequence datasets is crucial for subsequent phylogeny reconstruction and that the manual post-processing of multiple sequence alignments improves the reliability of such reconstructions amid low genetic distances between taxa.

## INTRODUCTION

The five genera of the sunflower family *Gynoxys* Cass., *Aequatorium* B. Nord., *Nordenstamia* Lundin, *Paracalia* Cuatrec., and *Paragynoxys* (Cuatrec.) Cuatrec. form a lineage of high-elevation shrubs and trees that are distributed in the Andean region of northwestern South America (Nordenstam et al., 2009). Under the current taxonomic circumscription, the group comprises 150–170 species (Cuatrecasas, 1951; Nordenstam, 2007; Beck and Garcia, 2014), with *Gynoxys* containing the most species. This clade is commonly known as the ‘Gynoxoid group’, and novel species continue to be described (e.g., Beltran and Campos de la Cruz, 2009). Biogeographically, the group ranges from northern Venezuela to northern Argentina and represents an example of high-elevation Andean plant diversification (Vision and Dillon, 1996; Lundin, 2006): most of its species are characteristic elements of montane forests or solitary shrubs and trees in the paramo (Tinoco et al., 2013), while only a few exhibit a scandent growth form and inhabit lower elevation montane forests (Beck and Garcia 2014; Figure 1). Most of the species of the Gynoxoid group are restricted to relatively small distribution ranges and occur in habitats threatened by anthropogenic land use and climate change, thus making this a group of conservation concern (Beltran et al., 2006; Morillo and Briceno, 2000; Hind, 2007). Based on the most recent phylogenetic investigations of the Senecioneae, the Gynoxoid group is part of the subtribe Tussilagininae of the tribe Senecioneae (Pelser et al., 2007, 2010).

**Figure 1.**
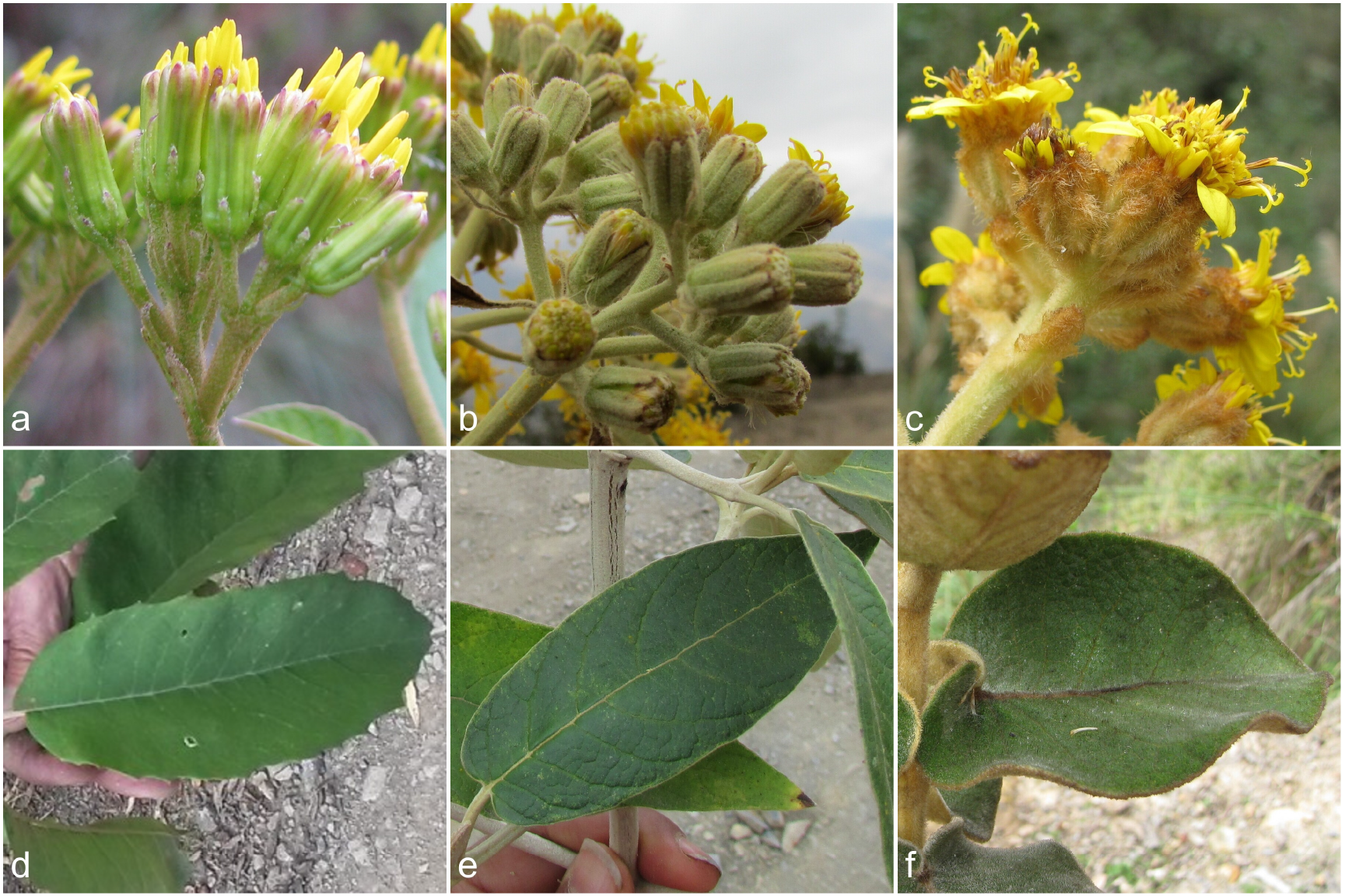
Morphological variability among three species of the Gynoxioid group. Displayed are the leaves and capitulescences of *Nordenstamia repanda* (a and d), which represents a characteristic element of upper cloud forests (bosque yungueño de Ceja de monte), *Gynoxys asterotricha* (b and e), which represents a characteristic element of lower cloud forests, and *Gynoxys tomentosissima* (c and f), which represents a characteristic element of low montane forests.

The phylogenetic relationships and species limits within the Gynoxoid group are poorly understood, as few molecular phylogenetic investigations have included taxa of this group or focused on aspects other than their relationships. Kadereit and Jeffrey (1996) conducted a study on the phylogeny of the Senecioneae using chloroplast restriction site data and included one species of *Gynoxys* in their dataset. Their results indicated that *Gynoxys* was most closely related to *Tussilago* L., *Roldana* La Llave, and *Brachyglottis* J.R. Forst. & G. Forst. Similarly, Pelser et al. (2007) aimed to infer relationships within the Senecioneae using the internal transcribed spacer (ITS) region and recovered a clade comprising the genera *Aequatorium, Gynoxys, Nordenstamia*, and *Paragynoxys*, but with mixed levels of statistical support. Their results indicated that the current taxonomic circumscriptions within the Gynoxoid group were not fully substantiated by DNA sequence data, as *Nordenstamia* was found nested within a paraphyletic *Gynoxys*. Pelser et al. (2010) extended their previous taxon sampling of the Gynoxoid group by including one sample of *Paracalia* and recovered the group as monophyletic. Recently, Quedensley et al. (2018) included taxa of *Aequatorium* and *Gynoxys* in a study of North and Central American representatives of the Tussilaginineae and recovered a monophyletic Gynoxoid group with strong statistical support; however, no insight into the intergeneric relationships of the group was investigated. In summary, the intergeneric relationships of the Gynoxoid group have remained largely unresolved, and the delimitation of its genera is mostly unsubstantiated by phylogenetic methods.

Plastid phylogenomic studies have been shown to be efficient in resolving the phylogenetic relationships of species groups in the Asteraceae. Vargas et al. (2017) investigated the relationships of *Diplostephium* Kunth and related genera that exemplified low levels of molecular variability (Vargas and Madrinan, 2012). The authors sequenced complete plastid genomes of 14 different genera (91 samples) and inferred a highly resolved phylogeny. Similarly, Pouchon et al. (2018) sequenced plastid genomes in an analysis of the phylogenetic relationships of *Espeletia* Mutis ex Bonpl. and relatives, which previous studies could not resolve due to an insufficient number of informative DNA sequence characters available. Specifically, the authors sequenced complete plastid genomes of eight different genera (41 species) and recovered well-supported clades. Zhang et al. (2019) sequenced plastid genomes to establish a robust phylogenetic framework for the species-rich genus *Saussurea* DC., for which previous work had generated conflicting infrageneric classifications. By analyzing complete plastid genomes of 136 species of *Saussurea*, the authors found that approximately 2,000 parsimony informative sites were needed to produce a resolved and well-supported phylogeny. More recently, Knope et al. (2020) sequenced plastid genomes to reconstruct the phylogeny of Hawaiian endemics of the genus *Bidens* L. in light of a decade-long effort to clarify the evolutionary history of this rapidly radiating lineage and were able to generate a highly supported phylogeny. Evidently, the use of plastid genomes for phylogenetic inference can be instrumental in clarifying the relationships of plant groups that exhibit low molecular variability. This utility is not restricted to Asteraceae but has been demonstrated in numerous lineages of flowering plants (e.g., Givnish et al., 2018; Yao et al., 2019).

The process of sequencing and comparing plastid genomes for the reconstruction of phylogenetic relationships is often perceived as simple, but several studies have indicated that the evolution and structure of plastid genomes and, by extension, their application in phylogenetic inference, is complex. Based on observations that the majority of plastid genomes display strong structural conservation (Mower and Vickrey, 2018), are uniparentally (mostly maternally) inherited (Greiner et al., 2015), and do not experience biparental recombination (Marechal and Brisson, 2010), many investigations have operated under the assumption of a congruent phylogenetic signal across the entire plastid genome. However, several studies have cast doubt on the validity of this assumption and instead highlighted the presence of a discordant phylogenetic signal across different regions of the plastid genome. Investigations on the more variable regions of plastid genomes, for example, found mosaic-like patterns of molecular evolution (Borsch and Quandt, 2009), a hierarchical structure of phylogenetic signal (Müller et al., 2006; Barniske et al., 2012), and lineage-specific lengths and positions of these regions (Korotkova et al., 2014). Recent phylogenomic investigations corroborated these reports by identifying considerable phylogenetic incongruence across different regions of the plastid genome, which may result in inefficient or even incorrect phylogenetic inferences if the entire genome is analyzed under the same model parameters. Goncalves et al. (2019), for example, identified significant incongruence among the gene and species trees of different plastome regions in a phylogenomic study on rosids. The authors reported that the concatenation of all plastid coding regions produced highly supported phylogenies that were nonetheless incongruent to individual plastid gene trees. Similarly, Gruenstaeudl (2019) detected phylogenetic incongruence across different loci of the plastid genome in a phylogenomic investigation of water lilies and relatives. Walker et al. (2019) reported gene tree conflict among various plastid genes based on a broad sampling of angiosperm plastid genomes and noted numerous strongly supported but conflicting nodes between different gene trees. Similar observations were made in plastid phylogenomic analyses of Fabaceae (Zhang et al., 2020) and Bignoniaceae (Thode et al., 2020). Furthermore, Koehler et al. (2020) identified and ranked plastid genome regions by phylogenetic informativeness and found topological incongruence between the phylogenetic trees inferred from complete plastid genome sequences and those inferred from the five and ten most variable plastid regions only. Based on these and similar studies, Goncalves et al. (2020) cautioned that ‘one or a few genes that have high phylogenetic signal may bias the inference’ (Goncalves et al., 2020, p. 4) and, thus, recommended the continued exploration of more phylogenetically information among different regions of the plastid genome.

The positional homology among the nucleotides of a multiple sequence alignment (MSA) represents an essential aspect of phylogenetic tree inference and has not received sufficient attention in many phylogenomic investigations. Early plastid phylogenomic studies primarily employed the coding regions of the genomes for phylogenetic reconstruction (e.g., Leebens-Mack et al., 2005; Moore et al., 2010), which are largely conserved in their length and sequence and, thus, require relatively little adjustment upon standard MSA. Phylogenetic investigations that employ non-coding plastid DNA, by contrast, routinely inspect software-generated MSAs and adjust them according to the criterion of explicit sequence motifs to ensure correct positional homology (Kelchner, 2000; Loehne and Borsch, 2005; Morrison, 2006). In phylogenomic analyses, such adjustments are sometimes dismissed as impractical due to the large amounts of sequence data involved (Wu et al., 2012), but numerous investigations have demonstrated the impact of alignment errors on phylogenetic inference (reviewed by Wong et al., 2008). To reduce this impact while simultaneously avoiding time-expensive evaluations of the MSAs, many studies tend to automatically exclude nucleotide positions in software-generated MSAs that are deemed unreliable (e.g., Bellot et al., 2020). In practice, such procedures may go as far as excluding alignment positions that exhibit a gap in only one of the aligned sequences, which substantially reduces the proportion of genome sequence employed for phylogenetic reconstruction (e.g., Gernandt et al., 2018). Along with the reduction of potential informativeness, such exclusions do not guarantee correct positional homology in the remaining alignment, as demonstrated for cases of small genomic inversions (Ochoterena, 2008). Given the larger share of non-coding compared to coding DNA in plastid genomes as well as the higher frequency of substitutions and microstructural mutations in non-coding DNA, manual adjustments of software-derived MSAs may even have a considerable impact on plastid phylogenomic reconstruction. In fact, in species groups with low genetic distances, a large proportion of potentially informative sequence characters will likely be encoded in the non-coding regions of the plastid genome.

The present investigation has two major goals: (1) to infer the phylogenetic relationships within the Gynoxoid group of the Tussilaginineae using complete plastid genomes to account for the low genetic distances expected within the target group, and (2) to assess the phylogenetic signal of different sections of the plastid genome, particularly with regard to motif-based adjustments of software-generated MSAs. Specifically, we contrast the results of phylogenetic tree inference based on different coding regions, introns, and intergenic spacers of the plastid genome before and after the manual adjustment of the MSAs. To achieve these goals, we sequenced and annotated complete plastid genomes of 17 species of the Gynoxoid group as well as four species of closely related members of the Tussilaginineae to form a dataset that reflects the genetic distances within the Gynoxoid group and to other members of the Tussilaginineae. Based on this dataset, we ask four specific questions: (i) Does phylogenetic analysis of complete plastid genomes yield well-resolved and well-supported phylogenetic trees for the Gynoxoid group? (ii) Do different partitions of the plastid genome (i.e., coding sequences, intergenic spacers, and introns) support different phylogenetic hypotheses? (iii) Does the manual adjustment of MSAs have a measurable impact on phylogenetic reconstruction? (iv) Are the results of the plastome-based reconstructions congruent with the current generic classification of the Gynoxoid group?

## MATERIALS AND METHODS

### Taxon sampling and DNA extraction

A total of 21 samples of different species of the Tussilaginineae were collected for DNA extraction and subsequent plastid genome sequencing (Table 1). Of these, 17 samples represent genera of the Gynoxoid group (i.e., *Aequatorium, Gynoxys, Nordenstamia, Paracalia*, and *Paragynoxys*), while four represent other genera of the Tussilaginineae that form the sister clade to the Gynoxoid group (i.e., *Arnoglossum* Raf., *Roldana* La Llave, and *Telanthophora* H. Rob. & Brettell; see Quedensley et al. 2018 for details). Particular emphasis in our taxon sampling was placed on the inclusion of (i) more than one species per genus, where possible, to approximate the genetic variability within each genus, (ii) multiple species of *Gynoxys* to represent its species diversity across the Andes and to accommodate the previous results that *Gynoxys* may be non-monophyletic, and (iii) the type species of the genera *Gynoxys, Nordenstamia*, and *Paracalia* to enable comparisons of our reconstructions with the current taxonomic classification of the Gynoxoid group. As an outgroup, we included the previously published plastid genome of *Ligularia fischeri* (Ledeb.) Turcz. (GenBank accession NC 039352; Chen et al., 2018). The taxon names and generic concepts applied follow Nordenstam (2007). Herbarium vouchers of all newly sequenced samples of the Gynoxoid group were deposited in B, with duplicates in LPB, USM, HSP, or HUT.

**Table 1.** Species name, collection location and number, herbarium voucher information, and GenBank accession number for each plastid genome sequenced in this investigation. n.a. = not applicable.

| Species name | Location information | Collection number | Herbarium voucher | DNA Bank code | GenBank |
| --- | --- | --- | --- | --- | --- |
| *Aequatorium jamesonii* | Ecuador: Napo, Parque Nacional Llanganates | Vargas H. et al. 2594 | MO-2940002 (MO) | DB38638 | MT528247 |
| *Arnoglossum atriplicifolium* | USA: Spring Lake Park, Omaha, Nebraska | Quedensley T.S. s.n. | TSQ2011cp001 (TEX) | n.a. | MK170176 |
| *Gynoxys asterotricha* | Bolivia: Parque Nacional Madidi, Laji Sorapata | Escobari B. et al. 36 | B-100720926 (LPB,B) | DB27411 | MK044798 |
| *Gynoxys baccharoides* | Ecuador: Azuay, Parque Nacional Cajas | Jorgensen P. et al. 1616 | MO-1879904 (AAU,MO) | DB38633 | MT528244 |
| *Gynoxys ignaciana* | Ecuador: Loja, Guararas | Emperaire L. 1318 | P-03833291 (P) | DB38767 | MT528252 |
| *Gynoxys longifolia* | Peru: Arequipa, Cailloma | Beck S. 26384 | LPB0002593 (LPB,MO,US) | DB38565 | MT528245 |
| *Gynoxys mandonii* | Bolivia: La Paz, Nor Yungas, Unduavi | Cayola L. 5570 | MO-3151322 (LPB,MO) | DB27416 | MK056106 |
| *Gynoxys megacephala* | Bolivia: Nor Yungas, Ecovia | Escobari B. et al. 46 | LPB0002594 (LPB) | DB27426 | MN328892 |
| *Gynoxys sp. SEN301* | Peru: Amazonas, Leymebamba | Escobari B. et al. 593 | B-101098545 (USM,HUT,B) | DB38744 | MN328891 |
| *Gynoxys tomentosissima* | Peru: Amazonas, Leymebamba | Escobari B. et al. 590 | B-101098697 (USM,HUT,B) | DB38742 | MN328890 |
| *Gynoxys violacea* | Venezuela: Merida | Walter E. 166 | B-101139853 (B) | DB38780 | MT528243 |
| *Nordenstamia cajamarcensis* | Peru: Pasco, Oxapampa | Castillo G. 460 | MO-2940516 (F,MO,USM) | DB38626 | MT528248 |
| *Nordenstamia kingii* | Bolivia: Cochabamba, Monte Puncu | Solomon J. 18067 | LPB0002595 (LPB,MO) | DB38764 | MT528249 |
| *Nordenstamia repanda* | Bolivia: Parque Nacional Madidi, Laji Sorapata | Cayola L. 5585 | LPB0002597 (LPB,MO) | DB27409 | MK086040 |
| *Paracalia jungioides* | Peru: Ancash, Huaraz | Diaz C. 1975 | MO-3030471 (LPB,MO) | DB43895 | MT528251 |
| *Paracalia pentamera* | Bolivia: Sud Yungas, Chulumani | Gallegos S. 3850 | LPB0002596 (LPB) | DB40913 | MT942595 |
| *Paragynoxys martingrantii* | Colombia: Cesar | Gentry A. 79156 | MO-1962013 (MO) | DB43896 | MT528250 |
| *Paragynoxys venezuelae* | Venezuela: Merida, Paramo de Las Coloradas | Cuatrecasas J. et al. 28996 | MA-01-00896838 (MA,S) | DB43902 | MT528246 |
| *Roldana aschenborniana* | Mexico: Oaxaca (cult. ex-situ in California) | Quedensley T.S. s.n. | TSQ2011cp002 (TEX) | n.a. | MK170177 |
| *Roldana barba-johannis* | Mexico: Oaxaca (cult. ex-situ in California) | Quedensley T.S. s.n. | TSQ2011cp003 (TEX) | n.a. | MK170178 |
| *Telanthophora grandifolia* | Mexico: Oaxaca (cult. ex-situ in California) | Quedensley T.S. s.n. | TSQ2011cp004 (TEX) | n.a. | MK170179 |

Extractions of total genomic DNA were generated from silica gel-dried leaf material using the NucleoSpin Plant II kit (Macherey-Nagel, Dueren, Germany) or from herbarium specimens using the CTAB DNA isolation method as modified by Borsch et al. (2003). Following DNA extraction, aliquots of the isolates were deposited in the DNA bank of the Botanic Garden and Botanical Museum Berlin (BGBM; Table 1). Unless DNA isolates were fragmented due to age, samples were sheared to an average fragment size of 600 bp using a Covaris S220 sonicator (Covaris, Woburn, MA, USA). Upon shearing, all fragments of a length between 400 bp and 900 bp were selected and maintained by applying the BluePippin protocol (Sage Science, Beverly, CA, USA). The final concentration of DNA samples was measured using Qubit 2.0 Fluorometer dsDNA BR Assay kits (Life Technologies-Thermo Fisher Scientific, Saint Aubin, France) and the final fragment size distribution using an Agilent 2100 Bioanalyzer (Agilent Technologies, Santa Clara, CA, USA). All steps upon DNA extraction were conducted according to manufacturer specifications.

### Genomic library preparation and DNA sequencing

Plastid genome sequencing was conducted via a genome skimming approach upon preparation of genomic libraries. For each DNA sample, a barcoded genomic library was constructed using the TruSeq DNA library preparation kit (Illumina, San Diego, CA, USA). Standard indexing adapters were ligated to the fragment ends to generate single-index libraries. Libraries were validated via qPCR on a Mastercycler ep realplex (Eppendorf AG, Hamburg, Germany) using the KAPA library quantification kit (KAPA Biosystems, Wilmington, MA, USA). Following qPCR, indexed DNA libraries were normalized and pooled in equal volumes. Pooled libraries were sequenced as paired-end reads either on an Illumina MiSeq or an Illumina HiSeq X platform (Illumina, San Diego, CA, USA). Sequencing was performed in-house at the Berlin Center for Genomics in Biodiversity Research (Berlin, Germany) or the Genome Sequencing and Analysis Facility of the University of Texas at Austin, or outsourced to Macrogen Inc. (Seoul, Republic of Korea).

### Genome assembly and annotation

Upon DNA sequencing, raw sequence reads were filtered for quality and successful pairing using scripts 1 and 2 of the pipeline of Gruenstaeudl et al. (2018), followed by a mapping of the quality-filtered reads to the plastid genome of *Jacobaea vulgaris* Gaertn. (accession NC 015543; Doorduin et al., 2011) to extract plastome reads using bowtie2 v.2.3.4 (Langmead and Salzberg, 2012). Contigs were assembled *de novo* with either IOGA v.20160908 (Bakker et al., 2015) or NOVOPlasty v.2.7.2 (Dierckxsens et al., 2017) based on the subset of reads that mapped against the reference genome, using a range of different kmer values to optimize contig length (kmer = 33–97, in increments of 4). Unless already circular, final contigs were circularized manually with Geneious v.11.1.4 (Kearse et al., 2012), using the plastid genome of *Jacobaea vulgaris* as a reference for contig position and orientation. A circular, quadripartite structure of the plastid genome and equality of its inverted repeat (IR) regions were confirmed by blasting each assembly against itself using script 4 of the pipeline of Gruenstaeudl et al. (2018). Sequence ambiguities in the final assembly, if present, were resolved by mapping the quality-filtered reads against the circularized assembly using bowtie2. Final assemblies were annotated via the annotation server DOGMA (Wyman et al., 2004), followed by a manual inspection and, where necessary, correction of the annotations in Geneious. Specifically, sequence annotations were corrected regarding the presence of start and stop codons, the absence of internal stop codons, and their length as a multiple of three for each coding region. Upon annotation, all newly sequenced plastid genomes were deposited to GenBank; their accession numbers are listed in Table 1. Plastome maps were drawn with OGDRAW v.1.3.1 (Greiner et al., 2019).

### Data partitioning, sequence alignment, and alignment adjustments

The coding and non-coding regions of the plastid genomes were extracted and aligned using a four-step procedure. First, one of the IRs was removed from each genome to avoid redundancy among the extracted loci. Second, all coding and non-coding regions (except tRNAs and rRNAs) were excised bioinformatically from each genome using script 9 of the pipeline of Gruenstaeudl et al. (2018) and then grouped by region name. Third, all sequences of the same region were aligned into preliminary MSAs using MAFFT v.7.394 (Katoh and Standley, 2013) under the default settings of the software. Specifically, MSAs of 81 coding regions, 20 introns, and 111 intergenic spacers, each consisting of the sequences of 22 taxa, were constructed. Fourth, these preliminary MSAs were evaluated by eye and, where necessary, adjusted manually to improve positional homology across nucleotides using PhyDE v.0.9971 (Müller et al., 2010). The adjustments followed the rules of Loehne and Borsch (2005) and also included the masking of mutational hotspots for those sections of the MSAs where correct positional homology could not be established. For example, mono- or dinucleotide microsatellites (e.g., poly-A repeats) were masked by excluding them from subsequent reconstructions. Small sequence inversions were masked by a manual re-inversion of the sequence motifs, followed by a re-alignment to the other sequences and the recording of each inversion as a single-step event to be later added to the indel matrix. Examples of the masking of microsatellites and sequence inversions are illustrated in Figure 2; a summary of the positions and lengths of masked sequence regions within the MSAs is given in Table 2. Following these alignment adjustments, all MSAs were assessed for length and sequence variability and excluded from the dataset if any of the following criteria were met: lack of sequence variability, length less than 10 bp, more ambiguous than variable nucleotides if length less than 50 bp, and vicinity to trans-spliced genes. For example, the MSAs of the intergenic spacers *ndhH*–*ndhA, rpoB*–*rpoC1, ndhK*–*psbG*, and *psbF*–*psbE* were excluded because they were only one, five, seven, and nine bp long, respectively. Similarly, the spacer *psbT*–*psbN* was excluded due to the combination of short alignment length and more ambiguous than variable nucleotides. All tRNA and rRNA genes were excluded due to minimal, if any, sequence variability. The intergenic spacers adjacent to *rps12* (i.e., *rpl20*–*rps12, rps12*–*clpP*, and *rps7*–*rps12*) were excluded due to the trans-splicing nature of this gene. All other MSAs were saved as NEXUS files for further processing. Insertions and deletions in each MSA were coded as binary characters using the simple indel coding (SIC) scheme of Simmons and Ochoterena (2000) as implemented in SeqState v.1.4.1 (Müller, 2005). The complete sets of MSAs representing 81 different coding regions, 20 different introns, and 103 different intergenic spacers were grouped by marker class (hereafter ‘plastid partitions’; i.e., coding regions, introns, and intergenic spacers) and then concatenated with and without indel codes. Specifically, the MSAs were concatenated within each set as well as across the three sets, both with and without the presence of indel codes, generating a total of eight different matrices. All matrices were deposited to Zenodo under DOI 10.5281/zenodo.4428211 and employed for phylogenetic tree reconstruction. For brevity, coding regions are abbreviated with the term ‘CDS’ in our analyses, introns with ‘INT’, and intergenic spacers with ‘IGS’. Similarly, the concatenation of all MSAs of the coding regions is abbreviated with ‘81 CDS concat’, the concatenation of all MSAs of the introns as ‘20 INT concat’, and the concatenation of all MSAs of the intergenic spacers as ‘103 IGS concat’.

**Table 2.**
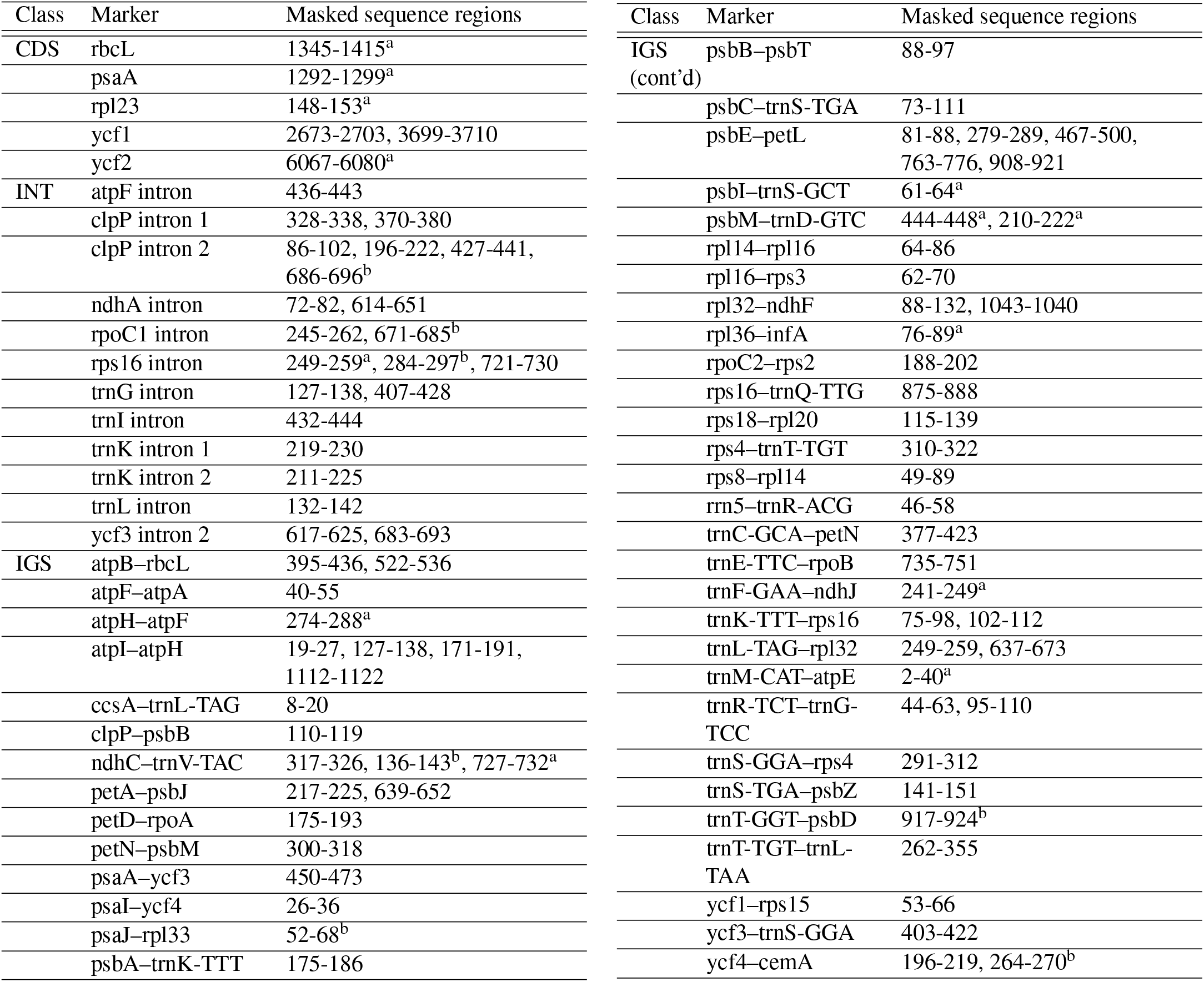
Summary of the position and length of the microsatellites and small sequence inversions in the MSAs that were masked during alignment adjustment. By default, the listed regions represent microsatellites with a DNA motif comprising an A and/or T, unless marked otherwise: ^a^ = sequence inversions with length *<*50 bp; ^b^ = microsatellites with a DNA motif comprising a G and/or C.

**Figure 2.**
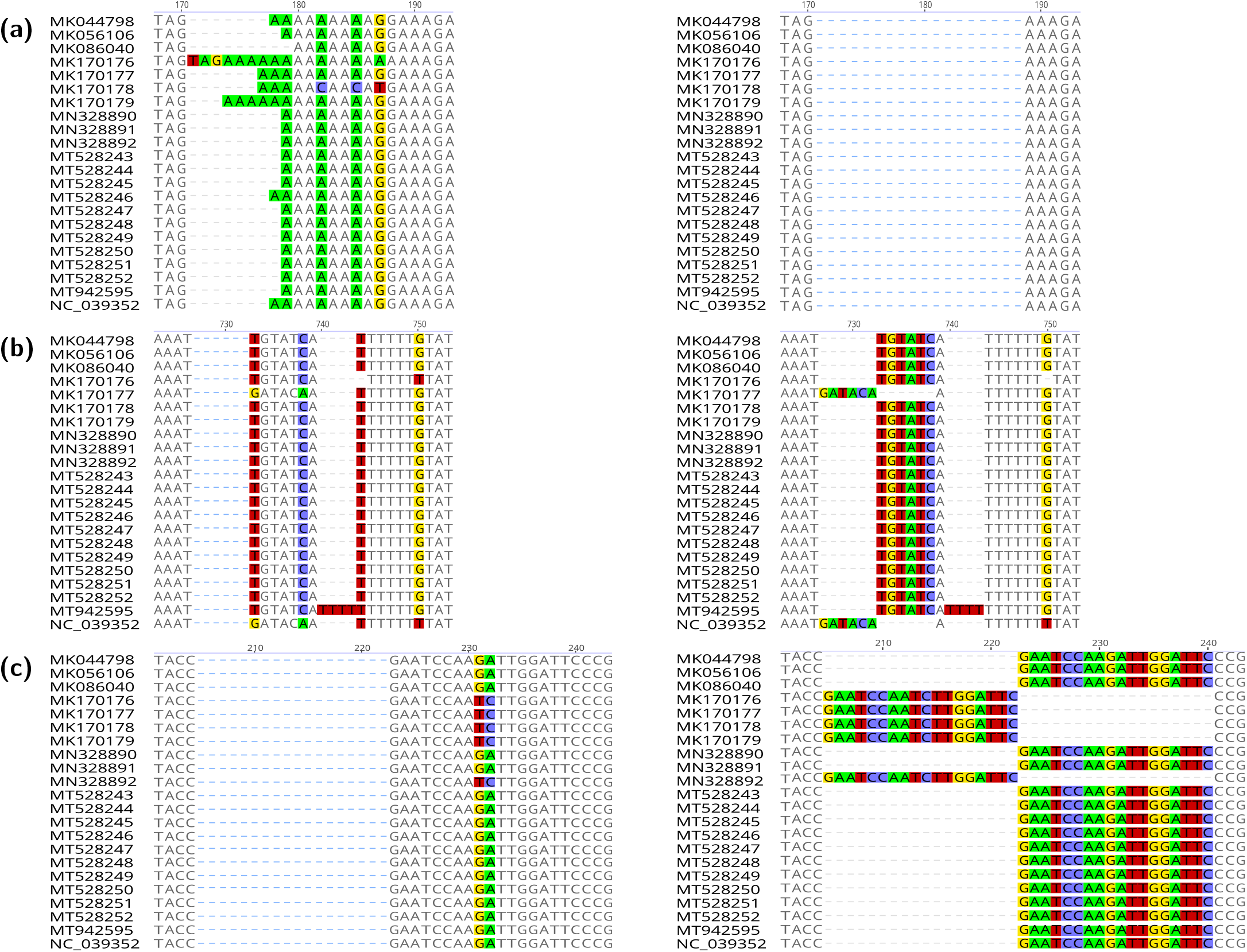
Illustration of the manual adjustment of sequence alignments as exemplified by three different intergenic spacers of the plastid genome. For each spacer, the software-generated MSA is displayed on the left, the MSA after manual adjustment on the right. Polymorphic nucleotides are highlighted in color. Row (a) illustrates the adjustment of the MSA of spacer *atpI*–*atpH*, which contains a poly-A region with internal nucleotide polymorphism; the poly-A region was removed due to uncertain positional homology. Row (b) illustrates the adjustment of the MSA of spacer *ndhC*–*trnV*, which contains shared sequence inversions of a length of 6 bp; the inversions were manually inverted to avoid incorrect positional homology. Row (c) illustrates the adjustment of the MSA of spacer *psbM*–*trnD*, which contains shared sequence inversions of a length of 18 bp; the inversions were manually inverted to avoid incorrect positional homology. Gaps represented by blue dashes were added to the visualization of the alignments for easier comparisons.

### Alignment metrics and sequence variability

To assess the reliability of the MSAs and the impact of manual alignment adjustment on them, we calculated a total of eight different alignment metrics for each MSA before and after alignment adjustment. The inferred metrics were: (i) alignment length, (ii) GC content, (iii) fraction of polymorphic sites, (iv) fraction of parsimony-informative sites, (v-vii) the three homoplasy indices ‘consistency index’ (Kluge and Farris, 1969), ‘rescaled consistency index’, and ‘retention index’ (both Farris, 1989), and (viii) the largest uncorrected p-distance between all sequences as defined in equation 3.1 of Nei and Kumar (2000). Each metric was calculated in R (Team, 2019) using the R packages ape v.5.2 (Paradis and Schliep, 2018) or phangorn v.2.4.0 (Schliep, 2011). For each MSA, the three homoplasy indices were calculated on the Neighbor-joining tree of that MSA. The values of the homoplasy indices are negatively correlated with the level of homoplasy in the MSA, with high index values indicating low levels of homoplasy. Indels were taken into account during the calculation of alignment length and the fraction of polymorphic sites but were disregarded in the calculation of all other metrics in accordance with the original settings of the R functions. To quantify sequence variability across the plastid genomes under study, we calculated the nucleotide diversity index *π* (Nei and Li, 1979) *with DnaSP v*.*6*.*12*.*03 (Rozas et al., 2017)* for each MSA as well as their concatenation across all partitions using a sliding window algorithm with a step size of 200 bp and window size of 600 bp. To visualize sequence variability across the genomes, we generated variability plots using mVISTA (Frazer et al., 2004) following a global pairwise alignment of the sequences with LAGAN (Brudno et al., 2003). The IRa was stripped from each plastid genome before alignment and visualization with mVISTA; coding regions *<*10 nucleotides as well as the trans-spliced gene *rps12* were not annotated in the visualizations. The plastid genome of *Ligularia fischeri* was selected as a reference for the calculation of sequence similarity values in mVISTA.

### Phylogenetic inference

Phylogenetic tree inference was conducted under the maximum likelihood (ML) and the Bayesian inference (BI) criterion on the concatenation of all MSAs of the coding regions, the concatenation of all MSAs of the introns, the concatenation of all MSAs of the intergenic spacers, and the concatenation of all three plastid partitions. Tree inference was hereby conducted both before and after the manual adjustment of the MSAs. Tree inference under ML was performed using RAxML v.8.2.9 (Stamatakis, 2014), including the option for a thorough optimization of the best-scoring ML tree. Tree inference under BI was performed with MrBayes v.3.2.6 (Ronquist and Huelsenbeck, 2003) using four parallel Markov Chain Monte Carlo (MCMC) runs for a total of 50 million generations. Branch support under ML was calculated through 1,000 bootstrap (BS) replicates under the rapid BS algorithm (Stamatakis et al., 2008). Branch support under BI was calculated as posterior probability (PP) values. The nucleotide substitution model GTR+G+I was applied by default to model substitution rates during tree inference under both optimality criteria. In analyses under BI, the sampling of independent generations and the convergence of the Markov chains were confirmed in Tracer v.1.7 (Rambaut et al., 2018); the initial 50% of all MCMC trees were discarded as burn-in, and post-burn-in trees were summarized as a 50% majority rule consensus tree.

## RESULTS

### Genome structure and gene content

Genome structure and length as well as the number of genes per genome were found to be highly conserved across the plastid genomes of the Gynoxoid group. The genomes exhibit the standard circular and quadripartite structure, comprising one large (LSC) and one small single copy (SSC) region, separated by two identical IRs, and display minor, if any, length variability. The variability in total sequence length between the largest and the smallest plastid genome in this group was less than one kb (except for *Gynoxys tomentosissima* Cuatrec., see below; Table 3). All of the genomes of the Gynoxoid group consist of a total of 81 protein-coding regions (seven of which are duplicated in the IRs), 30 transfer RNA (tRNA) genes (seven duplicated in the IRs), and four ribosomal RNA (rRNA) genes (all duplicated in the IRs), resulting in a total of 133 functional coding regions per genome. Also, the same genes contain one or more introns across the genomes: *atpF, ndhA, ndhB, petB, petD, rpl2, rpoC1* and *rps16* contain one intron each; *clpP* and *ycf3* contain two introns each. The plastid genome of *Gynoxys tomentosissima* is slightly different than the other genomes of the Gynoxoid group: with 155,060 bp, making it the largest sequenced in this study. Compared to the other plastid genomes of the Gynoxoid group, its sequence is approximately four kb longer (Figure 3). This is caused by the expansion of its IRs into the LSC, so the genes *rpl14, rpl16, rps3, rpl22* and *rps19* are additionally duplicated in the IRs. The GC content of all plastid genomes of the Gynoxoid group was between 37.2% and 37.9% and is, thus, within the typical bandwidth of plastid GC content (Smith, 2009).

**Table 3.** Overview of length (total and by region) and GC content of the plastid genomes of the Gynoxoid group.

| Species | Total length (bp) | LSC (bp) | SSC (bp) | IR (bp) | GC% |
| --- | --- | --- | --- | --- | --- |
| *Aequatorium jamesonii* | 151,015 | 83,176 | 18,193 | 24,823 | 37.5 |
| *Arnoglossum atriplicifolium* | 150,837 | 83,036 | 18,149 | 24,826 | 37.2 |
| *Gynoxys asterotricha* | 151,022 | 83,213 | 18,195 | 24,807 | 37.5 |
| *Gynoxys baccharoides* | 151,071 | 83,215 | 18,150 | 24,853 | 37.4 |
| *Gynoxys ignaciana* | 150,979 | 83,140 | 18,191 | 24,824 | 37.5 |
| *Gynoxys longifolia* | 150,998 | 83,180 | 18,172 | 24,823 | 37.5 |
| *Gynoxys mandonii* | 151,004 | 83,169 | 18,189 | 24,823 | 37.5 |
| *Gynoxys megacephala* | 151,014 | 83,178 | 18,192 | 24,822 | 37.5 |
| *Gynoxys sp. SEN301* | 151,013 | 83,316 | 18,193 | 24,752 | 37.5 |
| *Gynoxys tomentosissima* | 155,060 | 79,946 | 18,194 | 28,460 | 37.4 |
| *Gynoxys violaceae* | 151,006 | 83,159 | 18,141 | 24,853 | 37.5 |
| *Nordenstamia cajamarcensis* | 150,907 | 83,111 | 18,122 | 24,837 | 37.5 |
| *Nordenstamia kingii* | 150,891 | 83,049 | 18,196 | 24,823 | 37.5 |
| *Nordenstamia repanda* | 151,031 | 83,192 | 18,193 | 24,823 | 37.5 |
| *Paracalia junguioides* | 151,014 | 83,183 | 18,183 | 24,824 | 37.4 |
| *Paracalia pentamera* | 151,640 | 83,095 | 18,551 | 24,997 | 37.9 |
| *Paragynoxys martingrantii* | 150,996 | 83,168 | 18,180 | 24,824 | 37.5 |
| *Paragynoxys venezuelae* | 150,927 | 83,100 | 18,181 | 24,823 | 37.5 |
| *Roldana aschenborniana* | 151,018 | 83,195 | 18,147 | 24,838 | 37.4 |
| *Roldana barba-johannis* | 151,016 | 83,230 | 18,127 | 24,829 | 37.4 |
| *Telanthophora grandifolia* | 151,093 | 83,255 | 18,148 | 24,845 | 37.4 |

**Figure 3.**
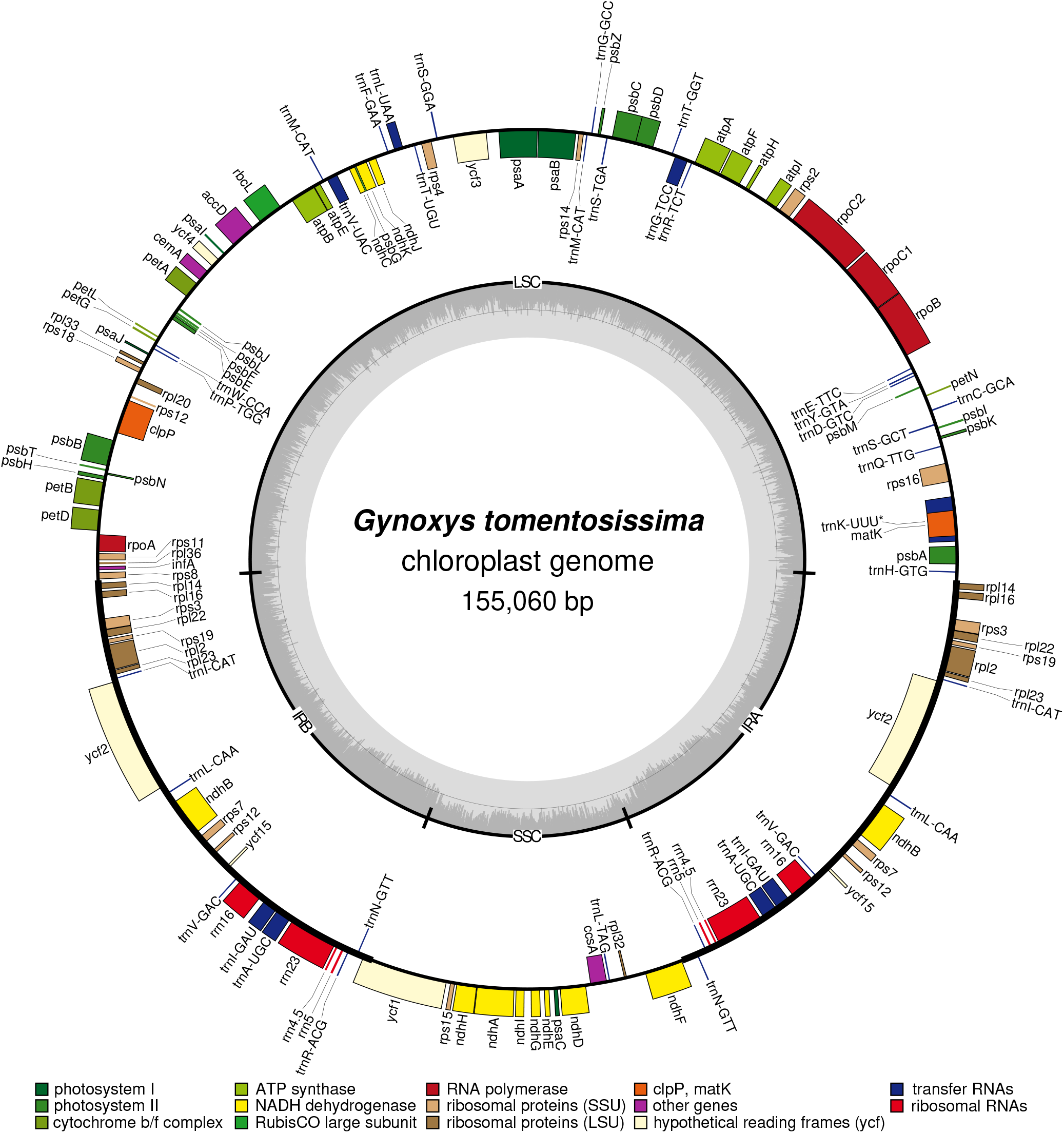
Plastome map of *Gynoxys tomentosissima*, the longest of the sequenced plastid genomes of the Gynoxoid group.

### Sequence variability across genomes

Sequence variability across the plastid genomes of the Gynoxoid group was low and located almost exclusively in the non-coding regions of the genomes, with coding regions exhibiting only occasional, if any, nucleotide polymorphism (Figure 4). Among the coding regions, only *ycf1* exhibited noticeable differences across sequences. The intergenic spacers were the most variable among the three plastid partitions (average *π* = 0.006; Table 4a), followed by the introns (*π* = 0.004) and the coding regions (*π* = 0.002). The MSAs of the intergenic spacers *psbA*–*trnK-TTT, rpl16*–*rps3*, and *rps18*–*rpl20*, for example, contained the highest nucleotide diversity (*π* = 0.20). Some sequence variability was observed in the MSAs of the introns (e.g., the actual non-coding domains of the intron of *trnK-TTT*), but most introns exhibited considerable sequence similarity across taxa. Interestingly, the visualization of sequence variability with mVISTA produced somewhat misleading results regarding their phylogenetic utility. The high level of sequence variability indicated for the intergenic spacer *trnT-GGT*–*psbD* (i.e., position 31,828–33,081 bp) was primarily the consequence of a DNA insertion shared by five sequences and only yields a single variable character for phylogenetic inference upon indel coding. In summary, the plastid genomes of the Gynoxoid group exhibit low levels of sequence variability, but considerable differences in variability do occur among genome regions.

**Table 4.** Alignment length, sequence variability, and ML tree statistics of the different datasets under study. (a) Alignment length and statistics on sequence variability, with percentages are shown in parentheses; (b) statistics on ML tree inference before and after the manual adjustment of alignments.

| (a) |  |  |  |  |  |  |  |  |  |  |
| --- | --- | --- | --- | --- | --- | --- | --- | --- | --- | --- |
| | Alignment length (bp) | | Gaps/Missing | | Polymorphic sites | | Parsimony informative sites | | Nucleotide diversity $\pi$ | |
|  | before | after | before | after | before | after | before | after | before | after |
| CDS (81) | 68,117 | 68,076 | 824<br>(1.21%) | 328<br>(0.48%) | 1,120<br>(1.64%) | 1,030<br>(1.51%) | 270<br>(0.40%) | 248<br>(0.36%) | 0.0023 | 0.0022 |
| INT (20) | 14,499 | 14,184 | 470<br>(3.24%) | 428<br>(3.02%) | 385<br>(2.66%) | 318<br>(2.24%) | 76<br>(0.52%) | 66<br>(0.47%) | 0.0037 | 0.0032 |
| IGS (103) | 38,752 | 37,051 | 2,623<br>(6.77%) | 2,458<br>(6.63%) | 1,453<br>(3.75%) | 1,194<br>(3.22%) | 352<br>(0.91%) | 275<br>(0.74%) | 0.0056 | 0.0048 |
| Concat. | 121,368 | 119,302 | 3,710<br>(3.11%) | 3,421<br>(2.82%) | 3,958<br>(3.26%) | 2,542<br>(2.13%) | 698<br>(0.58%) | 589<br>(0.49%) | 0.0035 | 0.0031 |

**Table 4.** Alignment length, sequence variability, and ML tree statistics of the different datasets under study.
| (b) |  |  |  |  |  |  |  |  |  |
| --- | --- | --- | --- | --- | --- | --- | --- | --- | --- |
|  | Indel coding | Alignment length (bp) |  | Best ML tree likelihood (-lnL) |  | Best ML tree length |  | Avg. BS support per node |  |
|  |  | before | after | before | after | before | after | before | after |
| CDS (81) | no | 68,117 | 68,076 | 103,459.8 | 102,518.0 | 0.0193 | 0.0177 | 68% | 66% |
|  | yes | 68,150 | 68,108 | 103,736.0 | 102,783.0 | 0.0199 | 0.0182 | 68% | 65% |
| INT (20) | no | 14,499 | 14,184 | 23,560.9 | 22,187.8 | 0.0370 | 0.0273 | 45% | 44% |
|  | yes | 14,658 | 14,259 | 25,318.6 | 22,760.0 | 2.1675 | 0.0344 | 42% | 43% |
| IGS (103) | no | 38,752 | 37,042 | 66,780.7 | 60,729.1 | 0.0580 | 0.0430 | 65% | 63% |
|  | yes | 39,318 | 37,385 | 71,559.8 | 63,146.6 | 0.0831 | 0.0556 | 62% | 68% |
| Concat. | no | 121,368 | 119,302 | 194,716.3 | 186,078.8 | 0.0337 | 0.0266 | 67% | 73% |
|  | yes | 122,126 | 119,752 | 201,615.7 | 189,627.2 | 0.0432 | 0.0315 | 68% | 73% |

**Figure 4.**
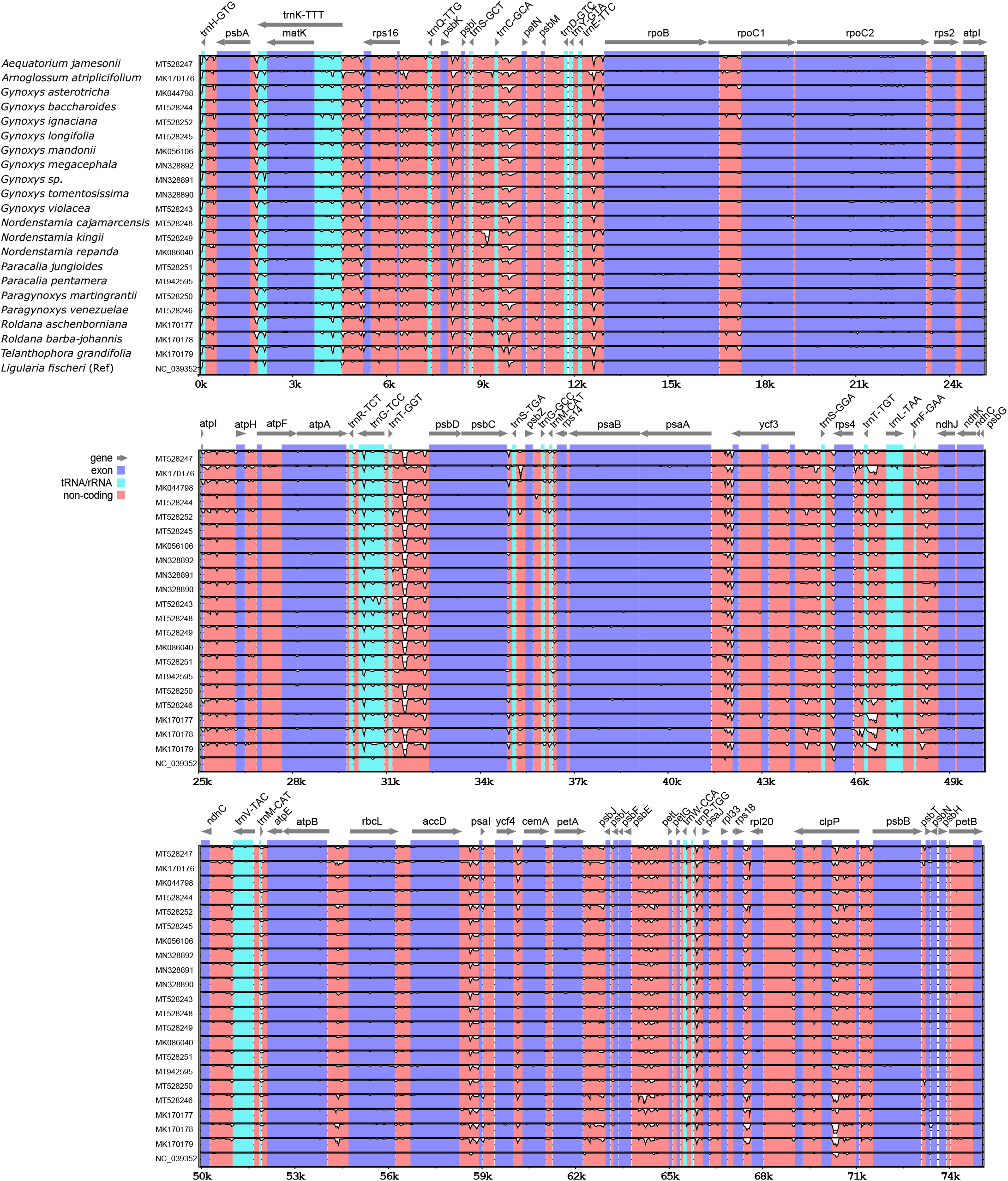
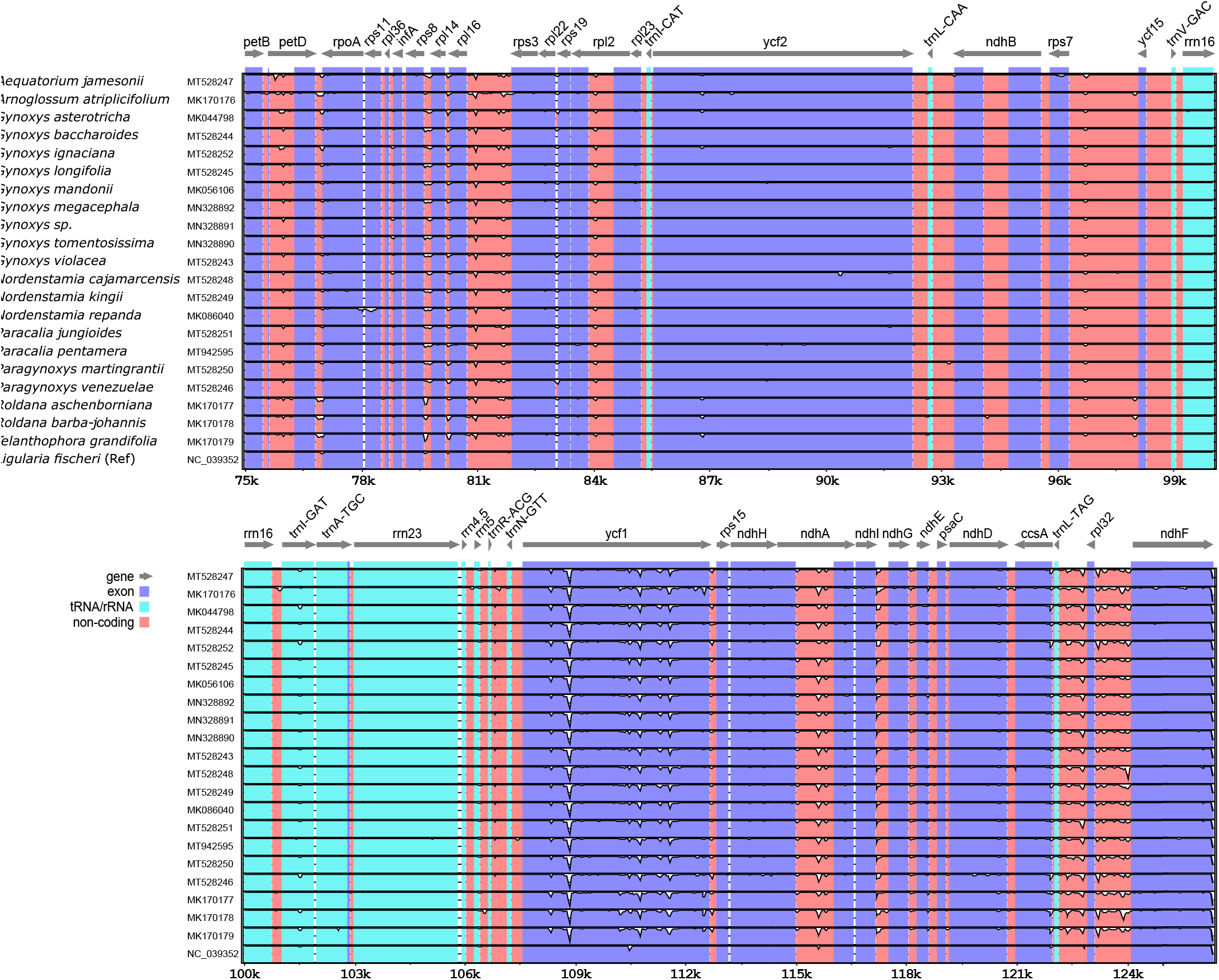
Visualization of sequence variability between the plastid genomes under study using mVISTA. The alignment was split into sequence batches of 25 kb length by mVISTA for easier visualization. Each lane represents a genome. In each lane, the proportion of missing similarity is indicated by white color, starting from the top of each lane. Coding regions are represented in blue, transfer and ribosomal RNAs in cyan, and non-coding regions in red. Gray arrows indicate the location and orientation of plastome genes.

### Effect of alignment adjustment on homoplasy indices

The evaluation and, when required, manual adjustment of the MSAs had a considerable effect on homoplasy in these alignments (Figure 5). Specifically, the MSAs of the intergenic spacers *atpB*–*rbcL, ndhC*–*trnV-TAC, psaA*–*ycf3, trnC-GCA*–*petN*, and *trnL-TAG*–*rpl32* exhibited considerably reduced levels of homoplasy upon alignment adjustment, with some spacer regions having their homoplasy index values improve by more than 50%. Improvements were particularly noticeable for values of the rescaled consistency index and the retention index among the intergenic spacers. The manual adjustment of the MSAs of the introns and the coding regions had a less intense impact but nonetheless resulted in reduced levels of homoplasy for the alignment of three introns and one coding region. Overall, these improvements in homoplasy levels due to alignment adjustments must be seen as conservative estimates, as alignment positions with indels were not taken into account during the calculation of the homoplasy indices.

**Figure 5.**
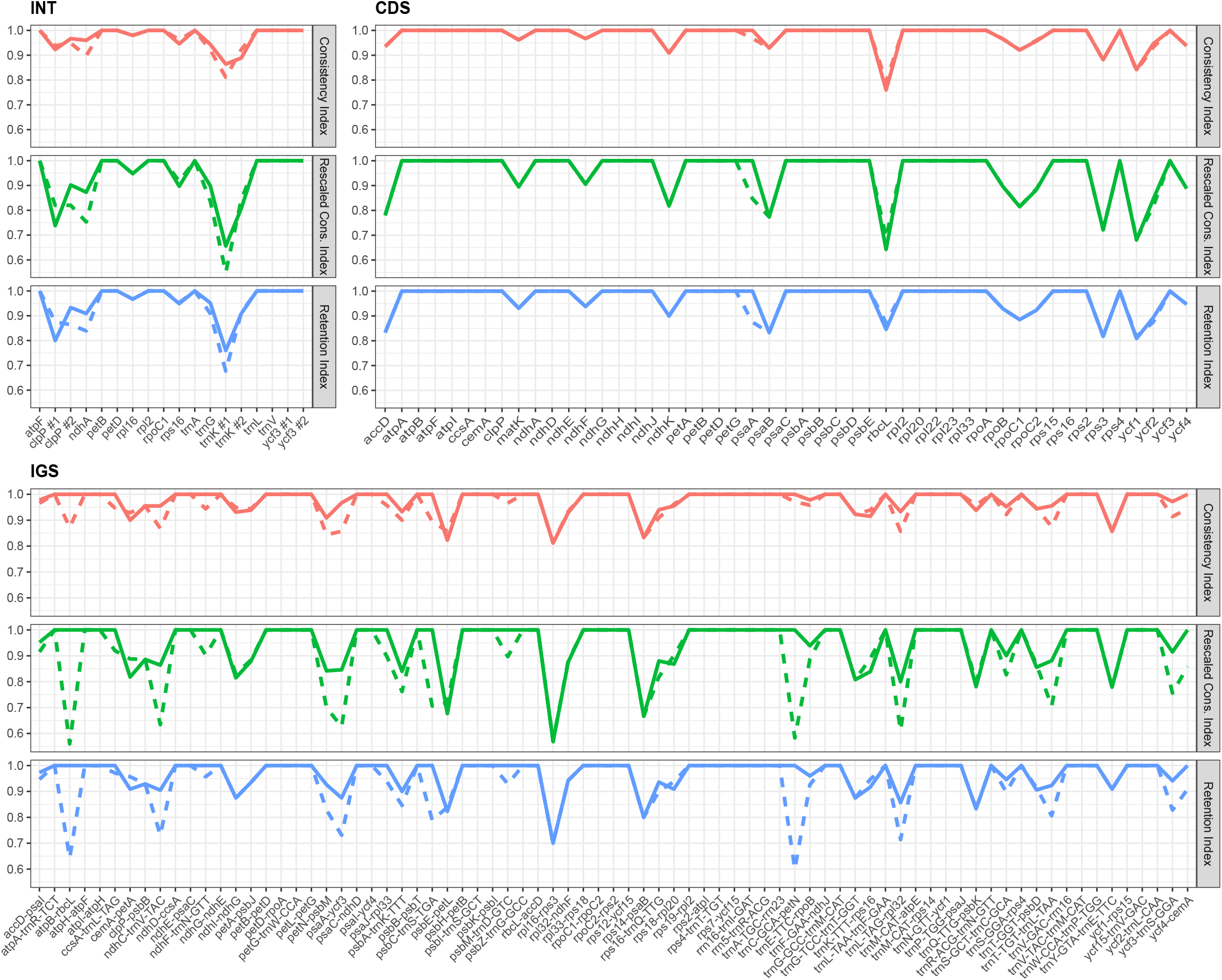
Comparison of the values of three homoplasy indices ‘consistency index’, ‘rescaled consistency index’, and ‘retention index’ across each MSA under study before (dashed lines) and after (full lines) alignment adjustment. The values of the consistency index are indicated in red, the values of the rescaled consistency index in green, and the values of the retention index in blue.

### Phylogenetic reconstructions

A comparison of the phylogenetic tree inference based on each of the three plastid partitions with tree inferences based on the concatenation of these partitions indicated that each partition contributed informative characters to the phylogenetic reconstruction, albeit in different proportions (Table 4). After alignment adjustment, the concatenation of all MSAs of the coding regions had a length of 68,076 bp, of which 1,030 (1.51%) were polymorphic sites; the concatenation of all MSAs of the introns had a length of 14,184 bp, of which 318 (2.24%) were polymorphic sites; and the concatenation of all MSAs of the intergenic spacers had a length of 37,033 bp after alignment adjustment, of which 1,194 (3.22%) were polymorphic sites. Consequently, the sequence matrix representing the concatenation of the three partitions had a total length of 119,311 bp after alignment adjustment, of which 113,059 (94.8%) were monomorphic sites, 2,542 (2.1%) were polymorphic sites, and 3,710 (3.1%) gaps or sites of missing data. Among the polymorphic sites, a total of 589 (23.2%) were parsimony informative, of which 248 (42.1%) originated in the coding regions, 275 (46.7%) in the intergenic spacers, and 66 (11.2%) in the introns.

In two instances, small but noticeable differences in the results of our phylogenetic reconstructions were identified when compared to the different tree inference methods and the coding of indels. First, the phylogenetic reconstructions conducted under different inference methods generated different trees, particularly with respect to node support. Node support for reconstructions under BI, for example, was generally high across the inferred trees, whereas node support for reconstructions under ML was considerably lower for many of earlier diverging nodes (Figures 6,7). Second, the coding of indels had a small but noticeable impact on tree inference; many of the trees exhibited minor topological changes upon inclusion of an indel coding matrix in the phylogenetic reconstruction, but the overall relationships, as well as the node support, exhibited minor differences. The phylogenetic trees reconstructed under the concatenation of all coding regions, experienced no topological changes upon the coding of indels, whereas trees reconstructed under the concatenation of all introns and all intergenic spacers exhibited differences in at least one node. Interestingly, the coding of indels caused considerably more topological changes in reconstructions under ML than under BI (Appendices S4–S7 in the Supplemental Materials).

**Figure 6.**
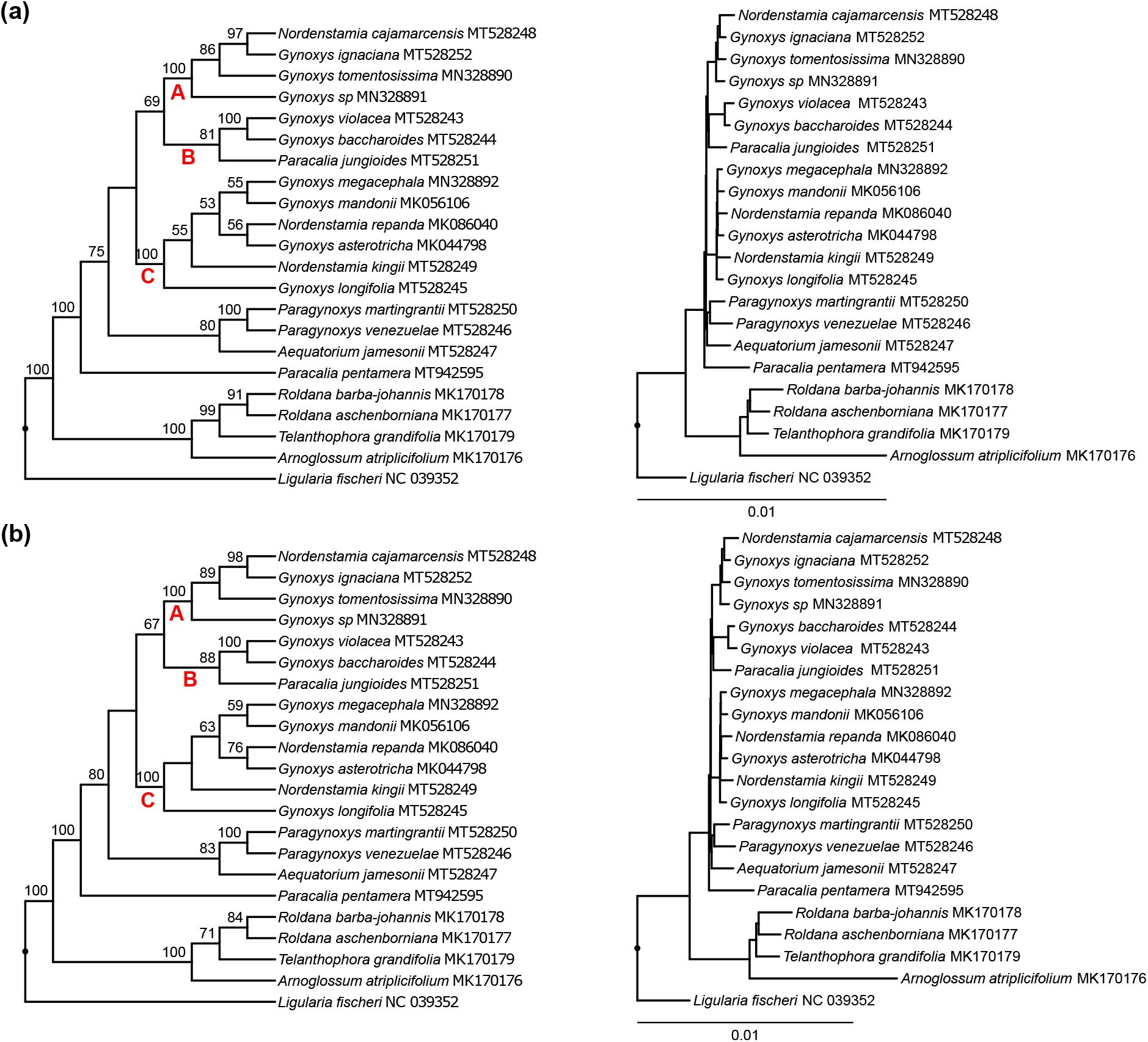
Results of the phylogenetic tree inference via ML on the concatenation of all three plastid partitions after alignment adjustment. The trees displayed represent the trees with the highest likelihood score (a) without and (b) with the coding of indels. The inferred relationships are visualized as cladograms with statistical node support (left) and corresponding phylograms with exact branch lengths (right). Bootstrap support values greater than 50% are given above the branches of each cladogram. Three clades (i.e., “A”, “B”, and “C”) are highlighted by red letters located next to the most recent common ancestor of each clade. All trees were rooted with using *Ligularia fischeri* as outgroup.

**Figure 7.**
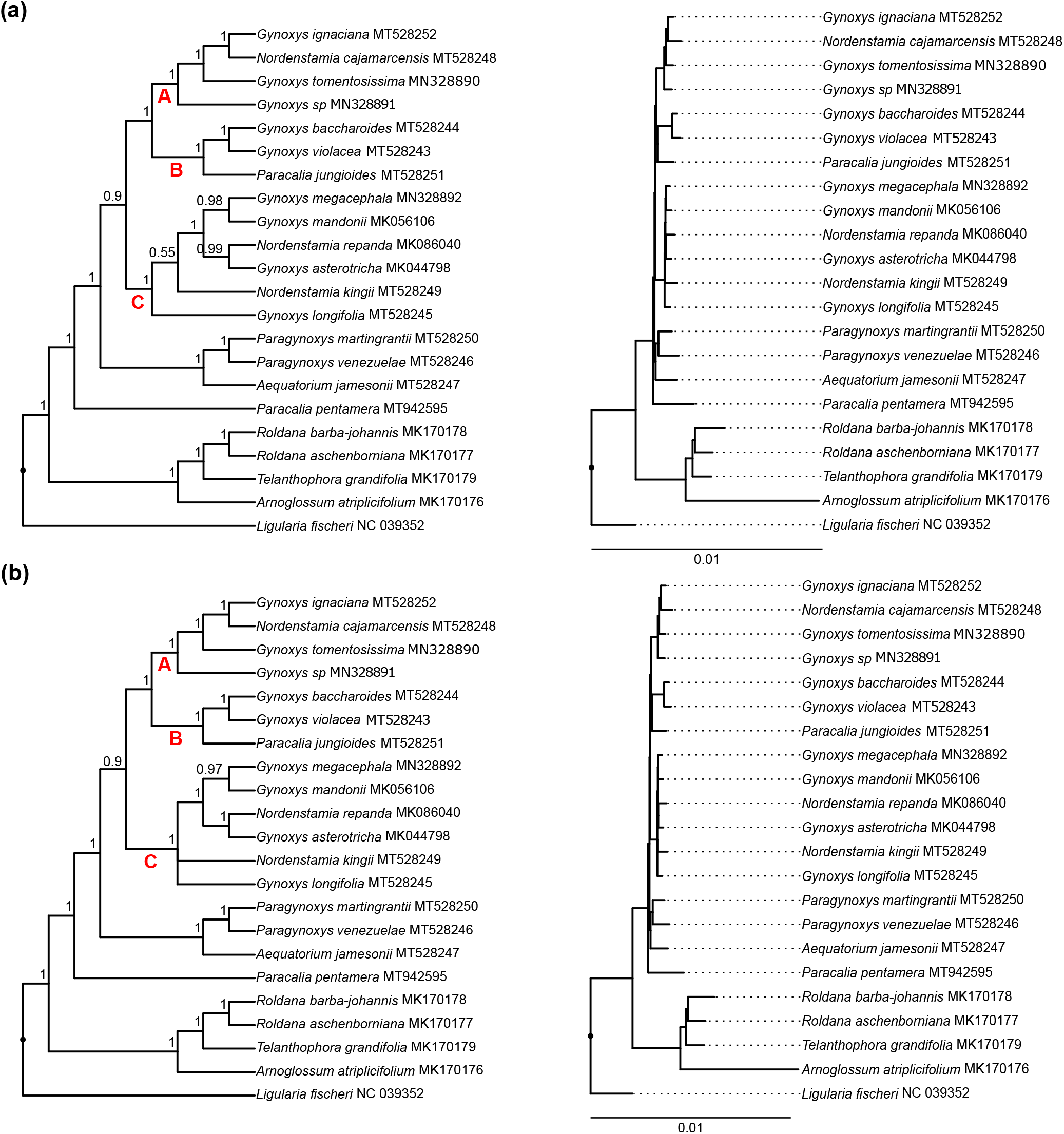
Results of the phylogenetic tree inference via BI on the concatenation of all three plastid partitions after alignment adjustment (a) without and (b) with the coding of indels. The trees displayed represent the 50% majority-rule consensus tree of the posterior tree distribution, visualized as cladograms with posterior probability values greater than 0.5 (left) and corresponding phylograms with exact branch lengths (right). All settings are as in Figure 6.

Considerable differences in the results of our phylogenetic reconstructions were identified based upon the different plastid partitions as well as the adjustment of alignments. The phylogenetic reconstructions based on different plastid partitions resulted in trees with different topologies and node support; the tree topologies inferred under the concatenation of all introns were notably different from the topologies inferred under the concatenation of all coding regions and intergenic spacers and also exhibited lower node support (Appendices S4 and S6). The average BS support per node in the best ML inferred under the concatenation of all introns was considerably lower than the average BS support per node in the trees inferred on the concatenation of all coding regions or all intergenic spacers. Only the sister relationship between the Gynoxoid group and the North and Central American members of the Tussilaginineae was consistently retrieved across the plastid partitions. Furthermore, the phylogenetic reconstructions before and after the adjustment of alignments generally resulted in trees with different topologies and node support. The best ML trees based on the concatenation of all coding regions produced different topologies before and after alignment adjustment, while BS support was *>*50% for all but one node (Appendices S4). In the tree inferred before alignment adjustment, *Gynoxys longifolia* Wedd. was sister to *Nordenstamia kingii* (H. Rob. & Cuatrec.) B. Nord. (BS 59%), and a clade comprising *Paragynoxys* and *Aequatorium* was sister to a clade of *G. baccharoides* (Kunth) Cass. and *G. violacea* Sch. Bip. ex Wedd. (BS 52%). In the tree inferred after alignment adjustment, fewer nodes with BS support *>*50% were retrieved and none of the previously identified relationships were supported. The best ML trees of the concatenation of all introns also exhibited different topologies before and after alignment adjustment, with BS support primarily *>*50%. By contrast, the best ML trees of the concatenation of all intergenic spacers inferred before and after alignment adjustment were highly similar in topology and generally recovered BS support *>*50%. In summary, the reconstruction of the phylogenetic relationships of the Gynoxoid group was strongly dependent on the plastid partition employed and the adjustment of the alignments. Moreover, our results demonstrated that each of the three plastid partitions alone was insufficient to fully resolve the phylogenetic relationships of this species group (Appendices S4 and S6).

Phylogenetic reconstruction on the concatenation of all plastid partitions resulted in trees that were consistent in topology and similar in node support across different tree inference methods and the coding of indels after the adjustment of alignments (Figures 6,7), but inconsistent in topology and node support before these adjustments (Appendices S5 and S7). Except for the most recent common ancestor of the two clades containing *Gynoxys*, and the node connecting *Nordenstamia kingii* and *Gynoxys megacephala* Rusby, all nodes of the best ML tree based on the concatenation of all plastid partitions exhibited some BS support upon alignment adjustment (Figure 6). In the 50% majority-rule consensus tree of the posterior tree distribution of the BI, most nodes contained high PP values (Figure 7). By contrast, the best ML tree inferred before the adjustment of alignments exhibited lower BS support as well as topological incongruence regarding the positions of *Gynoxys mandonii* Sch. Bip. Ex Rusby, *G. longifolia, G. megacephala*, and *Nordenstamia kingii* and the paraphyly of *Roldana* with respect to *Telantophora* (Appendix S6). These differences in tree topology before and after the adjustment of alignments were even more pronounced under BI and may be discordant given the high node support received under this optimality criterion (Appendix S7).

### Inference of relationships

The phylogenetic relationships recovered through our tree inferences on the concatenation of all plastid partitions resulted in high node support for most clades (Figures 6,7). The Gynoxoid group were recovered as a clade with maximum BS and PP support, comprising the genera *Aequatorium, Gynoxys, Nordenstamia, Paracalia*, and *Paragynoxys*. The non-Gynoxoid taxa of the Tussilagininae (i.e., *Roldana, Telanthophora*, and *Arnoglossum*) were recovered as sister to this Gynoxoid group in each reconstruction and with maximum support. *Paracalia pentamera* (Cuatrec.) Cuatrec., which constitutes the type species in the genus, was recovered as the earliest diverging lineage of the Gynoxoid group in each inference again with high node support. Except for *Paragynoxys*, all other genera of the Gynoxoid group that are represented by two or more species were found to be non-monophyletic. Specifically, *Paragynoxys martingrantii* (Cuatrec.) Cuatrec. and *P. venezuelae* (V.M. Badillo) Cuatrec. were recovered as sister species with maximum node support and identified to be sister to the specimen of *Aequatorium* included in this study (medium BS support, maximum PP support). *Gynoxys* and *Nordenstamia* were found to be highly polyphyletic and recovered in two separate subclades. One subclade comprised four species of *Gynoxys* and two species of *Nordenstamia*, and exhibited maximum node support under both ML and BI. Specifically, it comprised *Gynoxys asterotricha* Sch. Bip., *G. longifolia, G. mandonii*, and *G. megacephala*, as well as *Nordenstamia kingii* and *N. repanda* (Wedd.) Lundin, the latter of which constitutes the type species for the genus. However, the most recent common ancestor of *Nordenstamia kingii* and *Gynoxys megacephala* was poorly supported in this subclade and a part of a polytomy under BI when coding indels. The other subclade comprised five species of *Gynoxys*, one species of *Nordenstamia*, and one species of *Paracalia*, and had maximum PP but only medium BS support (BS*>*67). This clade comprised *G. baccharoides*, which constitutes the type species for *Gynoxys, G. ignaciana* Cuatrec., *G. tomentosissima, G. violacea*, and an as-of-yet unidentified species of *Gynoxys* (i.e., *Gynoxys sp*. SEN301). Moreover, the clade comprised *Paracalia jungioides* (Hook. & Arn.) Cuatrec. and *Nordenstamia cajamarcensis* (H. Rob. & Cuatrec.) B. Nord. The species *Paracalia jungioides, Gynoxys baccharoides*, and *G. violacea* formed a clade under both ML and BI, which exhibited maximum PP and high BS support (BS*>*81). The other species of this subclade were also recovered as a monophyletic group with maximum node support.

## DISCUSSION

### Plastid genomes of Senecioneae

The present investigation is the first to compare plastid genomes from different genera of the Tussilagininae and to employ their sequences in phylogenetic analysis. To date, 29 complete plastid genomes of the Senecioneae have been sequenced and are available through NCBI Nucleotide (accessed on 01-Dec-2020), representing five different genera and two of the four subtribes. Specifically, plastid genomes have been sequenced for *Dendrosenecio* (Hauman ex Hedberg) B. Nord., *Jacobaea* Mill., *Pericallis* D. Don, and *Senecio* L. of the subtribe Senecioninae, and *Ligularia* Cass. of the subtribe Tussilagininae. Gichira et al. (2019), sequenced and compared eleven plastid genomes of *Dendrosenecio* (Hauman ex Hedberg) B. Nord. and *Senecio* to identify variable regions for phylogenetic analysis. Similarly, Chen et al. (2018) generated plastid genomes of six species of *Ligularia* to examine the utility of these genomes as barcodes for identifying individual species. The present investigation expands the list of sequenced plastid genomes of the Tussilagininae by eight genera. All plastid genomes sequenced in this study display a highly conserved genome structure and exhibit the two large inversions that separate the Asteraceae from other flowering plant families (Kim et al., 2005). The size range of these newly sequenced plastid genomes and their gene order and content is highly similar to other Senecioneae (Gichira et al., 2019; Chen et al., 2018).

### Phylogenetic position of the Gynoxoid clade

The results of our phylogenetic reconstructions corroborate the previously supported relationships of the Tussilaginineae as inferred by Pelser et al. (2007). Specifically, our results support the phylogenetic relationships of the *Aequatorium*–*Arnoglossum* clade that were reported by Pelser et al. (2007) and confirm the Gynoxoid group as being monophyletic with high statistical support (Figures 6,7). While previous studies had already reported the Gynoxoid group as a clade (e.g., Pelser et al., 2010; Quedensley et al., 2018, both primarily based on ITS DNA sequences), their sampling of taxa and genomic regions was generally insufficient to infer the monophyly of the Gynoxoid group. Moreover, we recovered the Gynoxoid clade as sister to a clade of North and Central America taxa (i.e., *Arnoglossum, Telantophora*, and *Roldana*) that are distributed from the United States (Quedensley et al., 2018) to Panama (Funston, 2009; Clark and Pruski, 2015) and possibly Colombia (Calvo, 2016), and this sister relationship was previously illustrated (Pelser et al., 2010). Quedensley et al. (2018) reported that several genera in this sister clade to the Gynoxoid group were highly polyphyletic, and our results support this assessment, as we found *Roldana* non-monophyletic in several reconstructions (e.g., Appendix S7). Future studies on the phylogenetic relationships of the Tussilaginineae should further increase the taxon sampling.

### Phylogenetic relationships in the Gynoxoid clade

Our phylogenetic reconstructions recovered several relationships within the Gynoxoid clade with high clade support (Figures 6,7). For example, our results suggest that genus *Paracalia* is not monophyletic and that its type species (i.e., *Paracalia pentamera*) is sister to the rest of the Gynoxiod group, whereas *Paragynoxys* is monophyletic with full support. The monophyly of *Paragynoxys* is also supported by morphological characters such as discoid capitula and deep-lobed white corollas (Cuatrecasas, 1955). The reconstructions based on the full plastid genome revealed that *Aequatorium* and *Paragynoxys* were sister genera(1.0 PP and 80% ML-BS), whereas reconstructions based on individual plastid regions could not resolve this relationship (e.g., Pelser et al., 2010). Moreover, these complete plastome reconstructions identified three highly supported subclades within the Gynoxoid clade irrespective of the tree inference method applied: subclade A, which comprises *G. ignaciana, G. tomentosissima, G. sp*, and *Nordenstamia cajamarcensis* and is also supported by morphological characters such as opposite leaves and granular hairs along the veins on the upper face of the leaves; subclade B, which comprises *G. baccharioides, G. violacea*, and *Paracalia jungioides*; and subclade C, which comprises *G. megacephala, G. mandonii, Nordenstamia repanda, G. asterotricha, Nordemstamia kingii*, and *G. longifolia*. All internal nodes in these subclades were found to be well supported, except for the position of *N. kingii* in subclade C (weakly supported in the ML and BI trees when ignoring indel coding and unsupported upon inclusion of the indel coding matrix). The type species of *Gynoxys* and *Nordenstamia* (i.e., *Gynoxys baccharoides* and *Nordenstamia repanda*, respectively) were recovered in a clade in which the species of both genera did not segregate, indicating the non-monophyly of both genera.

### Importance of manual alignment adjustment

The present investigation highlights the importance of evaluating and, where necessary, adjusting software-generated MSAs prior to phylogenomic analysis. While different alignment algorithms have been implemented in the various software tools available for automatic DNA sequence alignment (e.g., Needleman-Wunsch algorithm in ClustalW; Thompson et al., 1994), their mechanistic processes often depart from the actual biological processes that shape the molecular evolution of DNA sequences. For example, biological processes often comprise the instantaneous insertion, deletion, inversion, or translocation of multiple nucleotides, yet many alignment algorithms cannot replicate these mechanisms (Graham et al., 2000; Borsch and Quandt, 2009; Ochoterena, 2008). Molecular phylogenetic studies, thus, often adjust their DNA sequence alignments manually using motif-based approaches. Such approaches were conceptualized as ‘motif alignments’ that follow defined rules (e.g., Kelchner, 2000; Loehne and Borsch, 2005; Morrison, 2006, 2015) and reflect putative microstructural mutations while assessing positional homology (de Pinna, 1991). Numerous investigations have demonstrated the impact that the selection of an alignment method can have on the reconstruction of phylogenetic trees (e.g., Morrison and Ellis, 1997; Simmons et al., 2010b,a; Wong et al., 2008), and simulation studies have shown that alignment accuracy is highly dependent on the frequency of genomic insertions and deletions (e.g., Pervez et al., 2014). Hence, the manual adjustment of DNA sequence alignments following a motif-based approach has become a common practice among molecular phylogenetic studies, particularly those based on single genetic markers. Phylogenomic studies, by contrast, often dismiss the practice of manual alignment adjustment, as the amount of sequence data under analysis is considered to outweigh the ability of any researcher to correct software-induced alignment errors for all but the smallest datasets (Wu et al., 2012). Instead, most phylogenomic studies often exclude those regions of an MSA that are deemed unreliable via positional filtering processes (e.g., Ali et al., 2019). While such approaches may eliminate some of the erroneous statements of positional homology among aligned nucleotides, they often fail in their removal of genomic regions that contain unrecognized inversions (Figure 2) or lack indels despite being non-homologous. The inadvertent analysis of DNA matrices that harbor sequence inversions is particularly problematic, as these inversions are highly homoplastic (Kelchner and Wendel, 1996) and often lead to spurious phylogenetic results (Joly et al., 2010). Unsurprisingly, the manual examination of the plastid phylogenomic dataset of this investigation for alignment errors has led to the identification of numerous sequence inversions that were not recognized at the stage of the software-driven sequence alignment (Figure 2, Table 2).

### Process of manual alignment adjustment

To alleviate the problem of missing positional homology upon automatic sequence alignment, we conducted visual examinations and manual corrections of the software-derived MSAs using a motif approach. Specifically, we removed the nucleotide positions from the final sequence matrix for which positional homology could not be established. For example, we removed or truncated length-variable poly-A/T microsatellites that originated through repeated and independent insertions of single or multiple nucleotides and did not form recognizable sequence motifs. Next to improving the positional homology for individual MSAs (Figure 5), this strategy allowed us to (i) identify and re-invert naturally occurring sequence inversions that cannot be automatically aligned and would introduce erroneous nucleotide polymorphisms if left unedited (Chen et al., 2016), and (ii) mask and, thus, exclude those nucleotide positions for which positional homology could not be reasonably identified. Time-efficient work was facilitated through automatically partitioning of annotated genomic regions into individual datasets, which could then be edited manually in PhyDE without affecting the overall alignment. Upon alignment adjustment, individual MSAs were then concatenated automatically using the pipeline of Gruenstaeudl et al. (2018). The success of our manual alignment correction strategy was demonstrated by the improvement of homoplasy indices upon alignment adjustment (Figure 5). The change in the level of homoplasy was also reflected by the tree topologies of the concatenated MSAs when compared before and after alignment adjustment (Table 4b; Appendices S4–S7 in the Supplemental Materials). Interestingly, the use of an uncorrected alignment had an effect on the inferred topology within subclade C of the core Gynoxoid clade and the *Arnoglossum*–*Roldana* clade (compare Figures 6,7 to Appendices S6–S7) both under BI and ML and with and without the indel matrix. When phylogenetic trees were inferred from any of the plastid partitions individually (i.e., genes, introns, and intergenic spacers alone), these effects were only visible in the *Arnoglossum*–*Roldana* clade, since nodes within the shallow core Gynoxoid clade were not receiving sufficient statistic confidence to allow a comparison. Plastid phylogenomic reconstructions should, thus, consider the evaluation and, where necessary, correction of automatic alignment results.

### Mosaic-like evolution of plastid genomes

The large majority of angiosperm plastid genomes display a highly conserved structure, uniparental inheritance, and a general absence of recombination between chromosomes (Marechal and Brisson 2010, but see Ruhlman et al. 2017; Li et al. 2020). Hence, many researchers assume that the plastid genome evolves as a single linkage unit (Bock, 2007), with different genomic regions sharing the same evolutionary history (e.g., Lu et al., 2018). However, recent studies have reported widespread phylogenetic incongruence between different regions of the plastid genome (e.g., Goncalves et al., 2019; Gruenstaeudl, 2019; Walker et al., 2019), which indicates that the plastid genome may not represent a homogeneous genetic locus. This phylogenetic incongruence may partially be the result of the different mutation rates and selective constraints across the plastid genome, which is illustrated by the co-existence of the relatively slowly evolving gene *rbcL*, the more rapidly evolving gene *matK* with nearly equal site rates in all three codon positions, and the even faster evolving non-coding markers *trnL* intron, *trnT–L* intergenic spacer, and *trnL–F* intergenic spacer in the same genome (Müller et al., 2006). Examples for selective constraints are directed selection on certain nucleotide positions and compensatory base changes (Kelchner, 2002; Borsch et al., 2003), which can cause patterns of homoplasy and lead to a spurious phylogenetic signal, especially when the number of variable nucleotides is limited. This observed phylogenetic incongruence may also be the result of the different structural constraints in the plastid genome, which comprises a succession of conserved and variable elements that can display secondary DNA structure. For example, the highly variable and AT-rich stem loops of non-coding plastid DNA often exhibit accelerated, lineage-specific nucleotide substitution rates and a high frequency of microstructural mutations (Korotkova et al., 2014). The presence of phylogenetic incongruence due to systematic error, such as the selection of incorrect nucleotide substitution models or incorrect homology statements during MSA (e.g., Zhang et al., 2020), represents another possible explanation, and its extent is under active investigation (Goncalves et al., 2020). In summary, the plastid genome exhibits a mosaic-like pattern of molecular evolution, and if that pattern is not adequately modeled during phylogenetic reconstruction, the resulting inferences may be highly supported but largely spurious Walker et al. (2019); Thode et al. (2020); Zhang et al. (2020). Plastid phylogenomics is, thus, facing a new paradigm in which the differential phylogenetic signal of particular genome regions must be accounted for (Goncalves et al., 2020).

### Phylogenetic utility of different plastome regions

The different regions of the plastid genome exhibit different molecular constraints and, thus, a different utility for phylogenetic reconstructions. The coding regions of the plastid genome have traditionally been used for the reconstruction of deep-level phylogenetic relationships (Graham and Olmstead, 2000; Xi et al., 2012; Walker et al., 2019), but these may be biased when genes with incongruent phylogenetic signal coexist in the genome (e.g., Gruenstaeudl, 2019; Goncalves et al., 2019; Walker et al., 2019). Goncalves et al. (2020), suggested to infer phylogenetic relationships on individual genes and to compare the resulting topologies to the reconstructions based on the concatenation of all coding regions. Since many regions of the plastid genome may exhibit an incongruent phylogenetic signal, this approach should be extended to the complete genome length. The non-coding regions of the plastid genome experience only limited selective pressure and are known to evolve at faster rates than the coding regions (Kim et al., 1999). Molecular phylogenetic studies, thus, preferentially employ non-coding genome regions to reconstruct the relationships of taxa that exhibit shallow levels of sequence divergence (Kelchner, 2002; Shaw et al., 2007; Androsiuk et al., 2020). Such levels are often found among recently radiated plant lineages, and the non-coding proportion of the plastid genome is particularly useful in acquiring phylogenetic resolution, as demonstrated here for the Gynoxoid group (Appendices S1–S3 in the Supplemental Materials). However, the presence of mutational hotspots and indels can impede the assessment of homology during the alignment of non-coding sequences, and this frequently leads to the loss of the phylogenetic signal (Barniske et al., 2012). Kelchner (2000) reviewed the molecular mechanisms under which the non-coding regions of the plastid genome evolve and argues that the established methodologies for phylogenetic inference may not always be suitable for non-coding DNA. Within non-coding DNA, introns are described as non-independent sequences whose mutational dynamics are restricted by their molecular structure and function (Kelchner, 2002). Accordingly, Brozynska et al. (2016) reported that the least variable regions of the plastid genome of *Oryza* are found among introns. However, most phylogenetic studies have demonstrated that introns can contain a level of variability for molecular phylogenetic investigations sufficient for low-level phylogenetic reconstructions (Creer, 2007). The intron of *rpl16*, for instance, is one of the fastest evolving introns in the plastid genome of land plants and has been used extensively for reconstructing phylogenetic relationships at the inter-and infragenic level (Kelchner, 2002). To better explain the phylogenetic utility of introns, Borsch et al. (2003) suggested a mosaic-like sequence structure within introns that is governed by different splicing mechanisms and evolutionary constraints, and both Creer (2007) and Kelchner (2002) proposed methodologies for effective phylogenetic analyses with introns.

### Harnessing all regions of the plastid genome

The results of this investigation suggest that the analysis of all regions of the plastid genome is essential in acquiring a reliable phylogenetic reconstruction of recently diverged plant lineages. This conclusion can likely be generalized to many plastid phylogenomic studies at the species level, as the phylogenetic signal of any plastid partition individually (i.e., genes, introns, and intergenic spacers alone) is likely insufficient to retrieve fully resolved and well-supported trees. Several studies have demonstrated that phylogenetic tree inference is affected by the selection of plastid genomic regions employed (e.g., Lu et al., 2018; Wikstroem et al., 2020). Thode et al. (2020), for example, recovered congruent phylogenetic trees from coding and non-coding plastid regions of neotropical lianas, but the nodes exhibited stronger support values in reconstructions using non-coding sequences. Similarly, Koehler et al. (2020) discovered that only a subset of the plastid genome regions was particularly useful in the phylogenetic reconstruction of the subfamily Opuntioideae (Cactaceae). Furthermore, Zhang et al. (2020) encountered considerable phylogenetic incongruence in a phylogenomic analysis of 36 tribes of the Fabaceae, including incongruence among strongly supported nodes and in relation to coding versus non-coding sections of the plastid genome. Despite these findings, many plastid phylogenomic investigations ignore the non-coding sections of the plastid genome during phylogenetic tree inference (e.g., Ma et al., 2014; Ross et al., 2015), often due to challenges in the alignment of these regions (e.g., Zhang et al., 2017). This preferential selection of coding over non-coding regions in plastid phylogenomic studies may have led to reduced power in many reconstructions, and few, if any, biological reasons exist to exclude such genome regions from phylogenetic analysis. Our study contributes to the ongoing discussion of how the phylogenetic signal of the complete plastid genome can be partitioned and employed more effectively for phylogenetic inference (Thode et al., 2020; Koehler et al., 2020). More research is needed to address this question and should include different lineages of land plants and different levels of genetic distance among taxa and sequences.

## CONCLUSIONS

In this plastid phylogenomic investigation, we analyzed the phylogenetic relationships of the Gynoxoid group, an Andean lineage of the Asteraceae with low genetic distances between taxa. Our results indicated that at least two, and possibly three, of the five genera are polyphyletic. Moreover, our results illustrated that the group may correspond with the evolution of other rapidly radiating Andean lineages that represent “shallow clades” and for which a substantial amount of DNA sequence information is typically needed to obtain phylogenetic resolution. Furthermore, our results demonstrated that the inclusion of all plastid genome partitions was needed to infer fully resolved phylogenetic trees of the Gynoxoid group and that manual correction of sequence alignments had a considerable effect on tree inference. Our results also indicated that the adjustment of software-derived DNA sequence alignments may constitute an important step toward improved phylogenetic analyses in plastid phylogenomic studies. Specifically, the same standards of DNA sequence alignment and matrix construction that have been applied in studies of individual genomic regions should also be applied to plastid phylogenomic data sets. The impact of incorrect positional homology in a sequence matrix may be particularly severe among plastid datasets with low genetic distances, as the misaligned regions may contain a high proportion of the potentially informative sites. Consequently, species-level investigations that require the analysis of complete plastid genomes to resolve phylogenetic relationships should apply the utmost rigor in the motif-based alignment of nucleotide sequences and consider excluding areas of uncertain homology from their alignments. Overall, the Gynoxoid group may be an apt model system among recently diverged Andean plant lineages to study speciation and diversification through time and space.

## SUPPORTING INFORMATION

**Appendix S1–S3**. Alignment metrics of (S1) the coding sequences, (S2) the introns, and (S3) the intergenic spacers of the plastid genomes before and after alignment adjustment. **Appendix S4**. Results of phylogenetic tree inference via ML on the concatenation of all MSAs of the coding regions, the introns, or the intergenic spacers before and after alignment adjustment. **Appendix S5**. Results of the phylogenetic tree inference via ML on the concatenation of all three plastid partitions before alignment adjustment. **Appendix S6**. Results of phylogenetic tree inference via BI on the concatenation of all MSAs of the coding regions, the introns, or the intergenic spacers before and after alignment adjustment. **Appendix S7**. Results of the phylogenetic tree inference via BI on the concatenation of all three plastid partitions before alignment adjustment.

## DECLARATIONS

### Ethics approval and consent to participate

Not applicable.

### Consent for publication

Not applicable.

### Availability of data and materials

All datasets analyzed during the present investigation are available from Zenodo at DOI 10.5281/zen- odo.4428211.

## Author contributions

BE and TSQ conducted the field work, generated herbarium vouchers, and extracted DNA. BE and MG conducted the lab work, and assembled and annotated the plastid genomes. BE and TB conducted the visual inspection and adjustment of sequence alignments. BE and MG conducted the phylogenetic analyses and generated all figures and tables. BE and MG lead the writing of the manuscript, with additional contributions by TB and TSQ. All authors approved the final version of the manuscript.

## Acknowledgements

The authors thank Norbert Kilian, Tilo Henning, Daniel Montesinos, Stephan Beck, Carla Maldonado, and Huber Villca for assistance during sample collection. The authors also thank Sabine Scheel of the Freie Universität Berlin for assistance with DNA isolations. Moreover, the authors thank Robert K. Jansen of the University of Texas at Austin for guidance in the sequencing and assembly of several plastid genomes presented here. The authors acknowledge the high-performance computing service of the ZEDAT of the Freie Universität Berlin for providing allocations of computing time.

