## Supporting Information: Appendix S1-S7 for "Plastid phylogenomics of the Gynoxoid group (Senecioneae, Asteraceae) highlights the importance of motif-based sequence alignment amid low genetic distances"

**Appendix S1**. Alignment metrics and homoplasy indices of the MSAs of the coding regions before and after alignment adjustment. The columns are: alignment length, GC content, fraction of polymorphic sites, fraction of parsimony informative sites, consistency index, retention index, rescaled consistency index, maximum uncorrected p-distance. The table is ordered by the value of the rescaled consistency index, but the concatenated dataset of all markers is always on top. Fractions refer to the full alignment length.

| | Alignm. (bp) | | GC content | | Polymorph. sites | | Pars. inform. sites | | Consist. index | | Retent. index | | Rescaled index | | Uncorr. p-dist. | |
|---|---|---|---|---|---|---|---|---|---|---|---|---|---|---|---|---|
| | before | after | before | after | before | after | before | after | before | after | before | after | before | after | before | after |
| 81 CDS | 68,118 | 68,076 | 0.38 | 0.38 | 0.02 | 0.03 | 0.00 | 0.00 | 0.92 | 0.92 | 0.87 | 0.87 | 0.80 | 0.80 | 0.09 | 0.09 |
| *atpA* | 1,524 | 1,524 | 0.41 | 0.41 | 0.01 | 0.01 | 0.00 | 0.00 | 1.00 | 1.00 | 1.00 | 1.00 | 1.00 | 1.00 | 0.09 | 0.09 |
| *atpB* | 1,494 | 1,494 | 0.42 | 0.42 | 0.01 | 0.01 | 0.00 | 0.00 | 1.00 | 1.00 | 1.00 | 1.00 | 1.00 | 1.00 | 0.08 | 0.08 |
| *atpF* | 594 | 594 | 0.37 | 0.37 | 0.01 | 0.01 | 0.00 | 0.00 | 1.00 | 1.00 | 1.00 | 1.00 | 1.00 | 1.00 | 0.08 | 0.08 |
| *atpI* | 741 | 741 | 0.38 | 0.38 | 0.02 | 0.02 | 0.00 | 0.00 | 1.00 | 1.00 | 1.00 | 1.00 | 1.00 | 1.00 | 0.12 | 0.12 |
| *ccsA* | 966 | 966 | 0.32 | 0.32 | 0.02 | 0.02 | 0.00 | 0.00 | 1.00 | 1.00 | 1.00 | 1.00 | 1.00 | 1.00 | 0.11 | 0.11 |
| *cemA* | 687 | 687 | 0.33 | 0.33 | 0.02 | 0.02 | 0.00 | 0.00 | 1.00 | 1.00 | 1.00 | 1.00 | 1.00 | 1.00 | 0.10 | 0.10 |
| *clpP* | 585 | 585 | 0.43 | 0.43 | 0.01 | 0.01 | 0.00 | 0.00 | 1.00 | 1.00 | 1.00 | 1.00 | 1.00 | 1.00 | 0.10 | 0.10 |
| *ndhA* | 1,101 | 1,101 | 0.34 | 0.34 | 0.01 | 0.01 | 0.00 | 0.00 | 1.00 | 1.00 | 1.00 | 1.00 | 1.00 | 1.00 | 0.10 | 0.10 |
| *ndhD* | 1,503 | 1,503 | 0.36 | 0.36 | 0.02 | 0.02 | 0.00 | 0.00 | 1.00 | 1.00 | 1.00 | 1.00 | 1.00 | 1.00 | 0.11 | 0.11 |
| *ndhE* | 303 | 303 | 0.32 | 0.32 | 0.01 | 0.01 | 0.00 | 0.00 | 1.00 | 1.00 | 1.00 | 1.00 | 1.00 | 1.00 | 0.12 | 0.12 |
| *ndhG* | 528 | 528 | 0.35 | 0.35 | 0.02 | 0.02 | 0.01 | 0.01 | 1.00 | 1.00 | 1.00 | 1.00 | 1.00 | 1.00 | 0.12 | 0.12 |
| *ndhH* | 1,176 | 1,176 | 0.38 | 0.38 | 0.02 | 0.02 | 0.00 | 0.00 | 1.00 | 1.00 | 1.00 | 1.00 | 1.00 | 1.00 | 0.11 | 0.11 |
| *ndhI* | 498 | 498 | 0.35 | 0.35 | 0.01 | 0.01 | 0.00 | 0.00 | 1.00 | 1.00 | 1.00 | 1.00 | 1.00 | 1.00 | 0.09 | 0.09 |
| *ndhJ* | 474 | 474 | 0.40 | 0.40 | 0.02 | 0.02 | 0.00 | 0.00 | 1.00 | 1.00 | 1.00 | 1.00 | 1.00 | 1.00 | 0.14 | 0.14 |
| *petA* | 960 | 960 | 0.41 | 0.41 | 0.01 | 0.01 | 0.00 | 0.00 | 1.00 | 1.00 | 1.00 | 1.00 | 1.00 | 1.00 | 0.07 | 0.07 |
| *petB* | 645 | 645 | 0.40 | 0.40 | 0.01 | 0.01 | 0.01 | 0.01 | 1.00 | 1.00 | 1.00 | 1.00 | 1.00 | 1.00 | 0.10 | 0.10 |
| *petD* | 531 | 531 | 0.40 | 0.40 | 0.02 | 0.02 | 0.01 | 0.01 | 1.00 | 1.00 | 1.00 | 1.00 | 1.00 | 1.00 | 0.11 | 0.11 |
| *petG* | 111 | 111 | 0.37 | 0.37 | 0.03 | 0.03 | 0.01 | 0.01 | 1.00 | 1.00 | 1.00 | 1.00 | 1.00 | 1.00 | 0.16 | 0.16 |
| *psaC* | 243 | 243 | 0.44 | 0.44 | 0.01 | 0.01 | 0.00 | 0.00 | 1.00 | 1.00 | 1.00 | 1.00 | 1.00 | 1.00 | 0.11 | 0.11 |
| *psbA* | 1,059 | 1,059 | 0.42 | 0.42 | 0.01 | 0.01 | 0.00 | 0.00 | 1.00 | 1.00 | 1.00 | 1.00 | 1.00 | 1.00 | 0.07 | 0.07 |
| *psbB* | 1,524 | 1,524 | 0.44 | 0.44 | 0.01 | 0.01 | 0.00 | 0.00 | 1.00 | 1.00 | 1.00 | 1.00 | 1.00 | 1.00 | 0.09 | 0.09 |
| *psbC* | 1,419 | 1,419 | 0.43 | 0.43 | 0.01 | 0.01 | 0.00 | 0.00 | 1.00 | 1.00 | 1.00 | 1.00 | 1.00 | 1.00 | 0.07 | 0.07 |
| *psbD* | 1,059 | 1,059 | 0.43 | 0.43 | 0.01 | 0.01 | 0.00 | 0.00 | 1.00 | 1.00 | 1.00 | 1.00 | 1.00 | 1.00 | 0.07 | 0.07 |

Continued on next page

Appendix S1 – continued from previous page

| | Alignm. (bp) | | GC content | | Polymorph. sites | | Pars. inform. sites | | Consist. index | | Retent. index | | Rescaled index | | Uncorr. p-dist. | |
|---|---|---|---|---|---|---|---|---|---|---|---|---|---|---|---|---|
| | before | after | before | after | before | after | before | after | before | after | before | after | before | after | before | after |
| *psbE* | 249 | 249 | 0.41 | 0.41 | 0.00 | 0.00 | 0.00 | 0.00 | 1.00 | 1.00 | 1.00 | 1.00 | 1.00 | 1.00 | 0.06 | 0.06 |
| *rpl2* | 825 | 825 | 0.44 | 0.44 | 0.01 | 0.01 | 0.00 | 0.00 | 1.00 | 1.00 | 1.00 | 1.00 | 1.00 | 1.00 | 0.08 | 0.08 |
| *rpl20* | 360 | 360 | 0.36 | 0.36 | 0.01 | 0.01 | 0.00 | 0.00 | 1.00 | 1.00 | 1.00 | 1.00 | 1.00 | 1.00 | 0.07 | 0.07 |
| *rpl22* | 468 | 468 | 0.35 | 0.35 | 0.02 | 0.02 | 0.01 | 0.01 | 1.00 | 1.00 | 1.00 | 1.00 | 1.00 | 1.00 | 0.09 | 0.09 |
| *rpl23* | 279 | 285 | 0.40 | 0.40 | 0.01 | 0.03 | 0.01 | 0.00 | 1.00 | 1.00 | 1.00 | 1.00 | 1.00 | 1.00 | 0.10 | 0.06 |
| *rpl33* | 198 | 198 | 0.41 | 0.41 | 0.02 | 0.02 | 0.01 | 0.01 | 1.00 | 1.00 | 1.00 | 1.00 | 1.00 | 1.00 | 0.14 | 0.14 |
| *rpoA* | 1,005 | 1,005 | 0.35 | 0.35 | 0.03 | 0.03 | 0.01 | 0.01 | 1.00 | 1.00 | 1.00 | 1.00 | 1.00 | 1.00 | 0.12 | 0.12 |
| *rps15* | 276 | 276 | 0.33 | 0.33 | 0.02 | 0.02 | 0.00 | 0.00 | 1.00 | 1.00 | 1.00 | 1.00 | 1.00 | 1.00 | 0.10 | 0.10 |
| *rps16* | 261 | 261 | 0.39 | 0.39 | 0.03 | 0.03 | 0.02 | 0.02 | 1.00 | 1.00 | 1.00 | 1.00 | 1.00 | 1.00 | 0.17 | 0.17 |
| *rps2* | 708 | 708 | 0.37 | 0.37 | 0.02 | 0.02 | 0.00 | 0.00 | 1.00 | 1.00 | 1.00 | 1.00 | 1.00 | 1.00 | 0.09 | 0.09 |
| *rps4* | 603 | 603 | 0.40 | 0.40 | 0.02 | 0.02 | 0.00 | 0.00 | 1.00 | 1.00 | 1.00 | 1.00 | 1.00 | 1.00 | 0.11 | 0.11 |
| *ycf3* | 504 | 504 | 0.39 | 0.39 | 0.01 | 0.01 | 0.01 | 0.01 | 1.00 | 1.00 | 1.00 | 1.00 | 1.00 | 1.00 | 0.06 | 0.06 |
| *ndhF* | 2,232 | 2,232 | 0.32 | 0.32 | 0.03 | 0.03 | 0.01 | 0.01 | 0.97 | 0.97 | 0.94 | 0.94 | 0.91 | 0.91 | 0.12 | 0.12 |
| *rpoB* | 3,18 | 3,198 | 0.39 | 0.39 | 0.02 | 0.02 | 0.00 | 0.00 | 0.97 | 0.96 | 0.93 | 0.93 | 0.90 | 0.90 | 0.10 | 0.10 |
| *matK* | 1,509 | 1,509 | 0.34 | 0.34 | 0.03 | 0.03 | 0.01 | 0.01 | 0.96 | 0.96 | 0.93 | 0.93 | 0.90 | 0.90 | 0.12 | 0.12 |
| *ycf4* | 552 | 552 | 0.39 | 0.39 | 0.03 | 0.03 | 0.01 | 0.01 | 0.94 | 0.94 | 0.95 | 0.95 | 0.89 | 0.89 | 0.13 | 0.13 |
| *rpoC2* | 4,146 | 4,146 | 0.38 | 0.38 | 0.02 | 0.02 | 0.00 | 0.00 | 0.96 | 0.96 | 0.92 | 0.92 | 0.89 | 0.88 | 0.10 | 0.10 |
| *psaA* | 2,250 | 2,250 | 0.43 | 0.43 | 0.01 | 0.01 | 0.00 | 0.00 | 0.97 | 1.00 | 0.88 | 1.00 | 0.85 | 1.00 | 0.09 | 0.08 |
| *ndhK* | 675 | 675 | 0.38 | 0.38 | 0.01 | 0.01 | 0.00 | 0.00 | 0.91 | 0.91 | 0.90 | 0.90 | 0.82 | 0.82 | 0.11 | 0.11 |
| *ycf2* | 6,675 | 6,689 | 0.38 | 0.38 | 0.01 | 0.01 | 0.00 | 0.00 | 0.93 | 0.95 | 0.88 | 0.89 | 0.82 | 0.84 | 0.06 | 0.06 |
| *rpoC1* | 2,088 | 2,088 | 0.38 | 0.38 | 0.02 | 0.02 | 0.01 | 0.01 | 0.92 | 0.92 | 0.89 | 0.89 | 0.81 | 0.81 | 0.10 | 0.10 |
| *accD* | 1,494 | 1,494 | 0.36 | 0.36 | 0.02 | 0.02 | 0.01 | 0.01 | 0.94 | 0.94 | 0.83 | 0.83 | 0.78 | 0.78 | 0.09 | 0.09 |
| *psaB* | 2,202 | 2,202 | 0.40 | 0.40 | 0.01 | 0.01 | 0.00 | 0.00 | 0.93 | 0.93 | 0.83 | 0.83 | 0.77 | 0.77 | 0.06 | 0.06 |
| *rps3* | 654 | 654 | 0.34 | 0.34 | 0.02 | 0.02 | 0.01 | 0.01 | 0.88 | 0.88 | 0.82 | 0.82 | 0.72 | 0.72 | 0.10 | 0.10 |
| *rbcL* | 1,455 | 1,455 | 0.44 | 0.44 | 0.01 | 0.06 | 0.01 | 0.01 | 0.79 | 0.76 | 0.87 | 0.85 | 0.69 | 0.64 | 0.09 | 0.09 |
| *ycf1* | 5,151 | 5,074 | 0.29 | 0.29 | 0.08 | 0.07 | 0.01 | 0.01 | 0.84 | 0.84 | 0.82 | 0.81 | 0.69 | 0.68 | 0.14 | 0.13 |
| *atpE* | 399 | 399 | 0.41 | 0.41 | 0.01 | 0.01 | 0.00 | 0.00 | 1.00 | 1.00 | n.a. | n.a. | n.a. | n.a. | 0.07 | 0.07 |
| *atpH* | 243 | 243 | 0.45 | 0.45 | 0.01 | 0.01 | 0.00 | 0.00 | 1.00 | 1.00 | n.a. | n.a. | n.a. | n.a. | 0.09 | 0.09 |
| *infA* | 231 | 231 | 0.37 | 0.37 | 0.01 | 0.01 | 0.00 | 0.00 | 1.00 | 1.00 | n.a. | n.a. | n.a. | n.a. | 0.09 | 0.09 |

Continued on next page

Appendix S1 – continued from previous page

| | Alignm. (bp) | | GC content | | Polymorph. sites | | Pars. inform. sites | | Consist. index | | Retent. index | | Rescaled index | | Uncorr. p-dist. | |
|---|---|---|---|---|---|---|---|---|---|---|---|---|---|---|---|---|
| | before | after | before | after | before | after | before | after | before | after | before | after | before | after | before | after |
| *ndhB* | 1,530 | 1,530 | 0.37 | 0.37 | 0.00 | 0.00 | 0.00 | 0.00 | 1.00 | 1.00 | n.a. | n.a. | n.a. | n.a. | 0.05 | 0.05 |
| *ndhC* | 360 | 360 | 0.35 | 0.35 | 0.00 | 0.00 | 0.00 | 0.00 | n.a. | n.a. | n.a. | n.a. | n.a. | n.a. | 0.00 | 0.00 |
| *petL* | 93 | 93 | 0.34 | 0.34 | 0.01 | 0.01 | 0.00 | 0.00 | 1.00 | 1.00 | n.a. | n.a. | n.a. | n.a. | 0.10 | 0.10 |
| *petN* | 87 | 87 | 0.41 | 0.41 | 0.01 | 0.01 | 0.00 | 0.00 | 1.00 | 1.00 | n.a. | n.a. | n.a. | n.a. | 0.11 | 0.11 |
| *psaI* | 108 | 108 | 0.37 | 0.37 | 0.02 | 0.02 | 0.00 | 0.00 | 1.00 | 1.00 | n.a. | n.a. | n.a. | n.a. | 0.14 | 0.14 |
| *psaJ* | 132 | 132 | 0.40 | 0.40 | 0.00 | 0.00 | 0.00 | 0.00 | n.a. | n.a. | n.a. | n.a. | n.a. | n.a. | 0.00 | 0.00 |
| *psbF* | 117 | 117 | 0.44 | 0.44 | 0.00 | 0.00 | 0.00 | 0.00 | n.a. | n.a. | n.a. | n.a. | n.a. | n.a. | 0.00 | 0.00 |
| *psbG* | 57 | 57 | 0.28 | 0.28 | 0.05 | 0.02 | 0.00 | 0.00 | 1.00 | n.a. | n.a. | n.a. | n.a. | n.a. | 0.24 | 0.00 |
| *psbH* | 237 | 237 | 0.38 | 0.38 | 0.01 | 0.01 | 0.00 | 0.00 | 1.00 | 1.00 | n.a. | n.a. | n.a. | n.a. | 0.09 | 0.09 |
| *psbI* | 108 | 108 | 0.37 | 0.37 | 0.02 | 0.02 | 0.00 | 0.00 | 1.00 | 1.00 | n.a. | n.a. | n.a. | n.a. | 0.14 | 0.14 |
| *psbJ* | 120 | 120 | 0.42 | 0.42 | 0.00 | 0.00 | 0.00 | 0.00 | n.a. | n.a. | n.a. | n.a. | n.a. | n.a. | 0.00 | 0.00 |
| *psbK* | 177 | 177 | 0.38 | 0.38 | 0.01 | 0.01 | 0.00 | 0.00 | 1.00 | 1.00 | n.a. | n.a. | n.a. | n.a. | 0.11 | 0.11 |
| *psbL* | 114 | 114 | 0.34 | 0.34 | 0.00 | 0.00 | 0.00 | 0.00 | n.a. | n.a. | n.a. | n.a. | n.a. | n.a. | 0.00 | 0.00 |
| *psbM* | 102 | 102 | 0.28 | 0.28 | 0.00 | 0.00 | 0.00 | 0.00 | n.a. | n.a. | n.a. | n.a. | n.a. | n.a. | 0.00 | 0.00 |
| *psbN* | 138 | 138 | 0.43 | 0.43 | 0.00 | 0.00 | 0.00 | 0.00 | n.a. | n.a. | n.a. | n.a. | n.a. | n.a. | 0.00 | 0.00 |
| *psbT* | 99 | 99 | 0.33 | 0.33 | 0.01 | 0.01 | 0.00 | 0.00 | 1.00 | 1.00 | n.a. | n.a. | n.a. | n.a. | 0.10 | 0.10 |
| *psbZ* | 186 | 186 | 0.36 | 0.36 | 0.01 | 0.01 | 0.00 | 0.00 | 1.00 | 1.00 | n.a. | n.a. | n.a. | n.a. | 0.07 | 0.07 |
| *rpl14* | 366 | 366 | 0.40 | 0.40 | 0.01 | 0.01 | 0.00 | 0.00 | 1.00 | 1.00 | n.a. | n.a. | n.a. | n.a. | 0.07 | 0.07 |
| *rpl16* | 420 | 417 | 0.43 | 0.43 | 0.01 | 0.01 | 0.00 | 0.00 | 1.00 | 1.00 | n.a. | n.a. | n.a. | n.a. | 0.10 | 0.10 |
| *rpl32* | 162 | 162 | 0.34 | 0.34 | 0.01 | 0.01 | 0.00 | 0.00 | 1.00 | 1.00 | n.a. | n.a. | n.a. | n.a. | 0.08 | 0.08 |
| *rpl36* | 111 | 111 | 0.36 | 0.36 | 0.00 | 0.00 | 0.00 | 0.00 | n.a. | n.a. | n.a. | n.a. | n.a. | n.a. | 0.00 | 0.00 |
| *rps11* | 408 | 408 | 0.46 | 0.46 | 0.06 | 0.57 | 0.00 | 0.00 | 1.00 | 1.00 | n.a. | n.a. | n.a. | n.a. | 0.23 | 0.10 |
| *rps12* | 354 | 354 | 0.45 | 0.45 | 0.00 | 0.00 | 0.00 | 0.00 | 1.00 | 1.00 | n.a. | n.a. | n.a. | n.a. | 0.05 | 0.05 |
| *rps14* | 300 | 300 | 0.41 | 0.41 | 0.02 | 0.02 | 0.00 | 0.00 | 1.00 | 1.00 | n.a. | n.a. | n.a. | n.a. | 0.12 | 0.12 |
| *rps18* | 303 | 303 | 0.35 | 0.35 | 0.00 | 0.00 | 0.00 | 0.00 | 1.00 | 1.00 | n.a. | n.a. | n.a. | n.a. | 0.06 | 0.06 |
| *rps19* | 276 | 276 | 0.34 | 0.34 | 0.01 | 0.01 | 0.00 | 0.00 | 1.00 | 1.00 | n.a. | n.a. | n.a. | n.a. | 0.09 | 0.09 |
| *rps7* | 465 | 465 | 0.41 | 0.41 | 0.03 | 0.37 | 0.00 | 0.00 | 1.00 | 1.00 | n.a. | n.a. | n.a. | n.a. | 0.17 | 0.07 |
| *rps8* | 402 | 402 | 0.36 | 0.36 | 0.01 | 0.01 | 0.00 | 0.00 | 1.00 | 1.00 | n.a. | n.a. | n.a. | n.a. | 0.09 | 0.09 |
| *ycf15* | 189 | 189 | 0.49 | 0.49 | 0.00 | 0.00 | 0.00 | 0.00 | n.a. | n.a. | n.a. | n.a. | n.a. | n.a. | 0.00 | 0.00 |

**Appendix S2**. Alignment metrics and homoplasy indices of the MSAs of the introns before and after alignment adjustment. The columns are: alignment length, GC content, fraction of polymorphic sites, fraction of parsimony informative sites, consistency index, retention index, rescaled consistency index, maximum uncorrected p-distance. The table is ordered by the value of the rescaled consistency index, but the concatenated dataset of all markers is always on top. Fractions refer to the full alignment length.

| | Alignm. (bp) | | GC content | | Polymorph. sites | | Pars. inform. sites | | Consist. index | | Retent. index | | Rescaled index | | Uncorr. p-dist. | |
|---|---|---|---|---|---|---|---|---|---|---|---|---|---|---|---|---|
| | before | after | before | after | before | after | before | after | before | after | before | after | before | after | before | after |
| 20 INT | 14,499 | 14,184 | 0.35 | 0.36 | 0.06 | 0.05 | 0.01 | 0.01 | 0.90 | 0.92 | 0.84 | 0.87 | 0.76 | 0.79 | 0.12 | 0.10 |
| *atpF* | 668 | 665 | 0.32 | 0.32 | 0.03 | 0.04 | 0.00 | 0.00 | 1.00 | 1.00 | 1.00 | 1.00 | 1.00 | 1.00 | 0.12 | 0.12 |
| *petB* | 763 | 763 | 0.33 | 0.33 | 0.02 | 0.02 | 0.00 | 0.00 | 1.00 | 1.00 | 1.00 | 1.00 | 1.00 | 1.00 | 0.10 | 0.10 |
| *petD* | 693 | 694 | 0.33 | 0.33 | 0.06 | 0.06 | 0.01 | 0.00 | 1.00 | 1.00 | 1.00 | 1.00 | 1.00 | 1.00 | 0.12 | 0.11 |
| *rpl2* | 667 | 667 | 0.38 | 0.38 | 0.02 | 0.02 | 0.00 | 0.00 | 1.00 | 1.00 | 1.00 | 1.00 | 1.00 | 1.00 | 0.08 | 0.08 |
| *rpoC1* | 755 | 712 | 0.36 | 0.36 | 0.08 | 0.05 | 0.00 | 0.00 | 1.00 | 1.00 | 1.00 | 1.00 | 1.00 | 1.00 | 0.18 | 0.13 |
| *trnA* | 827 | 827 | 0.51 | 0.51 | 0.01 | 0.01 | 0.00 | 0.00 | 1.00 | 1.00 | 1.00 | 1.00 | 1.00 | 1.00 | 0.05 | 0.05 |
| *trnL* | 440 | 429 | 0.35 | 0.36 | 0.05 | 0.04 | 0.01 | 0.01 | 1.00 | 1.00 | 1.00 | 1.00 | 1.00 | 1.00 | 0.11 | 0.11 |
| *trnV* | 574 | 574 | 0.39 | 0.39 | 0.02 | 0.02 | 0.00 | 0.00 | 1.00 | 1.00 | 1.00 | 1.00 | 1.00 | 1.00 | 0.11 | 0.11 |
| *ycf3 #1* | 740 | 740 | 0.33 | 0.33 | 0.05 | 0.05 | 0.00 | 0.00 | 1.00 | 1.00 | 1.00 | 1.00 | 1.00 | 1.00 | 0.10 | 0.10 |
| *ycf3 #2* | 702 | 682 | 0.37 | 0.38 | 0.03 | 0.02 | 0.01 | 0.01 | 1.00 | 1.00 | 1.00 | 1.00 | 1.00 | 1.00 | 0.14 | 0.14 |
| *rpl16* | 1,048 | 1,048 | 0.29 | 0.29 | 0.09 | 0.09 | 0.01 | 0.01 | 0.98 | 0.98 | 0.97 | 0.97 | 0.95 | 0.95 | 0.16 | 0.16 |
| *clpP #2* | 829 | 763 | 0.32 | 0.34 | 0.10 | 0.06 | 0.01 | 0.01 | 0.95 | 0.97 | 0.86 | 0.93 | 0.82 | 0.90 | 0.21 | 0.16 |
| *rps16* | 848 | 813 | 0.34 | 0.34 | 0.08 | 0.06 | 0.01 | 0.01 | 0.96 | 0.95 | 0.95 | 0.95 | 0.92 | 0.90 | 0.17 | 0.15 |
| *trnG* | 726 | 704 | 0.30 | 0.31 | 0.14 | 0.20 | 0.01 | 0.01 | 0.92 | 0.94 | 0.91 | 0.95 | 0.84 | 0.90 | 0.15 | 0.12 |
| *ndhA* | 1,061 | 1,016 | 0.31 | 0.32 | 0.07 | 0.04 | 0.01 | 0.00 | 0.90 | 0.96 | 0.84 | 0.91 | 0.75 | 0.87 | 0.13 | 0.12 |
| *trnK #2* | 312 | 304 | 0.28 | 0.29 | 0.18 | 0.16 | 0.02 | 0.02 | 0.92 | 0.89 | 0.91 | 0.91 | 0.84 | 0.81 | 0.23 | 0.17 |
| *clpP #1* | 633 | 611 | 0.32 | 0.33 | 0.05 | 0.04 | 0.01 | 0.00 | 0.94 | 0.92 | 0.88 | 0.80 | 0.82 | 0.74 | 0.13 | 0.12 |
| *trnK #1* | 763 | 759 | 0.32 | 0.33 | 0.08 | 0.08 | 0.02 | 0.01 | 0.81 | 0.86 | 0.68 | 0.76 | 0.55 | 0.66 | 0.16 | 0.15 |
| *ndhB* | 671 | 647 | 0.40 | 0.40 | 0.01 | 0.00 | 0.00 | 0.00 | 1.00 | 1.00 | n.a. | n.a. | n.a. | n.a. | 0.12 | 0.06 |
| *trnI* | 779 | 766 | 0.49 | 0.50 | 0.01 | 0.01 | 0.00 | 0.00 | 1.00 | 1.00 | n.a. | n.a. | n.a. | n.a. | 0.05 | 0.05 |

**Appendix S3**. Alignment metrics and homoplasy indices of the MSAs of the intergenic spacers before and after alignment adjustment. The columns are: alignment length, GC content, fraction of polymorphic sites, fraction of parsimony informative sites, consistency index, retention index, rescaled consistency index, maximum uncorrected p-distance. The table is ordered by the value of the rescaled consistency index, but the concatenated dataset of all markers is always on top. Fractions refer to the full alignment length.

| | Alignm. (bp) | | GC content | | Polymorph. sites | | Pars. inform. sites | | Consist. index | | Retent. index | | Rescaled index | | Uncorr. p-dist. | |
|---|---|---|---|---|---|---|---|---|---|---|---|---|---|---|---|---|
| | before | after | before | after | before | after | before | after | before | after | before | after | before | after | before | after |
| 103 IGS | 38,752 | 37,051 | 0.31 | 0.32 | 0.10 | 0.10 | 0.01 | 0.01 | 0.88 | 0.92 | 0.79 | 0.86 | 0.69 | 0.79 | 0.15 | 0.14 |
| *atpA–trnR-TCT* | 116 | 116 | 0.19 | 0.19 | 0.07 | 0.07 | 0.01 | 0.01 | 1.00 | 1.00 | 1.00 | 1.00 | 1.00 | 1.00 | 0.21 | 0.21 |
| *atpH–atpF* | 392 | 407 | 0.30 | 0.30 | 0.08 | 0.11 | 0.02 | 0.00 | 1.00 | 1.00 | 1.00 | 1.00 | 1.00 | 1.00 | 0.22 | 0.16 |
| *atpI–atpH* | 1,139 | 1,086 | 0.31 | 0.32 | 0.09 | 0.07 | 0.00 | 0.00 | 1.00 | 1.00 | 1.00 | 1.00 | 1.00 | 1.00 | 0.14 | 0.13 |
| *ndhD–ccsA* | 282 | 282 | 0.30 | 0.30 | 0.08 | 0.08 | 0.01 | 0.01 | 1.00 | 1.00 | 1.00 | 1.00 | 1.00 | 1.00 | 0.17 | 0.17 |
| *ndhE–psaC* | 247 | 247 | 0.26 | 0.26 | 0.05 | 0.09 | 0.00 | 0.00 | 1.00 | 1.00 | 1.00 | 1.00 | 1.00 | 1.00 | 0.20 | 0.18 |
| *ndhG–ndhE* | 228 | 228 | 0.28 | 0.28 | 0.07 | 0.07 | 0.00 | 0.00 | 1.00 | 1.00 | 1.00 | 1.00 | 1.00 | 1.00 | 0.19 | 0.19 |
| *petB–petD* | 195 | 195 | 0.28 | 0.28 | 0.05 | 0.05 | 0.01 | 0.01 | 1.00 | 1.00 | 1.00 | 1.00 | 1.00 | 1.00 | 0.19 | 0.19 |
| *petD–rpoA* | 202 | 203 | 0.31 | 0.31 | 0.13 | 0.11 | 0.01 | 0.01 | 1.00 | 1.00 | 1.00 | 1.00 | 1.00 | 1.00 | 0.21 | 0.13 |
| *petG–trnW-CCA* | 117 | 117 | 0.35 | 0.35 | 0.03 | 0.03 | 0.01 | 0.01 | 1.00 | 1.00 | 1.00 | 1.00 | 1.00 | 1.00 | 0.18 | 0.18 |
| *petL–petG* | 153 | 153 | 0.31 | 0.31 | 0.03 | 0.03 | 0.01 | 0.01 | 1.00 | 1.00 | 1.00 | 1.00 | 1.00 | 1.00 | 0.16 | 0.16 |
| *psaC–ndhD* | 115 | 114 | 0.39 | 0.39 | 0.09 | 0.13 | 0.02 | 0.01 | 1.00 | 1.00 | 1.00 | 1.00 | 1.00 | 1.00 | 0.23 | 0.13 |
| *psaI–ycf4* | 391 | 380 | 0.35 | 0.36 | 0.04 | 0.03 | 0.00 | 0.00 | 1.00 | 1.00 | 1.00 | 1.00 | 1.00 | 1.00 | 0.09 | 0.09 |
| *psbB–psbT* | 196 | 186 | 0.31 | 0.32 | 0.20 | 0.18 | 0.01 | 0.01 | 1.00 | 1.00 | 1.00 | 1.00 | 1.00 | 1.00 | 0.20 | 0.16 |
| *psbH–petB* | 124 | 124 | 0.35 | 0.35 | 0.05 | 0.05 | 0.03 | 0.03 | 1.00 | 1.00 | 1.00 | 1.00 | 1.00 | 1.00 | 0.22 | 0.22 |
| *psbI–trnS-GCT* | 142 | 146 | 0.25 | 0.25 | 0.17 | 0.16 | 0.01 | 0.01 | 1.00 | 1.00 | 1.00 | 1.00 | 1.00 | 1.00 | 0.27 | 0.21 |
| *psbK–psbI* | 453 | 453 | 0.29 | 0.29 | 0.15 | 0.15 | 0.00 | 0.00 | 1.00 | 1.00 | 1.00 | 1.00 | 1.00 | 1.00 | 0.16 | 0.16 |
| *psbZ–trnG-GCC* | 316 | 316 | 0.33 | 0.33 | 0.09 | 0.09 | 0.02 | 0.02 | 1.00 | 1.00 | 1.00 | 1.00 | 1.00 | 1.00 | 0.15 | 0.15 |
| *rbcL–accD* | 525 | 525 | 0.31 | 0.31 | 0.06 | 0.06 | 0.01 | 0.01 | 1.00 | 1.00 | 1.00 | 1.00 | 1.00 | 1.00 | 0.15 | 0.14 |
| *rpl33–rps18* | 188 | 188 | 0.29 | 0.29 | 0.01 | 0.01 | 0.01 | 0.01 | 1.00 | 1.00 | 1.00 | 1.00 | 1.00 | 1.00 | 0.07 | 0.07 |
| *rpoC1–rpoC2* | 101 | 101 | 0.37 | 0.37 | 0.03 | 0.10 | 0.03 | 0.01 | 1.00 | 1.00 | 1.00 | 1.00 | 1.00 | 1.00 | 0.18 | 0.10 |
| *rpoC2–rps2* | 229 | 214 | 0.38 | 0.40 | 0.08 | 0.04 | 0.00 | 0.01 | 1.00 | 1.00 | 1.00 | 1.00 | 1.00 | 1.00 | 0.18 | 0.14 |
| *rps12–ycf15* | 955 | 955 | 0.37 | 0.37 | 0.02 | 0.02 | 0.00 | 0.00 | 1.00 | 1.00 | 1.00 | 1.00 | 1.00 | 1.00 | 0.07 | 0.07 |
| *rps19–rpl2* | 54 | 54 | 0.30 | 0.30 | 0.04 | 0.04 | 0.02 | 0.02 | 1.00 | 1.00 | 1.00 | 1.00 | 1.00 | 1.00 | 0.14 | 0.14 |

Continued on next page

Appendix S3 – continued from previous page

| | Alignm. (bp) | | GC content | | Polymorph. sites | | Pars. inform. sites | | Consist. index | | Retent. index | | Rescaled index | | Uncorr. p-dist. | |
|---|---|---|---|---|---|---|---|---|---|---|---|---|---|---|---|---|
| | before | after | before | after | before | after | before | after | before | after | before | after | before | after | before | after |
| *rps2–atpI* | 221 | 221 | 0.26 | 0.26 | 0.02 | 0.02 | 0.01 | 0.01 | 1.00 | 1.00 | 1.00 | 1.00 | 1.00 | 1.00 | 0.14 | 0.14 |
| *rps4–trnT-TGT* | 372 | 376 | 0.22 | 0.22 | 0.31 | 0.33 | 0.01 | 0.00 | 1.00 | 1.00 | 1.00 | 1.00 | 1.00 | 1.00 | 0.22 | 0.22 |
| *rps7–ycf15* | 1,805 | 1,805 | 0.38 | 0.38 | 0.02 | 0.02 | 0.00 | 0.00 | 1.00 | 1.00 | 1.00 | 1.00 | 1.00 | 1.00 | 0.07 | 0.07 |
| *rrn16–trnI-GAT* | 294 | 294 | 0.50 | 0.51 | 0.06 | 0.07 | 0.01 | 0.01 | 1.00 | 1.00 | 1.00 | 1.00 | 1.00 | 1.00 | 0.21 | 0.10 |
| *rrn5–trnR-ACG* | 251 | 238 | 0.43 | 0.45 | 0.06 | 0.05 | 0.00 | 0.00 | 1.00 | 1.00 | 1.00 | 1.00 | 1.00 | 1.00 | 0.07 | 0.07 |
| *trnA-TGC–rrn23* | 152 | 152 | 0.42 | 0.42 | 0.03 | 0.03 | 0.02 | 0.02 | 1.00 | 1.00 | 1.00 | 1.00 | 1.00 | 1.00 | 0.14 | 0.14 |
| *trnF-GAA–ndhJ* | 784 | 787 | 0.27 | 0.27 | 0.14 | 0.15 | 0.01 | 0.01 | 1.00 | 1.00 | 1.00 | 1.00 | 1.00 | 1.00 | 0.16 | 0.12 |
| *trnG-GCC–trnM-CAT* | 201 | 201 | 0.28 | 0.28 | 0.10 | 0.10 | 0.01 | 0.01 | 1.00 | 1.00 | 1.00 | 1.00 | 1.00 | 1.00 | 0.16 | 0.16 |
| *trnL-TAA–trnF-GAA* | 350 | 350 | 0.33 | 0.33 | 0.05 | 0.05 | 0.01 | 0.01 | 1.00 | 1.00 | 1.00 | 1.00 | 1.00 | 1.00 | 0.18 | 0.18 |
| *trnM-CAT–atpE* | 207 | 246 | 0.30 | 0.30 | 0.11 | 0.23 | 0.05 | 0.02 | 1.00 | 1.00 | 1.00 | 1.00 | 1.00 | 1.00 | 0.29 | 0.24 |
| *trnM-CAT–rps14* | 150 | 150 | 0.35 | 0.35 | 0.11 | 0.19 | 0.04 | 0.04 | 1.00 | 1.00 | 1.00 | 1.00 | 1.00 | 1.00 | 0.22 | 0.22 |
| *trnN-GTT–ycf1* | 333 | 333 | 0.37 | 0.37 | 0.02 | 0.02 | 0.00 | 0.00 | 1.00 | 1.00 | 1.00 | 1.00 | 1.00 | 1.00 | 0.08 | 0.08 |
| *trnP-TGG–psaJ* | 346 | 346 | 0.28 | 0.28 | 0.13 | 0.13 | 0.00 | 0.00 | 1.00 | 1.00 | 1.00 | 1.00 | 1.00 | 1.00 | 0.15 | 0.15 |
| *trnR-ACG–trnN-GTT* | 472 | 472 | 0.45 | 0.45 | 0.05 | 0.05 | 0.00 | 0.00 | 1.00 | 1.00 | 1.00 | 1.00 | 1.00 | 1.00 | 0.10 | 0.10 |
| *trnS-GGA–rps4* | 316 | 294 | 0.35 | 0.37 | 0.13 | 0.08 | 0.02 | 0.01 | 1.00 | 1.00 | 1.00 | 1.00 | 1.00 | 1.00 | 0.19 | 0.12 |
| *trnV-GAC–rrn16* | 225 | 225 | 0.46 | 0.46 | 0.00 | 0.00 | 0.00 | 0.00 | 1.00 | 1.00 | 1.00 | 1.00 | 1.00 | 1.00 | 0.07 | 0.07 |
| *trnV-TAC–trnM-CAT* | 176 | 176 | 0.29 | 0.29 | 0.03 | 0.03 | 0.02 | 0.02 | 1.00 | 1.00 | 1.00 | 1.00 | 1.00 | 1.00 | 0.15 | 0.15 |
| *trnW-CCA–trnP-TGG* | 164 | 164 | 0.34 | 0.34 | 0.10 | 0.10 | 0.01 | 0.01 | 1.00 | 1.00 | 1.00 | 1.00 | 1.00 | 1.00 | 0.16 | 0.16 |
| *ycf15–trnV-GAC* | 680 | 680 | 0.42 | 0.42 | 0.01 | 0.01 | 0.00 | 0.00 | 1.00 | 1.00 | 1.00 | 1.00 | 1.00 | 1.00 | 0.07 | 0.07 |
| *ycf1–rps15* | 214 | 200 | 0.19 | 0.20 | 0.07 | 0.04 | 0.01 | 0.01 | 1.00 | 1.00 | 1.00 | 1.00 | 1.00 | 1.00 | 0.15 | 0.16 |
| *ycf2–trnL-CAA* | 427 | 427 | 0.45 | 0.45 | 0.02 | 0.02 | 0.01 | 0.01 | 1.00 | 1.00 | 1.00 | 1.00 | 1.00 | 1.00 | 0.11 | 0.11 |
| *ccsA–trnL-TAG* | 120 | 107 | 0.23 | 0.25 | 0.12 | 0.07 | 0.05 | 0.01 | 0.95 | 1.00 | 0.97 | 1.00 | 0.92 | 1.00 | 0.28 | 0.22 |
| *ndhF–trnN-GTT* | 946 | 48 | 0.36 | 0.23 | 0.05 | 0.69 | 0.01 | 0.04 | 0.94 | 1.00 | 0.96 | 1.00 | 0.90 | 1.00 | 0.09 | n.a. |
| *psaJ–rpl33* | 437 | 424 | 0.32 | 0.31 | 0.07 | 0.06 | 0.01 | 0.01 | 0.96 | 1.00 | 0.94 | 1.00 | 0.90 | 1.00 | 0.19 | 0.15 |
| *psbM–trnD-GTC* | 649 | 672 | 0.33 | 0.33 | 0.05 | 0.08 | 0.01 | 0.01 | 0.97 | 1.00 | 0.93 | 1.00 | 0.90 | 1.00 | 0.17 | 0.14 |
| *ycf4–cemA* | 378 | 348 | 0.31 | 0.31 | 0.09 | 0.07 | 0.01 | 0.01 | 0.94 | 1.00 | 0.91 | 1.00 | 0.86 | 1.00 | 0.17 | 0.13 |
| *psbC–trnS-TGA* | 245 | 214 | 0.34 | 0.38 | 0.20 | 0.12 | 0.02 | 0.01 | 0.90 | 1.00 | 0.79 | 1.00 | 0.71 | 1.00 | 0.28 | 0.17 |
| *trnC-GCA–petN* | 836 | 789 | 0.31 | 0.32 | 0.16 | 0.12 | 0.00 | 0.00 | 0.97 | 1.00 | 0.60 | 1.00 | 0.58 | 1.00 | 0.20 | 0.19 |
| *atpB–rbcL* | 762 | 707 | 0.29 | 0.30 | 0.08 | 0.04 | 0.01 | 0.00 | 0.87 | 1.00 | 0.64 | 1.00 | 0.56 | 1.00 | 0.18 | 0.13 |

Continued on next page

Appendix S3 – continued from previous page

| | Alignm. (bp) | | GC content | | Polymorph. sites | | Pars. inform. sites | | Consist. index | | Retent. index | | Rescaled index | | Uncorr. p-dist. | |
|---|---|---|---|---|---|---|---|---|---|---|---|---|---|---|---|---|
| | before | after | before | after | before | after | before | after | before | after | before | after | before | after | before | after |
| *accD–psaI* | 696 | 708 | 0.23 | 0.23 | 0.15 | 0.17 | 0.03 | 0.02 | 0.97 | 0.98 | 0.95 | 0.97 | 0.92 | 0.95 | 0.20 | 0.17 |
| *trnE-TTC–rpoB* | 802 | 785 | 0.31 | 0.32 | 0.17 | 0.15 | 0.01 | 0.01 | 0.96 | 0.98 | 0.93 | 0.96 | 0.89 | 0.94 | 0.18 | 0.17 |
| *ycf3–trnS-GGA* | 891 | 871 | 0.29 | 0.30 | 0.11 | 0.09 | 0.01 | 0.01 | 0.91 | 0.97 | 0.83 | 0.94 | 0.76 | 0.92 | 0.17 | 0.15 |
| *trnS-GCT–trnC-GCA* | 764 | 763 | 0.29 | 0.29 | 0.20 | 0.20 | 0.02 | 0.08 | 0.92 | 0.95 | 0.90 | 0.95 | 0.83 | 0.90 | 0.19 | 0.18 |
| *clpP–psbB* | 466 | 456 | 0.30 | 0.31 | 0.11 | 0.10 | 0.01 | 0.01 | 0.95 | 0.95 | 0.93 | 0.93 | 0.89 | 0.89 | 0.17 | 0.17 |
| *petA–psbJ* | 764 | 741 | 0.31 | 0.32 | 0.09 | 0.08 | 0.01 | 0.01 | 0.94 | 0.94 | 0.94 | 0.94 | 0.88 | 0.88 | 0.17 | 0.17 |
| *rpl32–ndhF* | 1,098 | 1,047 | 0.22 | 0.23 | 0.19 | 0.18 | 0.04 | 0.03 | 0.93 | 0.93 | 0.94 | 0.94 | 0.88 | 0.88 | 0.21 | 0.18 |
| *rps16–trnQ-TTG* | 984 | 970 | 0.28 | 0.28 | 0.11 | 0.10 | 0.02 | 0.02 | 0.91 | 0.94 | 0.90 | 0.94 | 0.82 | 0.88 | 0.15 | 0.15 |
| *trnT-TGT–trnL-TAA* | 635 | 572 | 0.26 | 0.27 | 0.19 | 0.12 | 0.02 | 0.01 | 0.88 | 0.95 | 0.81 | 0.92 | 0.71 | 0.88 | 0.18 | 0.13 |
| *rps18–rpl20* | 344 | 321 | 0.30 | 0.32 | 0.28 | 0.25 | 0.02 | 0.02 | 0.96 | 0.95 | 0.94 | 0.91 | 0.90 | 0.87 | 0.23 | 0.20 |
| *trnT-GGT–psbD* | 1,283 | 1,275 | 0.31 | 0.30 | 0.20 | 0.20 | 0.01 | 0.01 | 0.95 | 0.94 | 0.91 | 0.91 | 0.87 | 0.86 | 0.14 | 0.13 |
| *ndhC–trnV-TAC* | 765 | 771 | 0.31 | 0.31 | 0.07 | 0.07 | 0.02 | 0.01 | 0.87 | 0.95 | 0.73 | 0.91 | 0.63 | 0.86 | 0.19 | 0.17 |
| *psaA–ycf3* | 734 | 710 | 0.30 | 0.31 | 0.14 | 0.13 | 0.01 | 0.01 | 0.86 | 0.97 | 0.73 | 0.88 | 0.63 | 0.85 | 0.16 | 0.15 |
| *trnK-TTT–rps16* | 877 | 850 | 0.25 | 0.26 | 0.17 | 0.20 | 0.03 | 0.03 | 0.94 | 0.92 | 0.95 | 0.92 | 0.89 | 0.84 | 0.20 | 0.16 |
| *psbA–trnK-TTT* | 295 | 255 | 0.30 | 0.30 | 0.12 | 0.12 | 0.02 | 0.01 | 0.90 | 0.93 | 0.85 | 0.90 | 0.76 | 0.84 | 0.23 | 0.22 |
| *petN–psbM* | 509 | 499 | 0.28 | 0.28 | 0.08 | 0.12 | 0.02 | 0.02 | 0.84 | 0.91 | 0.83 | 0.93 | 0.70 | 0.84 | 0.20 | 0.15 |
| *cemA–petA* | 244 | 245 | 0.32 | 0.32 | 0.05 | 0.04 | 0.03 | 0.01 | 0.93 | 0.90 | 0.96 | 0.91 | 0.89 | 0.82 | 0.19 | 0.16 |
| *ndhI–ndhG* | 376 | 377 | 0.22 | 0.22 | 0.14 | 0.17 | 0.02 | 0.02 | 0.95 | 0.93 | 0.88 | 0.88 | 0.83 | 0.81 | 0.24 | 0.21 |
| *trnG-TCC–trnT-GGT* | 188 | 188 | 0.31 | 0.31 | 0.20 | 0.20 | 0.02 | 0.02 | 0.92 | 0.92 | 0.88 | 0.88 | 0.81 | 0.81 | 0.18 | 0.18 |
| *trnL-TAG–rpl32* | 818 | 775 | 0.24 | 0.25 | 0.13 | 0.10 | 0.02 | 0.02 | 0.86 | 0.93 | 0.71 | 0.86 | 0.61 | 0.80 | 0.20 | 0.20 |
| *trnQ-TTG–psbK* | 368 | 368 | 0.26 | 0.26 | 0.09 | 0.10 | 0.01 | 0.01 | 0.96 | 0.94 | 0.83 | 0.83 | 0.80 | 0.78 | 0.18 | 0.12 |
| *trnY-GTA–trnE-TTC* | 203 | 203 | 0.33 | 0.33 | 0.04 | 0.04 | 0.02 | 0.02 | 0.86 | 0.86 | 0.91 | 0.91 | 0.78 | 0.78 | 0.14 | 0.14 |
| *psbE–petL* | 1,268 | 1,187 | 0.29 | 0.30 | 0.12 | 0.08 | 0.01 | 0.01 | 0.86 | 0.82 | 0.84 | 0.82 | 0.72 | 0.68 | 0.13 | 0.13 |
| *rps14–psaB* | 133 | 133 | 0.33 | 0.33 | 0.04 | 0.04 | 0.03 | 0.03 | 0.83 | 0.83 | 0.80 | 0.80 | 0.67 | 0.67 | 0.17 | 0.17 |
| *rpl16–rps3* | 145 | 145 | 0.23 | 0.23 | 0.09 | 0.09 | 0.03 | 0.03 | 0.81 | 0.81 | 0.70 | 0.70 | 0.57 | 0.57 | 0.20 | 0.20 |
| *rpl14–rpl16* | 120 | 98 | 0.25 | 0.28 | 0.24 | 0.22 | 0.01 | 0.00 | 1.00 | 1.00 | 1.00 | n.a. | 1.00 | n.a. | 0.35 | 0.23 |
| *rpl36–infA* | 115 | 129 | 0.32 | 0.33 | 0.08 | 0.15 | 0.04 | 0.00 | 1.00 | 1.00 | 1.00 | n.a. | 1.00 | n.a. | 0.21 | 0.13 |
| *rps8–rpl14* | 211 | 170 | 0.23 | 0.26 | 0.19 | 0.06 | 0.04 | 0.00 | 1.00 | 1.00 | 1.00 | n.a. | 1.00 | n.a. | 0.24 | 0.16 |
| *trnR-TCT–trnG-TCC* | 221 | 185 | 0.20 | 0.22 | 0.14 | 0.06 | 0.01 | 0.00 | 1.00 | 1.00 | 1.00 | n.a. | 1.00 | n.a. | 0.24 | 0.17 |

Continued on next page

Appendix S3 – continued from previous page

| | Alignm. (bp) | | GC content | | Polymorph. sites | | Pars. inform. sites | | Consist. index | | Retent. index | | Rescaled index | | Uncorr. p-dist. | |
|---|---|---|---|---|---|---|---|---|---|---|---|---|---|---|---|---|
| | before | after | before | after | before | after | before | after | before | after | before | after | before | after | before | after |
| *trnS-TGA–psbZ* | 340 | 329 | 0.36 | 0.36 | 0.17 | 0.15 | 0.00 | 0.00 | 1.00 | 1.00 | 1.00 | n.a. | 1.00 | n.a. | 0.12 | 0.12 |
| *atpF–atpA* | 78 | 62 | 0.21 | 0.24 | 0.15 | 0.08 | 0.00 | 0.00 | 1.00 | 1.00 | n.a. | n.a. | n.a. | n.a. | 0.23 | 0.25 |
| *infA–rps8* | 121 | 121 | 0.36 | 0.36 | 0.02 | 0.02 | 0.00 | 0.00 | 1.00 | 1.00 | n.a. | n.a. | n.a. | n.a. | 0.13 | 0.13 |
| *ndhA–ndhI* | 78 | 78 | 0.26 | 0.26 | 0.06 | 0.06 | 0.00 | 0.00 | 1.00 | 1.00 | n.a. | n.a. | n.a. | n.a. | 0.23 | 0.23 |
| *ndhB–rps7* | 289 | 289 | 0.33 | 0.33 | 0.02 | 0.02 | 0.00 | 0.00 | 1.00 | 1.00 | n.a. | n.a. | n.a. | n.a. | 0.12 | 0.12 |
| *ndhJ–ndhK* | 105 | 105 | 0.31 | 0.31 | 0.01 | 0.01 | 0.00 | 0.00 | 1.00 | 1.00 | n.a. | n.a. | n.a. | n.a. | 0.10 | 0.10 |
| *psbJ–psbL* | 148 | 148 | 0.39 | 0.39 | 0.05 | 0.05 | 0.00 | 0.00 | 1.00 | 1.00 | n.a. | n.a. | n.a. | n.a. | 0.14 | 0.14 |
| *rpl22–rps19* | 105 | 77 | 0.17 | 0.22 | 0.16 | 0.10 | 0.00 | 0.00 | 1.00 | 1.00 | n.a. | n.a. | n.a. | n.a. | 0.17 | 0.12 |
| *rpl23–trnI-CAT* | 165 | 165 | 0.33 | 0.33 | 0.01 | 0.01 | 0.00 | 0.00 | 1.00 | 1.00 | n.a. | n.a. | n.a. | n.a. | 0.08 | 0.08 |
| *rpoA–rps11* | 89 | 89 | 0.23 | 0.23 | 0.11 | 0.11 | 0.00 | 0.00 | 1.00 | 1.00 | n.a. | n.a. | n.a. | n.a. | 0.11 | 0.11 |
| *rps11–rpl36* | 105 | 105 | 0.32 | 0.32 | 0.02 | 0.02 | 0.00 | 0.00 | 1.00 | 1.00 | n.a. | n.a. | n.a. | n.a. | 0.14 | 0.14 |
| *rps15–ndhH* | 97 | 97 | 0.31 | 0.31 | 0.07 | 0.07 | 0.00 | 0.00 | 1.00 | 1.00 | n.a. | n.a. | n.a. | n.a. | 0.10 | 0.10 |
| *rrn23–rrn45* | 99 | 99 | 0.56 | 0.56 | 0.01 | 0.01 | 0.00 | 0.00 | 1.00 | 1.00 | n.a. | n.a. | n.a. | n.a. | 0.10 | 0.10 |
| *rrn45–rrn5* | 251 | 251 | 0.45 | 0.45 | 0.02 | 0.02 | 0.00 | 0.00 | 1.00 | 1.00 | n.a. | n.a. | n.a. | n.a. | 0.06 | 0.06 |
| *trnI-CAT–ycf2* | 111 | 111 | 0.32 | 0.32 | 0.01 | 0.01 | 0.00 | 0.00 | 1.00 | 1.00 | n.a. | n.a. | n.a. | n.a. | 0.10 | 0.10 |
| *trnL-CAA–ndhB* | 576 | 587 | 0.36 | 0.36 | 0.03 | 0.05 | 0.00 | 0.00 | 1.00 | 1.00 | n.a. | n.a. | n.a. | n.a. | 0.14 | 0.09 |
| *psaB–psaA* | 25 | 25 | 0.44 | 0.44 | 0.00 | 0.00 | 0.00 | 0.00 | n.a. | n.a. | n.a. | n.a. | n.a. | n.a. | 0.00 | 0.00 |
| *psbL–psbF* | 22 | 22 | 0.36 | 0.36 | 0.00 | 0.00 | 0.00 | 0.00 | n.a. | n.a. | n.a. | n.a. | n.a. | n.a. | 0.00 | 0.00 |
| *psbN–psbH* | 84 | 84 | 0.27 | 0.27 | 0.00 | 0.00 | 0.00 | 0.00 | n.a. | n.a. | n.a. | n.a. | n.a. | n.a. | 0.00 | 0.00 |
| *rpl2–rpl23* | 18 | 18 | 0.22 | 0.22 | 0.00 | 0.00 | 0.00 | 0.00 | n.a. | n.a. | n.a. | n.a. | n.a. | n.a. | 0.00 | 0.00 |
| *trnD-GTC–trnY-GTA* | 92 | 92 | 0.36 | 0.36 | 0.00 | 0.00 | 0.00 | 0.00 | n.a. | n.a. | n.a. | n.a. | n.a. | n.a. | 0.00 | 0.00 |
| *trnI-GAT–trnA-TGC* | 64 | 64 | 0.55 | 0.55 | 0.00 | 0.00 | 0.00 | 0.00 | n.a. | n.a. | n.a. | n.a. | n.a. | n.a. | 0.00 | 0.00 |

**Appendix S4**. Results of phylogenetic tree inference via ML on the concatenation of all MSAs of the coding regions, the concatenation of all MSAs of the introns, the concatenation of all MSAs of the intergenic spacers (i) before and (ii) after alignment adjustment. The trees displayed represent the trees with the highest likelihood score (a) without and (b) with the coding of indels. The inferred relationships are visualized as cladograms with statistical node support (left) and corresponding phylograms with exact branch lengths (right). Bootstrap support values greater than 50% are given above the branches of each cladogram. All trees were rooted with using *Ligularia fischeri* as outgroup.

### (i) Before alignment adjustment
#### (i–a) without indels

**CDS**

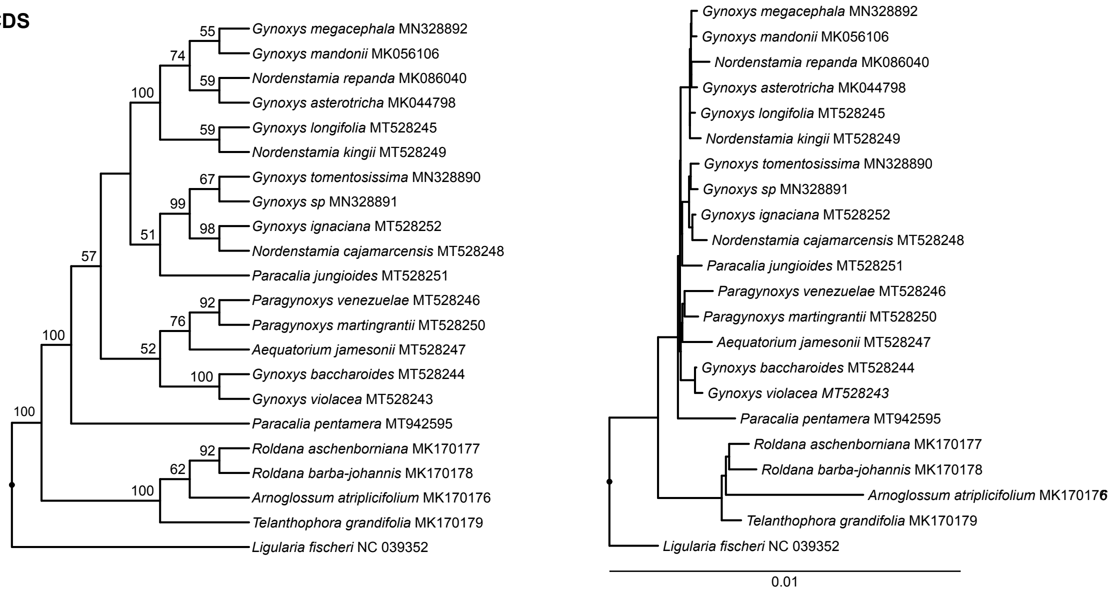


**INT**

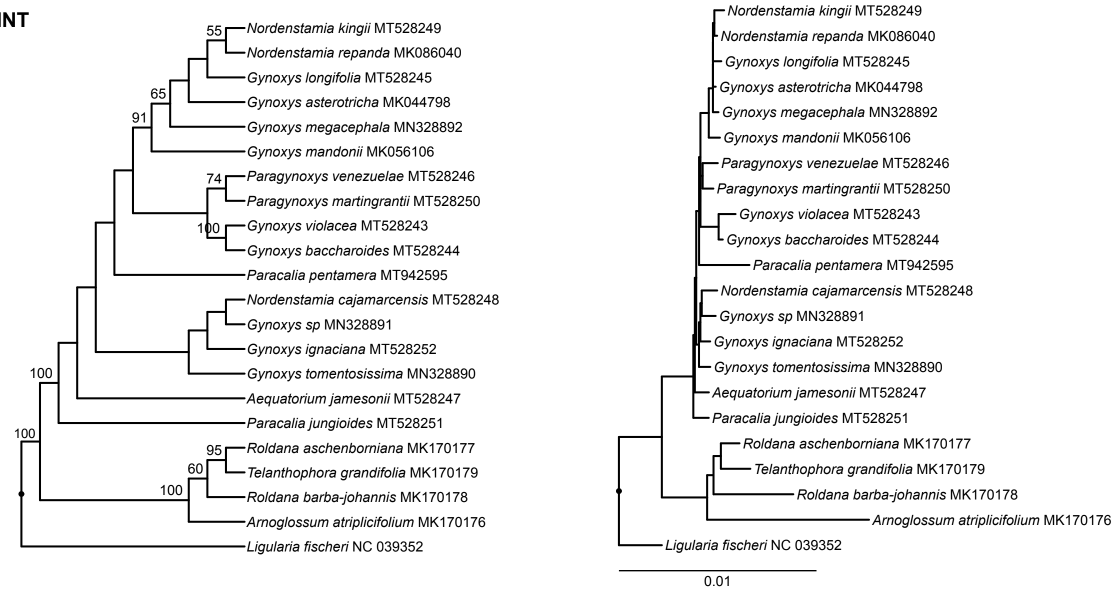


**IGS**

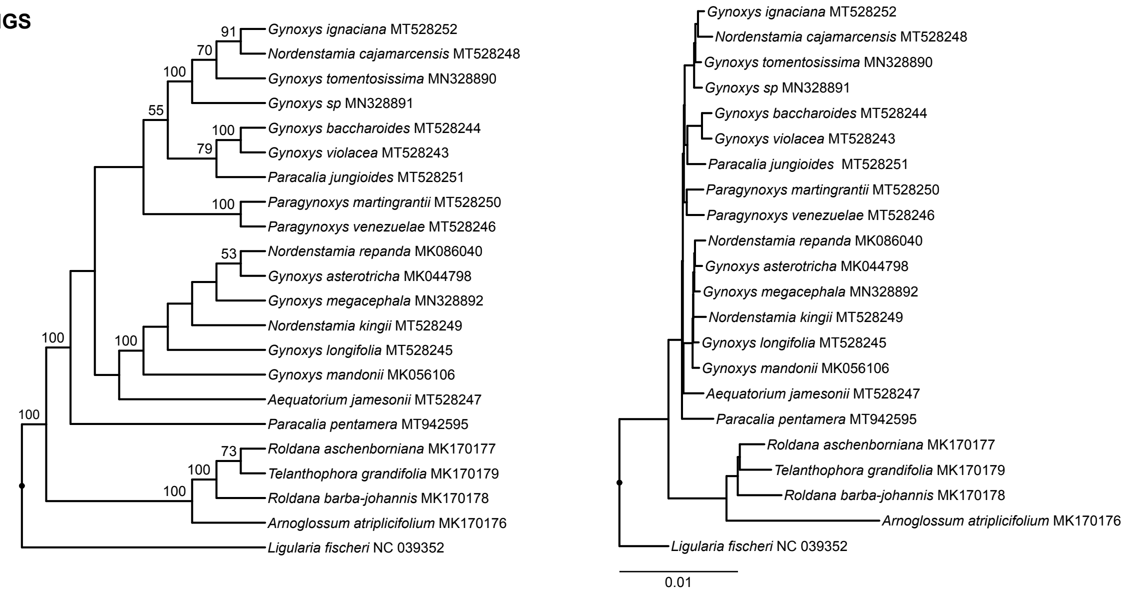

#### (i) Before alignment adjustment
##### (i–b) with indels

**CDS**

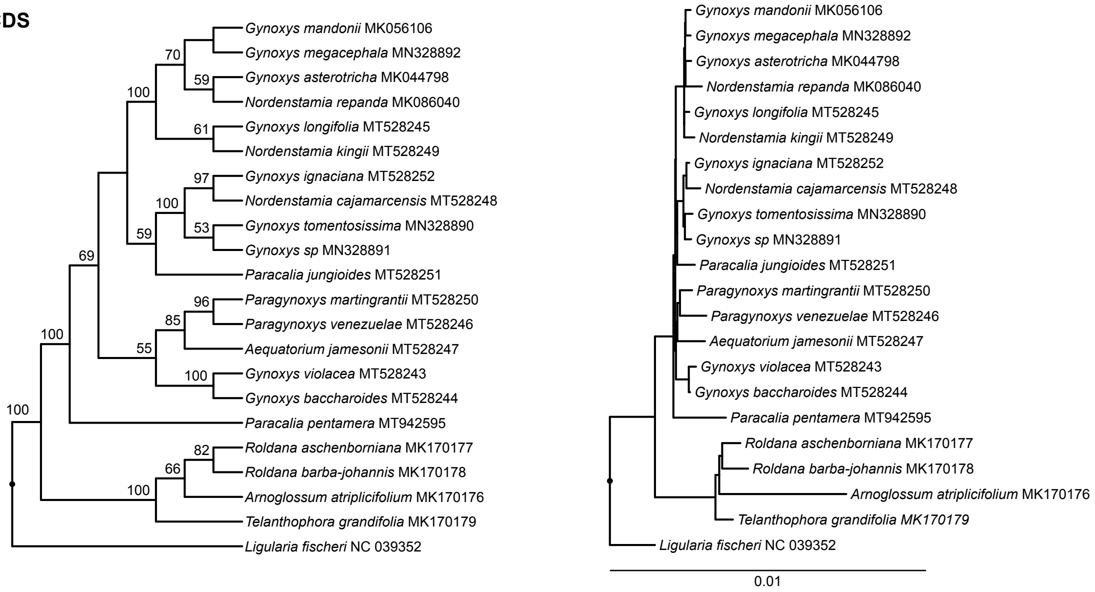


**INT**

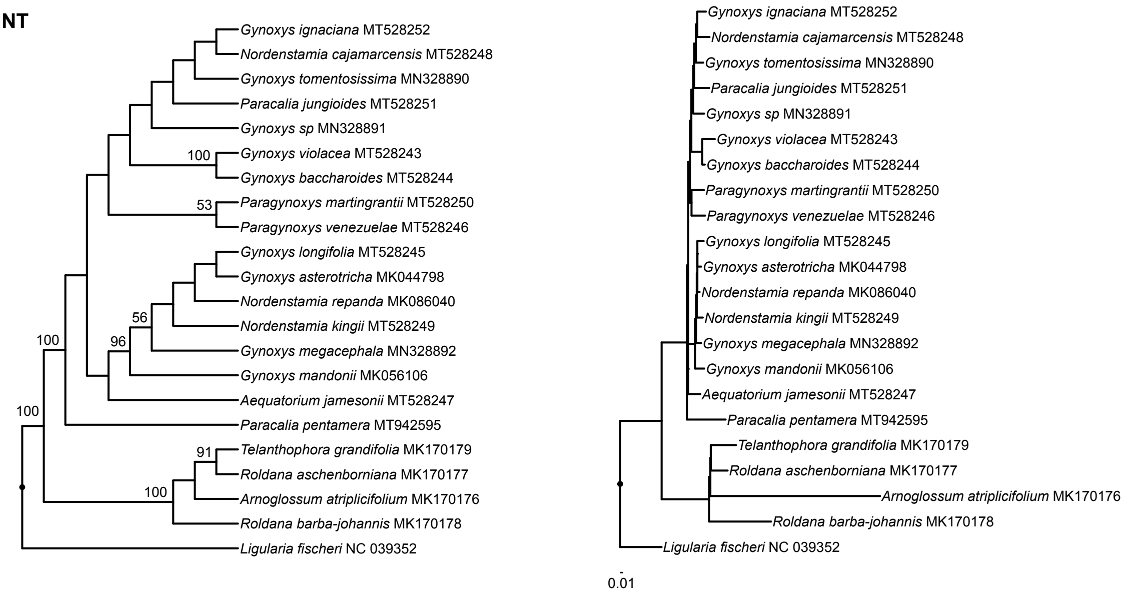


**IGS**

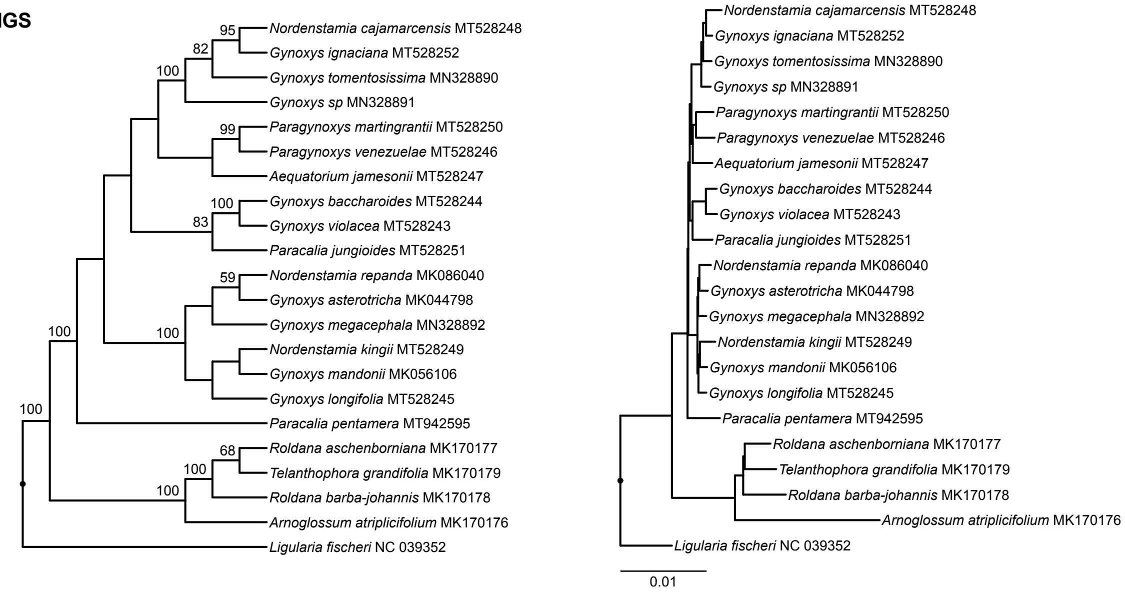

#### (ii) After alignment adjustment
##### (ii–a) without indels

**CDS**

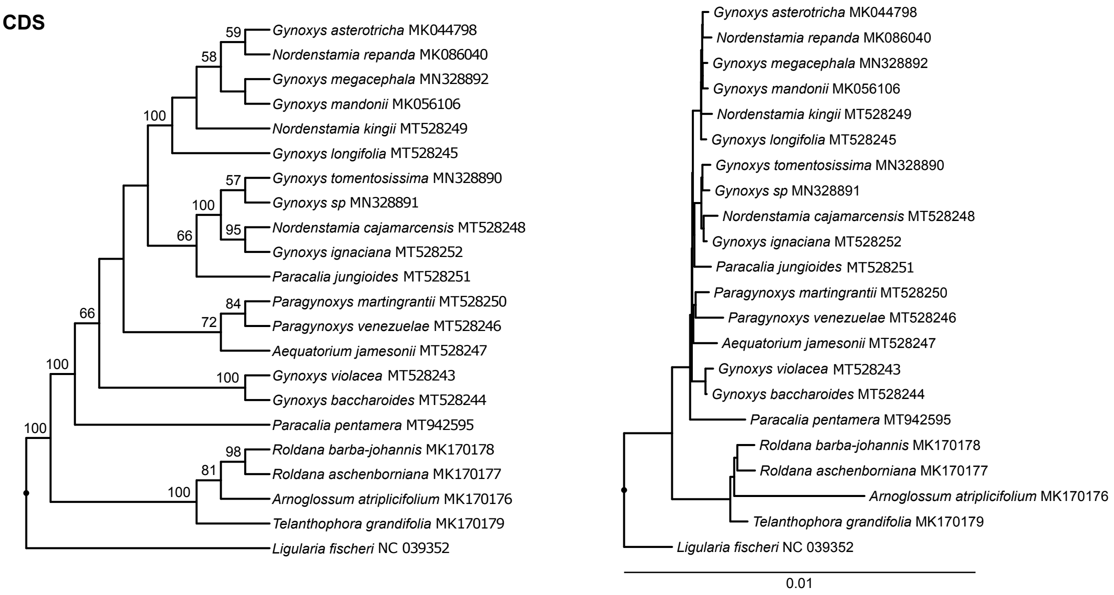


**INT**

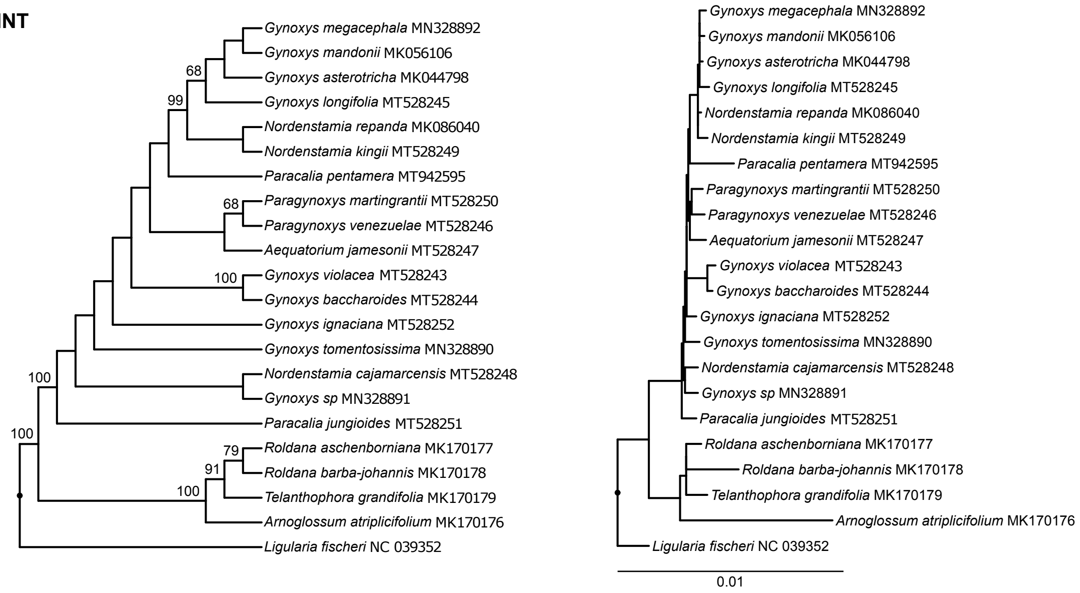


**IGS**

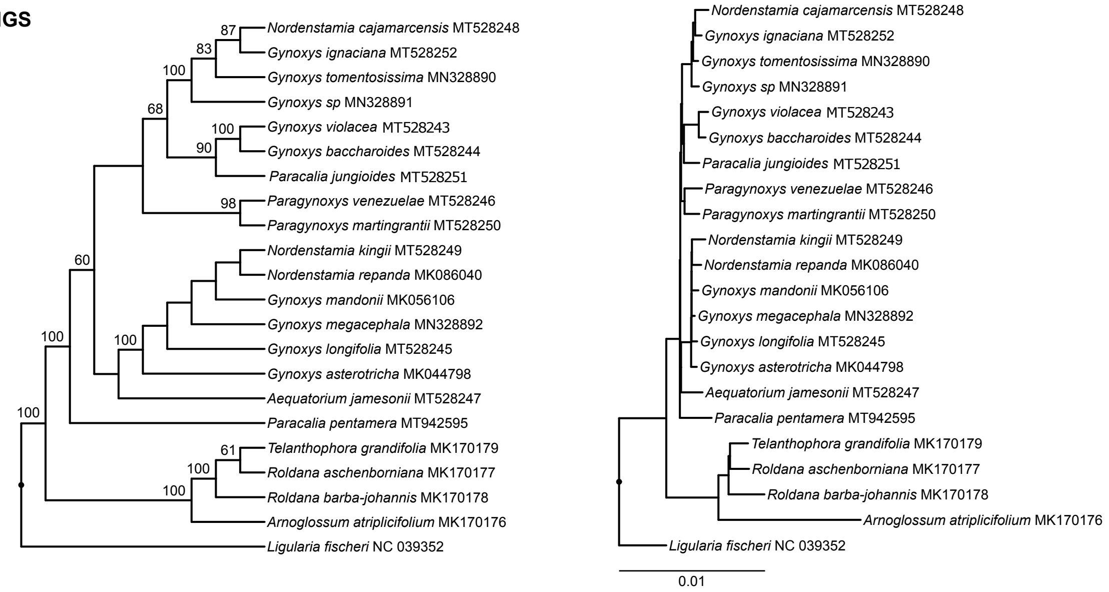

### (ii) After alignment adjustment
#### (ii–b) with indels

**CDS**

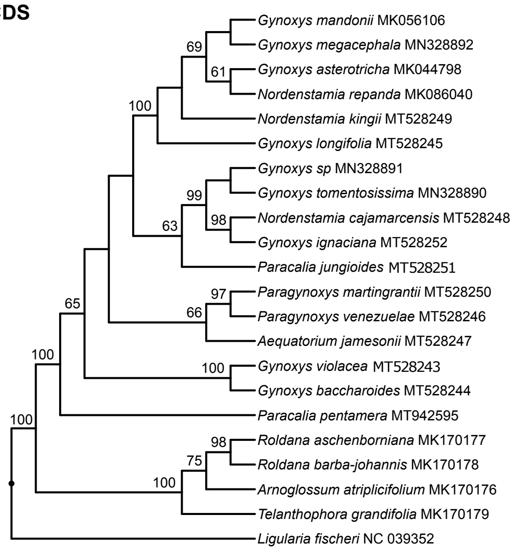


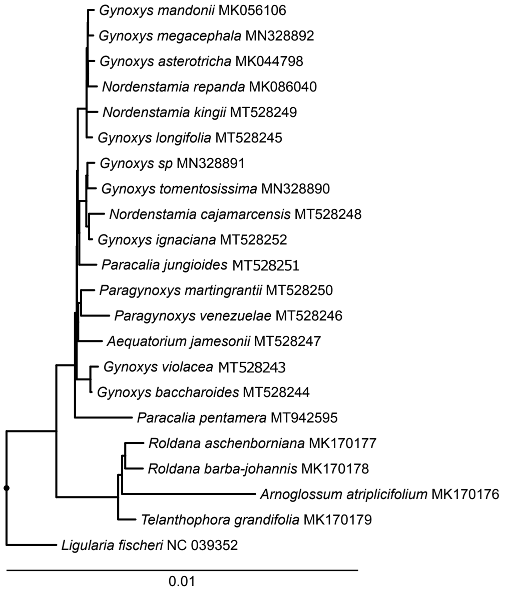


**INT**

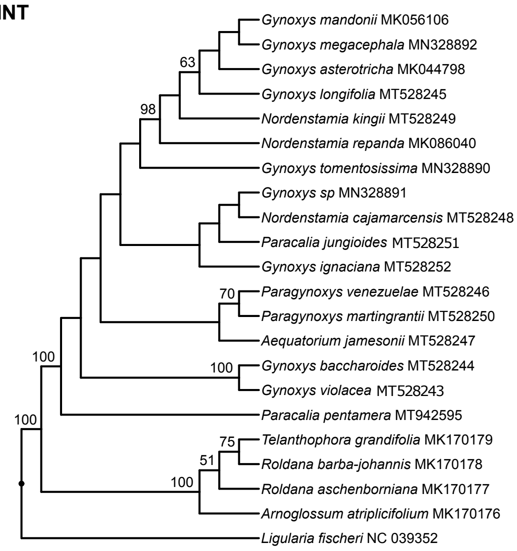


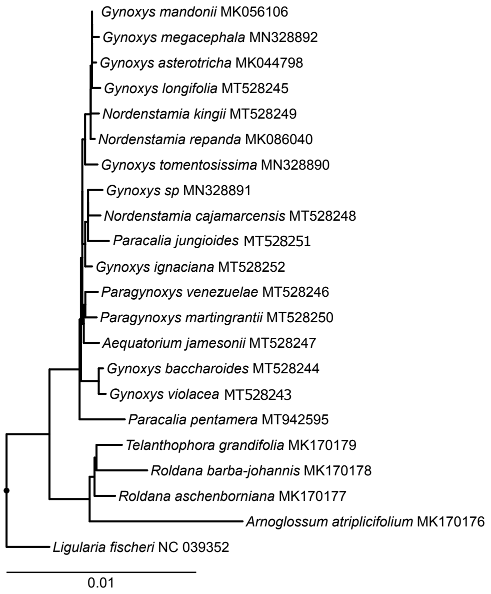


**IGS**

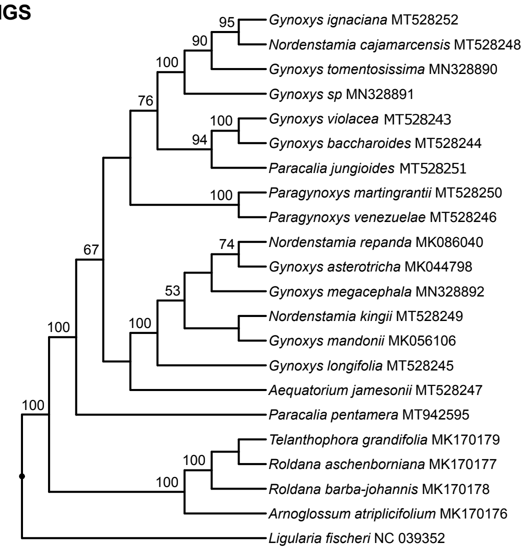


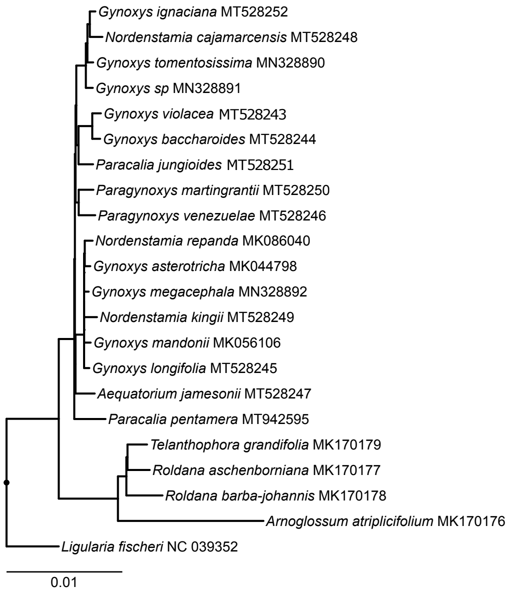

**Appendix S5**. Results of the phylogenetic tree inference via ML on the concatenation of all three plastid partitions before alignment adjustment. The trees displayed represent the trees with the highest likelihood score (a) without and (b) with the coding of indels. Visualization and rooting are as in Appendix S4.

#### (i) Before alignment adjustment

##### (i–a) without indels

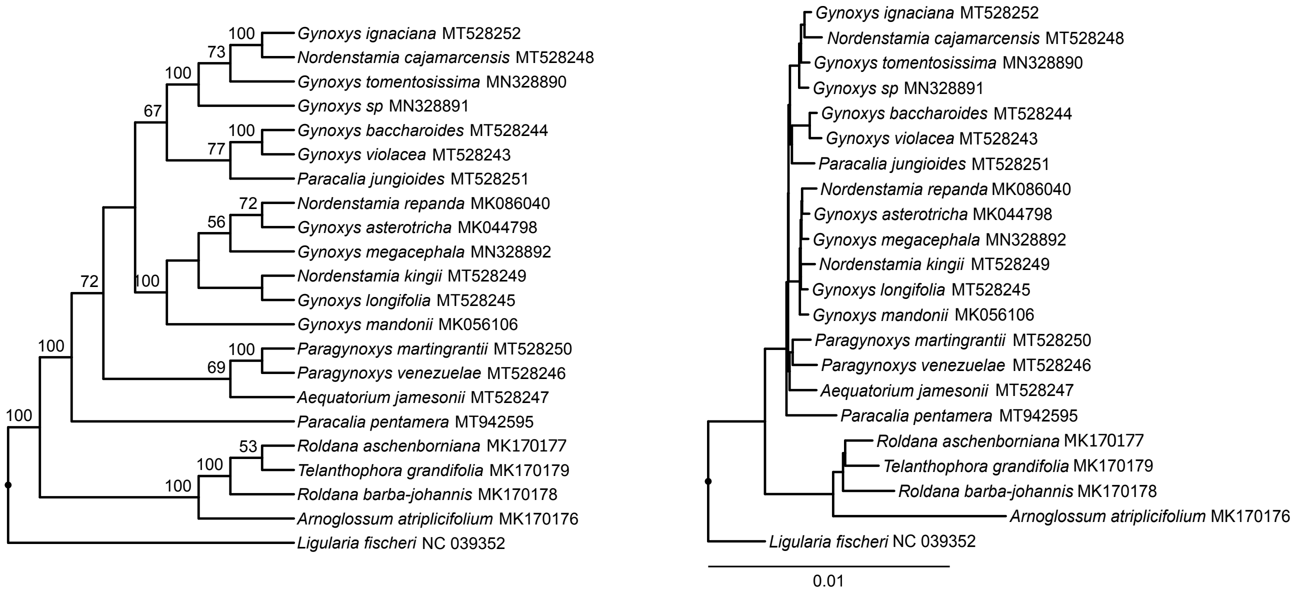


##### (i–b) with indels

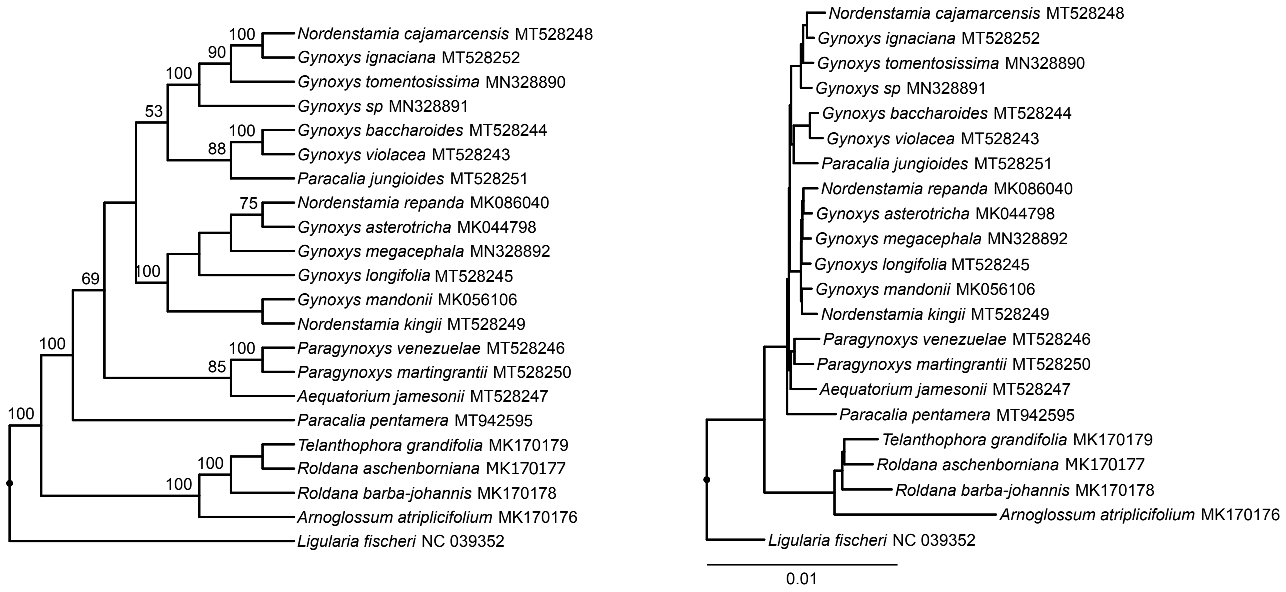

**Appendix S6**. Results of phylogenetic tree inference via BI on the concatenation of all MSAs of the coding regions, the concatenation of all MSAs of the introns, the concatenation of all MSAs of the intergenic spacers (i) before and (ii) after alignment adjustment. The trees displayed represent the 50% majority-rule consensus tree of the posterior tree distribution, visualized as cladograms with posterior probability values greater than 0.5 (left) and corresponding phylograms with exact branch lengths (right). All trees were rooted with using *Ligularia fischeri* as outgroup.

### (i) Before alignment adjustment

#### (i–a) without indels

**CDS**

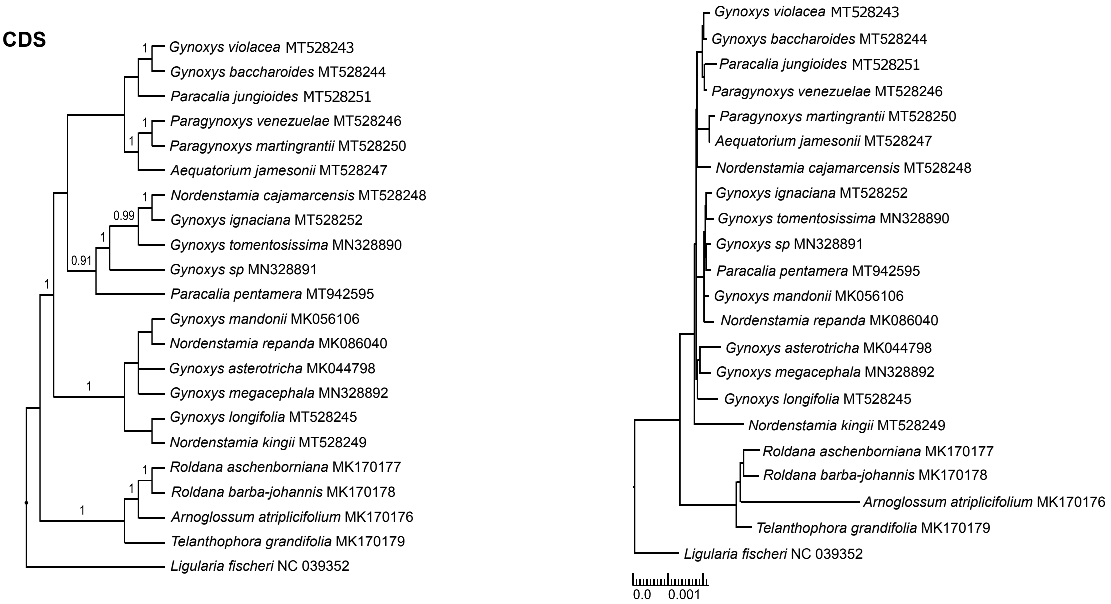


**INT**

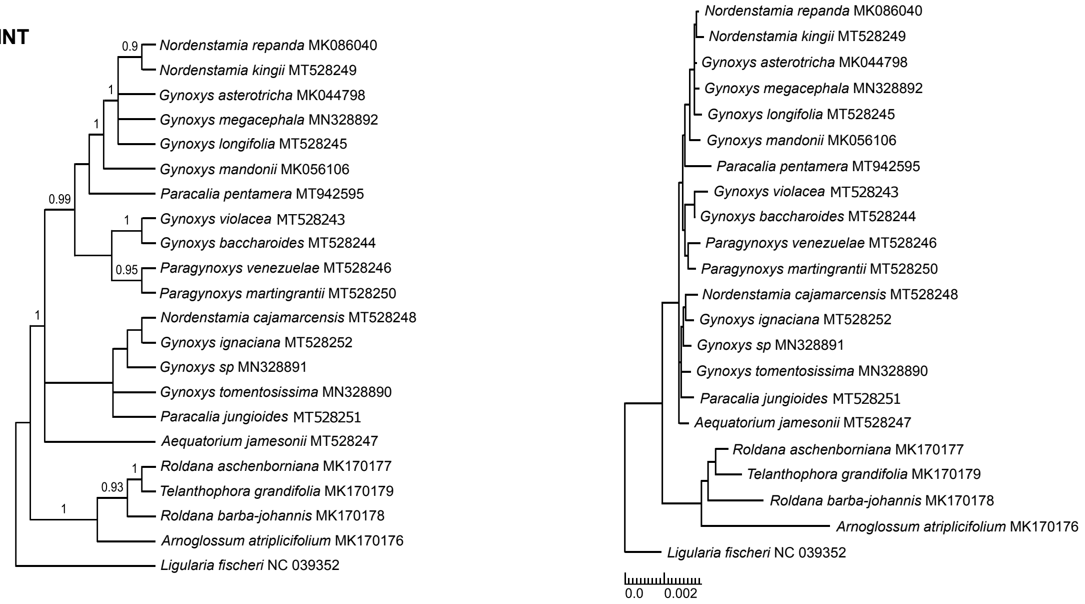


**IGS**

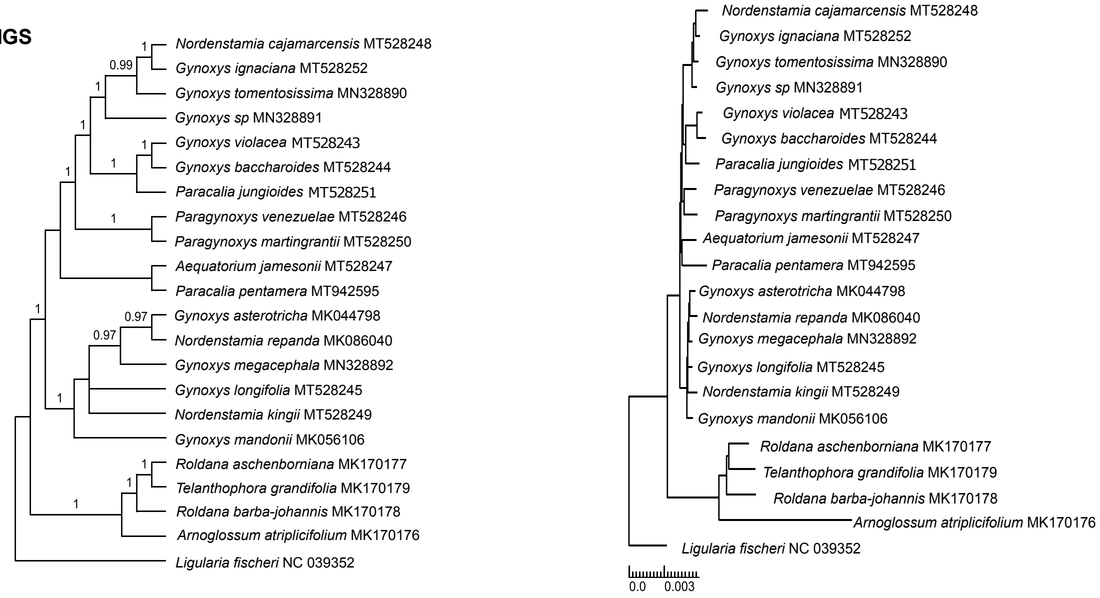

#### (i) Before alignment adjustment
##### (i–b) with indels

**CDS**

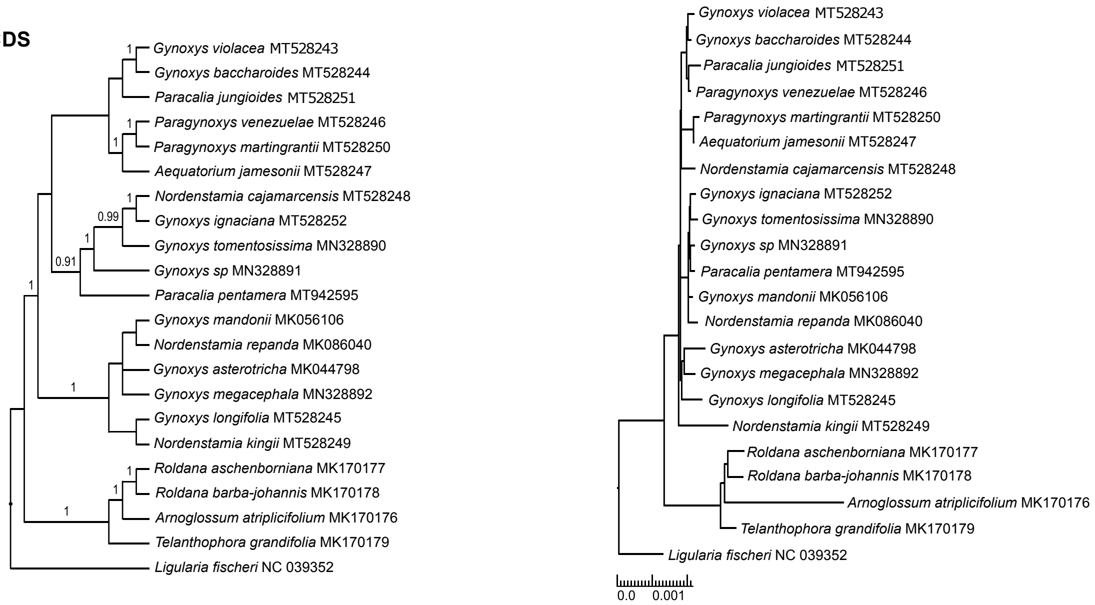


**INT**

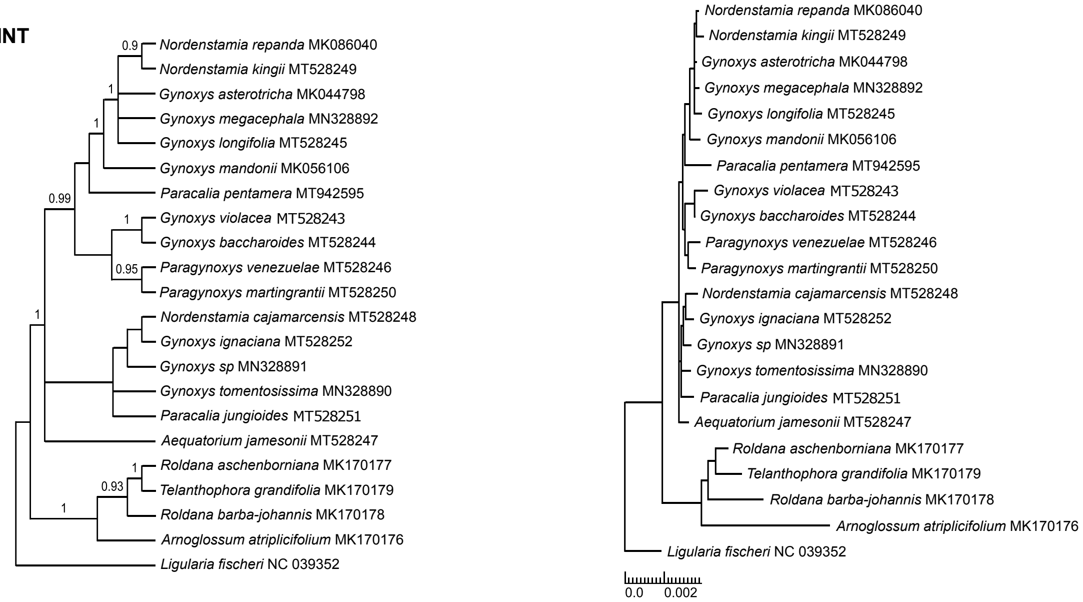


**IGS**

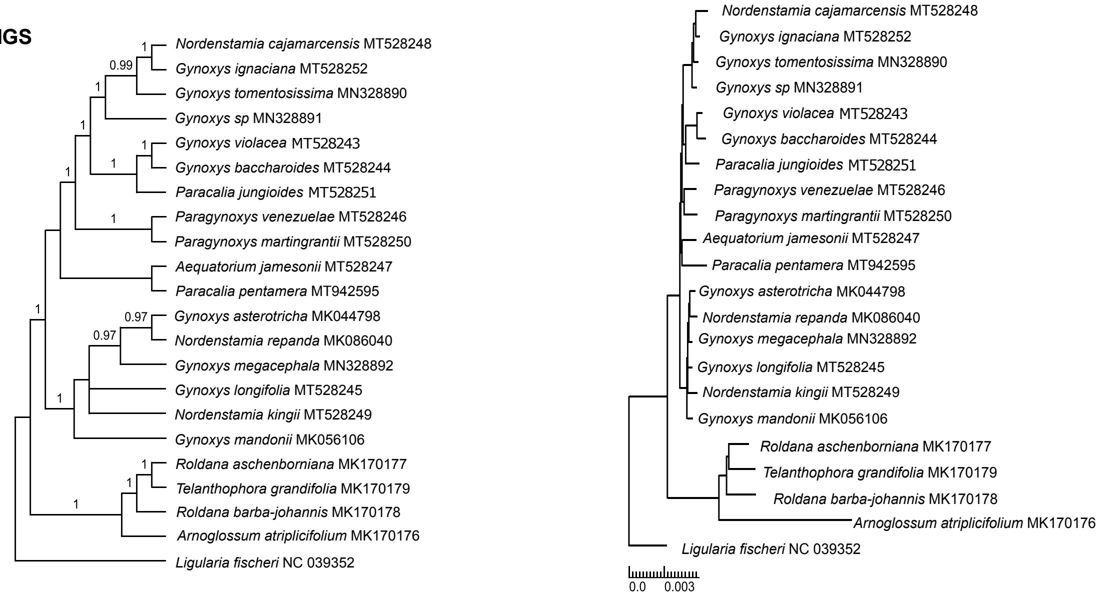

#### (ii) After alignment adjustment
##### (ii–a) without indels

**CDS**

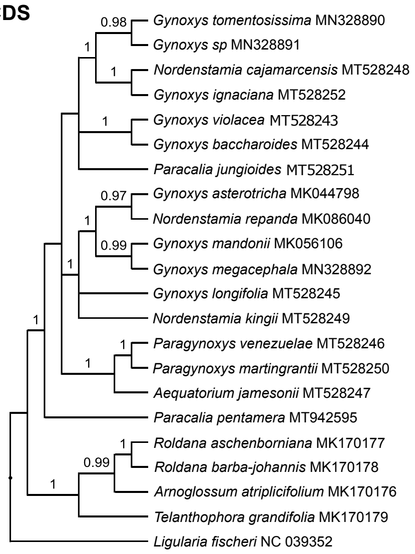


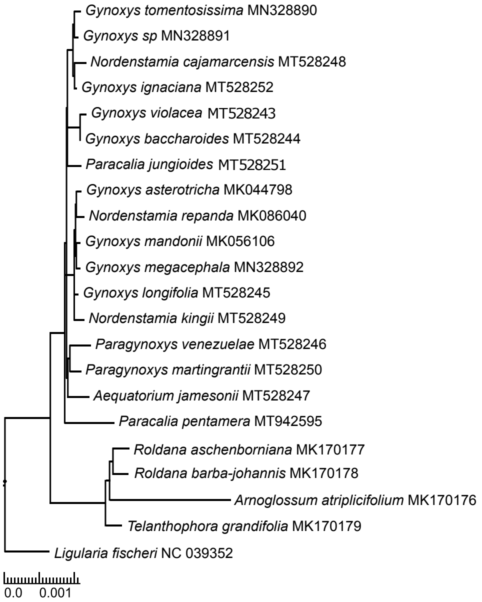


**INT**

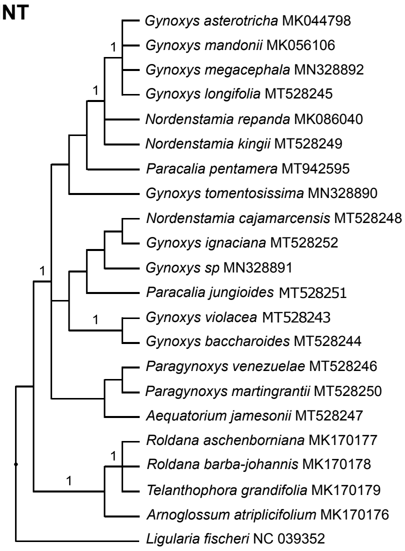


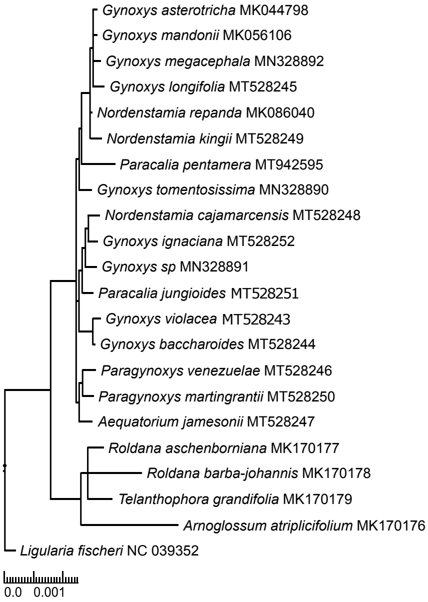


**IGS**

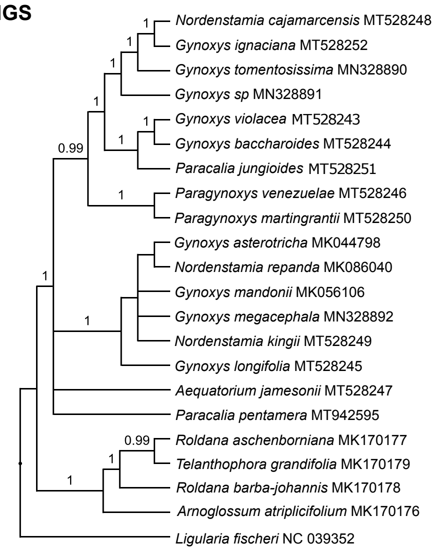


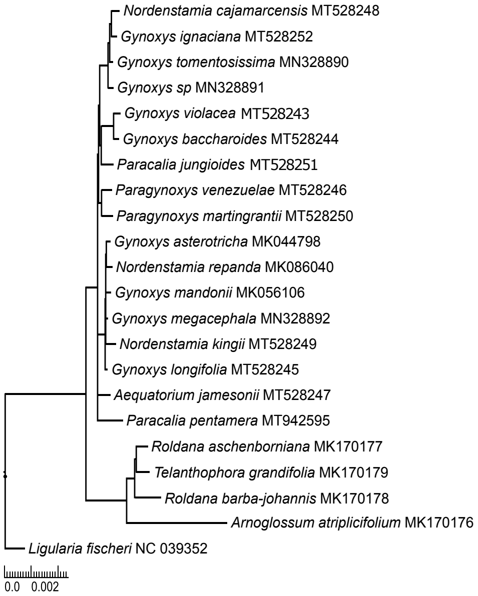

#### (ii) After alignment adjustment

##### (ii–b) with indels

**CDS**

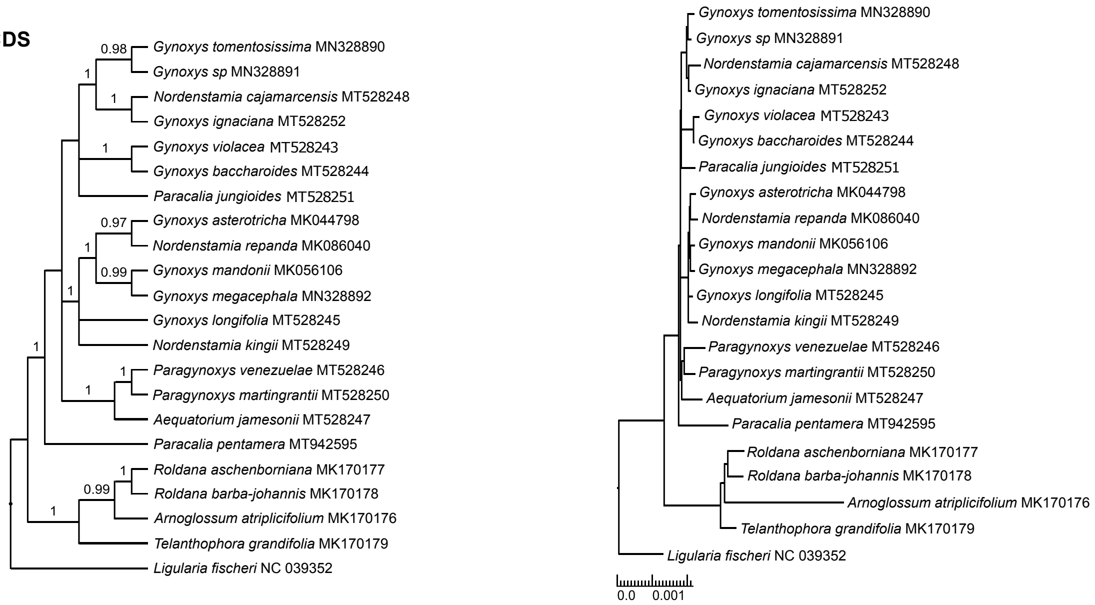


**INT**

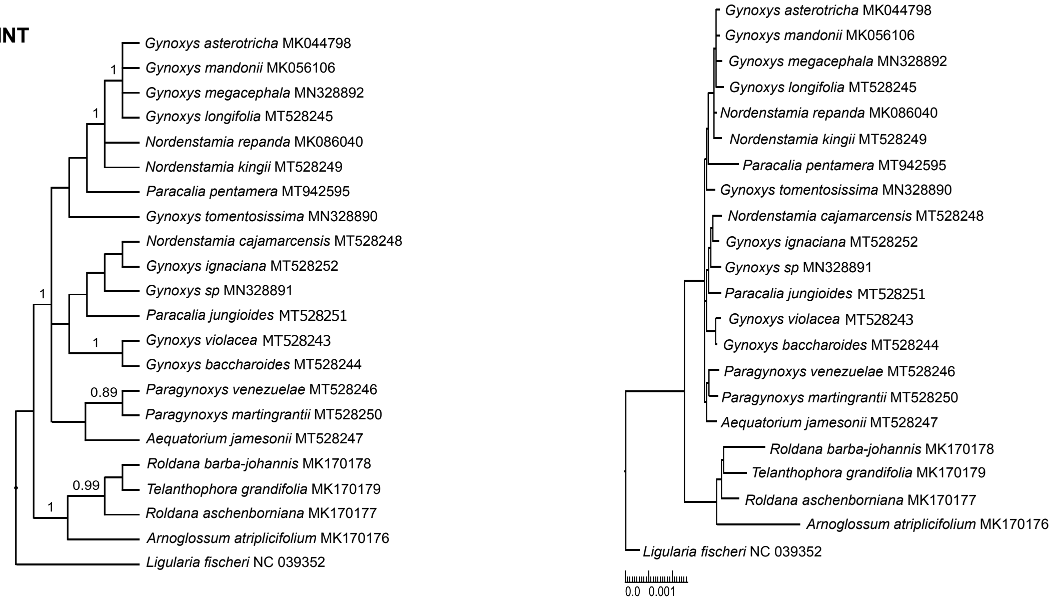


**IGS**

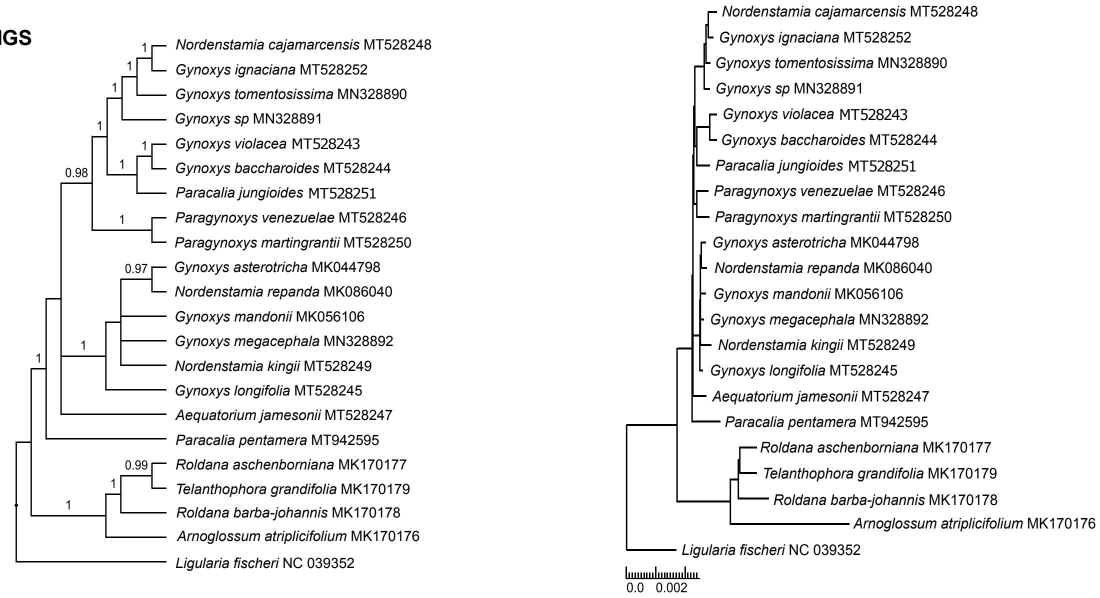

**Appendix S7**. Results of the phylogenetic tree inference via BI on the concatenation of all three plastid partitions before alignment adjustment. The trees displayed represent the 50% majority-rule consensus tree of the posterior tree distribution (a) without and (b) with the coding of indels. Visualization and rooting are as in Appendix S6.

#### (i) Before alignment adjustment

##### (i–a) without indels

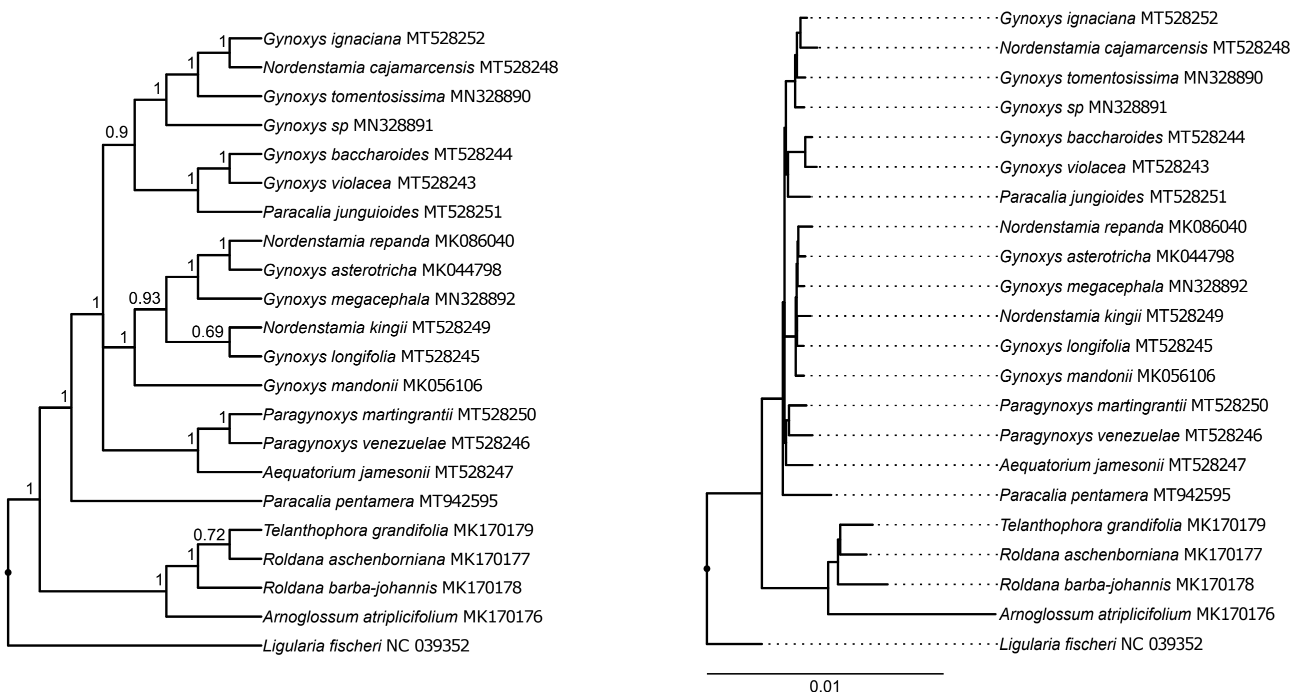


##### (i–b) with indels

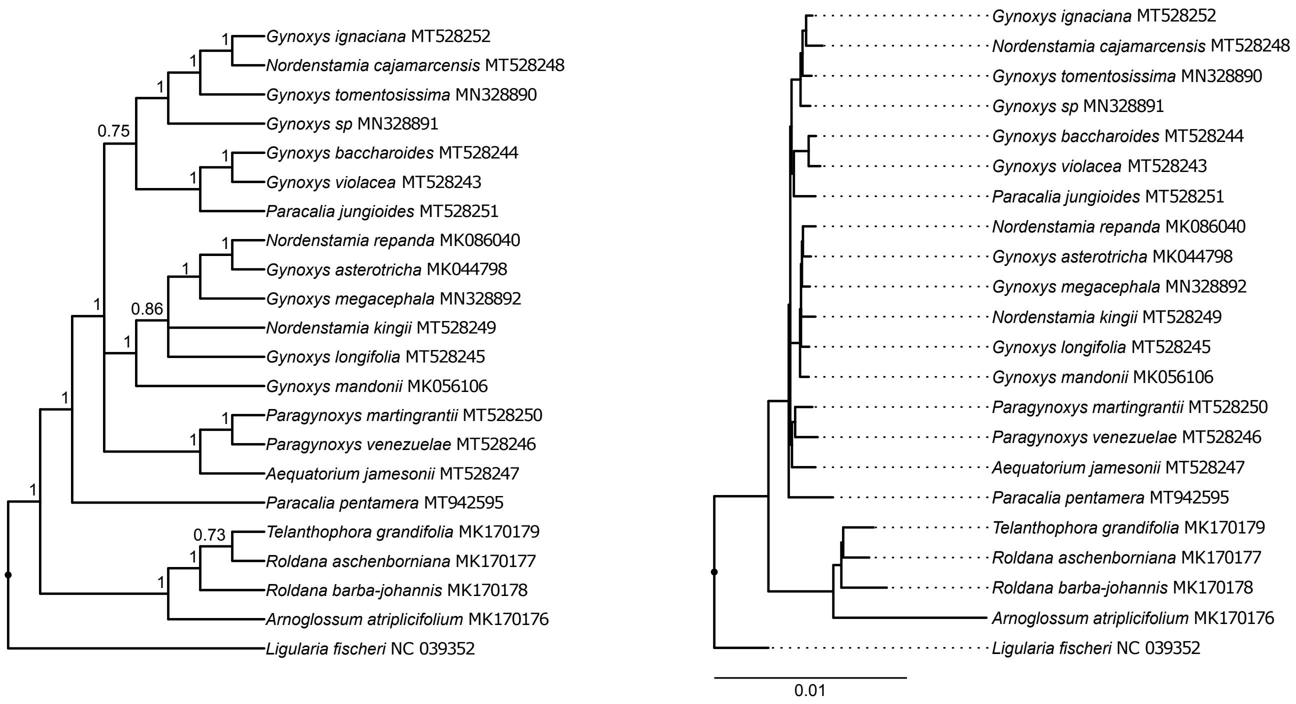
